# Exploring the adaptability and robustness of the central carbon metabolism of *Mycoplasma pneumoniae*

**DOI:** 10.1101/2022.08.08.503180

**Authors:** Niels A. Zondervan, Eva Yus, Daniel C. Sévin, Sira Martinez, Carolina Gallo, Peter J. Schaap, Maria Lluch-Senar, Luis Serrano, Vitor A. P. Martins dos Santos, Maria Suarez-Diez

## Abstract

In this study we explored the adaptability and robustness of glycolysis and pyruvate metabolism of *Mycoplasma pneumoniae* (*MPN*). We used a dual approach, we analysed metabolomics data collected for a large number of OE and KO mutants and perturbation samples. Furthermore, we trained a dynamic model of central carbon metabolism and tested the model’s capacity to predict these mutants and perturbation samples as well as identify key controlling factors in central carbon metabolism. Our analysis of metabolite data as well as our model analysis indicate *MPN* metabolism is inherently robust against perturbations due to its network structure. Two key control hubs of central carbon metabolism were identified.

## Introduction

*Mycoplasma* are gram positive bacteria adapted to an obligatory parasitic lifestyle able to infect a broad range of hosts^1^. It is estimated by the CDC that 2 million infections with *M. pneumoniae* (*MPN*) occur in the US alone on a yearly basis^2^. These infections lead to conditions ranging from mild to severe respiratory illness including life threatening conditions such as auto-immune diseases^3^. Therefore, it imperative to improve our understanding of *MPN*.

*MPN* has been established as a model organism for systems biology and large dataset collections are available informing on its genome, transcriptome^4^, proteome^5^, metabolome^6^, and transcripcional adaptations^7^. The metabolism and energetic expenditure of *MPN* have been thoroughly studied by combining a genome-scale constraint based model of metabolism with detailed experimental characterizations^8^. Neither energy production nor uptake of protein building blocks appear to limit growth of *MPN* only protein synthesis was found to be growth limiting^6,9^. Maintenance requirements are high for *MPN*, so most of the energy is devoted to maintenance instead of growth^8^.

MPN like other Mycoplasma’s has adopted to its pathogenic lifestyle leading to a severely reduced genome. Despite their small genome size and limited number of enzymes and relatively low number of regulators^10^, Mycoplasma’s are still able to adapt to a large number of conditions and still genes can be removed, as they have been seen not to be essential^11^. Many essential genes are however only present in some Mycoplasma species, suggesting alternatives to a minimal genome exist^12^. Since there are only few regulatory elements in the genome^10^ we hypothesized that a lot of the adaptability of *MPN* to adapt to changing growth conditions must be due to network structure and to allosteric control of its metabolism.

In this study we investigated the metabolism of *MPN* with a focus on central carbon metabolism and its allosteric control. Dynamic models were successfully used to investigate regulation and adaptation of central carbon metabolism to changing environmental conditions in other organism^13–20^.

Therefore, in this study we will combine analysis of metabolomics a large number of samples taken from varying environmental conditions, OE and KO mutants with a dynamic model of glycolysis and pyruvate metabolism, to identify key controlling metabolites and enzymes in central carbon metabolism. We tested single or combined addition of 1) an ATPase reaction, 2) O_2_ inhibition of Lactate Dehydrogenase and 3) NAD regeneration by NoxE using O_2_ to this model as potential mechanisms for *MPN’s* adaptability to various conditions.

We trained and tested the model’s ability to predict a wide range of environmental conditions, single and double overexpression mutants as well as mutants with single gene deletions. The model was able to predict these mutants with reasonable accuracy. In a recent study, local sensitivity analysis on a dynamic model of *E. coli* central carbon metabolism identified robustness as one of properties of central carbon metabolism of *E. coli* ^16^. This robustness is a system property resulting from the many feed-forward and feed-backward interactions in metabolism, such as allosteric control of glucose uptake as well as lactate and acetate metabolism. Another study in *E. coli* revealed that only three metabolites (FBP, F1P and cAMP) account for about 70% of the expression variability of central carbon metabolism enzymes through control of two transcription factors^21^. Similarly, our model predicts the central carbon metabolism of *MPN* to be inherently robust to changing conditions and identifies two main hubs of metabolic control. Clustering of samples of FBA OE and LDH KO mutants corroborate assumed allosteric control of LDH FBP. Additionally, the analysis of metabolomics data of *MPN* indicated that glycolipid metabolism might be linked to the high energy metabolites needed for growth of *MPN*. Our findings are in agreement with recent findings were some key lipids were identified to be needed for *MPN* growth on serum free medium^22^.

## Materials and methods

### Bacterial strains and culture conditions

*M. pneumoniae* strain M129 (passage 33-34) was grown in modified Hayflick medium and transformed by electroporation with the pMT85 transposon as previously described ^9^. Briefly, cells were split 1:10, and washed twice with 10 mL and collected in 300 μl Electroporation buffer (8 mM Hepes·HCl, 272 mM sucrose, pH 7.4) three days later. Cells (50 μl) were electroporated with 5 μg plasmid in 1 mm gapped cuvettes at 1.25 kV, 100 Ω, 25 μF (Gene Pulser Xcel Electroporator, Bio-Rad). Cells were recovered in Hayflick for 2 h at 37 °C, diluted 1:5 in Hayflick with 200 μg mL^-1^ gentamycin, selected for three days and then maintained with 80 μg mL^-1^ gentamycin. The cell lines used are detailed in **Error! Reference source not found.**.

### Transposon insertion mutants obtained by haystack mutagenesis

For the isolation of *M. pneumoniae* mutants, we used a collection of strains carrying insertions of transposon Tn4001^23^. The presence of the desired mutant was assayed by PCR using one primer that hybridizes to the transposon (directed outwards), and a second primer specific for the gene of interest. Mass spectroscopy is used to verify absence of the corresponding protein.

### Over Expression mutant construction

Genes to be overexpressed where cloned in the transposon Tn4001^24^ control of the promoter of the EF-tu gene ^24^.

### Growth curves

To obtain equal amounts of each sample, initial inocula for the growth curves were quantified. Briefly, cells were grown for 3 days in 25-cm^2^ flasks, collected in 1 mL medium and 100 μl was used for quantification using the BCA (bicinchoninic acid) protein assay kit (Pierce, see below). Same amounts of total protein (1 μg) were aliquoted per well in a 96-multiwell plate in duplicates. Two hundred μl of Hayflick medium was added per well and the cells were incubated in a Tecan Infinite plate reader at 37°C. The “growth index” (absorbance 430/560 nm, settle time at 300 msec and number of flashes equal to 25) was obtained every hour for 5 days as published^9^. To quantify growth, we determined two slopes of the growth curve. The first one is based on the time interval from 10 to 30 h (“early slope”) and the second one on the whole growth curve (“late”). The early slope was determined by considering the maximum median of the slope between two time points (eq. 1) separated by three time measurements over successive periods of 30 time points. The late slope was determined by considering the maximum median value of the slope between two time points separated by four time measurements (eq. 2) over successive periods of 30 time points.

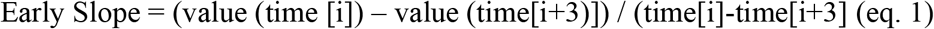

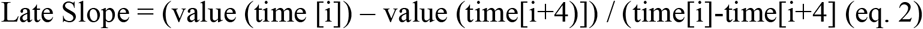

The early slope is more representative of growth, while the late slope reflects the metabolic activity.

On the other hand, biomass was quantified at 48 h (early stationary phase) by inoculating a twin 96-well plates, in the same conditions as above. After incubation for two days at 37°C under, medium was sucked out, cells were carefully washed twice with 200 μl PBS and lysed with 100 μl lysis buffer (10 mM Tris·HCl, 6 mM MgCl_2_, 1 mM EDTA, 100 mM NaCl, 0.1% Tx-100, pH 8, and 1× Protease Inhibitor Cocktail, Roche) at 4°C. In the same first 96-well plate, cell lysates were kept on ice and extracted protein was quantified by BCA Protein Assay Kit (Pierce, see below).

The protein concentrations at 48 h and early slope are more representatives of growth, while the late slope and the value of A430/560 at midpoint reflect the metabolic activity. These four parameters of growth and metabolism were analysed for each batch of experiments. Outliers (larger than quartile 3, Q3) by at least 1.5 times the interquartile range (IQR), or smaller than Q1 by at least 1.5 times the IQR) were removed to calculate the mean and the standard deviation of each of the parameters for each batch. Values larger or smaller than the mean by at least 2 times the standard deviation of each parameter were considered to determine fast- and slow-growing/metabolizing clones, respectively.

#### Strain cultivation and growth conditions of mutant and perturbation samples

A 300 cm^2^ flask was inoculated 1:10 with the lab stock and 100 mL of Hayflick and grown for 3-4 days at 37C. Then, medium was removed, and cells were scrapped and resuspended in 12 mL medium. From this inoculum, 75 cm^2^ flasks were seeded with 1 mL of inoculum in 20 mL of Hayflick. After 6 hours of incubation (i.e. when cells reached stationary growth phase) the cells were treated as follows, before the standard extraction protocol:

■ Glucose starvation: remove medium and add new Hayflick medium without glucose. Incubate sample for 5 h at 37C. Long incubation time is required to deplete glucose from Hayflick medium.
■ Amino acid starvation: take half of the medium, add 200 mg of DL-serine hydroxamate (10 mg/mL), mix and add again to the cells; incubate cells with for 15 min at 37C.
■ Fe^2+^ depletion: Add directly to the flask the iron chelator 2,2’-Bipyridine at a final concentration of 3 mM, incubate for 30 min at 37° C.
■ Oxidative stress: Add directly to the flask H_2_O_2_, to 0.5%, incubate 15 min at 37C.
■ Glycerol: Add directly to the flask glycerol to 1% -v/v, incubate 30 min at 37C.

#### Sample preparation for metabolomics

M. pneumoniae cells were grown in 6-well culture dishes as described above until reaching 80-90% confluency. Culture medium was aspirated, and cells were rapidly washed twice at 37° C with 1 mL of buffer (75 mM ammonium carbonate at pH 7.4 and 0.1% glucose). After aspiration of washing buffer, plates were immersed in liquid nitrogen to quench metabolism and stored at −80° C for less than 4 days until further processing. After aspiration of washing buffer, plates were immersed in liquid nitrogen to quench metabolism and stored at −80° C for less than 4 days until further processing.

To extract metabolites, plates were placed on a 75° C heating block and 700 μL of extraction solution (70%-v/v ethanol in water at 75° C) were added to each well. After incubating for 3 min, the supernatant was collected and transferred to ice, and the extraction was repeated once. Pooled extracts were dried under vacuum and stored at −80° C prior to metabolomics analyses.

#### Nontargeted metabolomics

All samples were measured in triplicate. Metabolomics samples were analysed by flow-injection time-of-flight MS with an Agilent 6550 iFunnel QToF instrument (Agilent, Santa Clara, CA, U.S.A.) operated in negative ionization mode at 4 GHz high-resolution in a range from 50-1,000 m/z using published settings^25^. The mobile phase was 60:40 isopropanol:water (v/v) and 1 mM NH4F at pH 9.0 supplemented with 10 nM hexakis(1H-, 1H-, 3H-tetrafluoropropoxy)phosphazine and 80 nM taurocholic acid for online mass correction. Spectral processing and ion annotation based on accurate mass within 0.001 Da of metabolites in the *M. pneumoniae* MyMPN database^26^, allowing for [M-H]^-^ and [M+F]^-^ ions and [1x^12^C->1x^13^C] neutral gain and keeping for each metabolite only the ion with lowest *m/z* in case of multiple matching ions, was performed using Matlab R2015b (The Mathworks, Nattick, MA, U.S.A.) as described previously^25^. Metabolomics data were normalized to the summed abundance of a group of amino acids (Ser, Pro, Ala, Val, Thr, Leu/Ile, Met, Phe, Tyr) found to strongly correlate in each sample. In mycoplasma amino acids are not made but imported and they are fairly constant. Therefore, we could use the summed values for the less variable amino acids to normalize. A similar approach is used in free label quantitative proteomics. Subsequently, log2-transformed fold-changes and *P*-values (two-sided *t* tests, with q-values computed from raw p-values to enable false discovery rate adjustment^27^) were calculated to determine relative metabolite abundances compared to control samples and their statistical significance.

#### Targeted metabolomics

Samples were injected into a Waters Acquity UPLC with a Waters T3 column (150 mm x 2.1 mm x 1.8 mm; Waters Corporation, Milford, MA) coupled to a Thermo TSQ Quantum Ultra triple quadrupole instrument (Thermo Fisher Scientific, Waltham, MA) with electrospray ionization. Compound separation was achieved by a gradient of two mobile phases (i) 10 mM tributylamine, 15 mM acetic acid, 5% (v/v) methanol and (ii) 2-propanol. In total, 138 metabolites covering carbohydrate and energy metabolism, amino acid metabolism, nucleotide metabolism and other pathways were targeted. Further details are published elsewhere^28^.

#### Proteomics

Cells were grown in a 25-cm2 flask for 3 days as above, washed with PBS and lysed/collected in 4% SDS, and 0.1 M Hepes·HCl pH 7.5. Samples were reduced with dithiothreitol (15 μM, 30 min, 56°C), alkylated in the dark with iodoacetamide (180 nmols, 30 min, 25°C) and digested with 3 μg LysC (Wako) O/N at 37°C and then with 3 μg of trypsin (Promega) for eight hours at 37°C following FASP procedure (Filter-aided sample preparation 48). After digestion, the peptide mix was acidified with formic acid and desalted with a MicroSpin C18 column (The Nest Group, Inc) prior to LC-MS/MS analysis. The peptide mixes were analysed using a LTQ-Orbitrap Velos Pro mass spectrometer (Thermo Fisher Scientific) coupled to an EasyLC (Thermo Fisher Scientific). Peptides were loaded onto the 2-cm Nano Trap column with an inner diameter of 100 μm packed with C18 particles of 5 μm particle size (Thermo Fisher Scientific) and were separated by reversed-phase chromatography using a 25-cm column with an inner diameter of 75 μm, packed with 1.9 μm C18 particles (Nikkyo Technos). Chromatographic gradients started at 93% buffer A and 7% buffer B with a flow rate of 250 nl min-1 for 5 minutes and gradually increased 65% buffer A and 35% buffer B in 120 min. After each analysis, the column was washed for 15 min with 10% buffer A and 90% buffer B. Buffer A: 0.1% formic acid in water. Buffer B: 0.1% formic acid in acetonitrile.

The mass spectrometer was operated in DDA mode and full MS scans with 1 micro scans at resolution of 60.000 were used over a mass range of m/z 350-2,000 with detection in the Orbitrap. Auto gain control (AGC) was set to 1 E6, dynamic exclusion (60 seconds) and charge state filtering disqualifying singly charged peptides was activated. In each cycle of DDA analysis, following each survey scan the top twenty most intense ions with multiple charged ions above a threshold ion count of 5,000 were selected for fragmentation at normalized collision energy of 35%. Fragment ion spectra produced via collision-induced dissociation (CID) were acquired in the Ion Trap, AGC was set to 5e4, isolation window of 2 m/z, activation time of 0.1 ms and maximum injection time of 100 ms was used. All data were acquired with Xcalibur software v2.2.

Proteome Discoverer software suite (v2.0, Thermo Fisher Scientific) and the Mascot search engine (v2.5, Matrix Science were used for peptide identification. Samples were searched against a *M. pneumoniae* database with a list of common contaminants and all the corresponding decoy entries (87,059 entries). Trypsin was chosen as enzyme and a maximum of three mis-cleavages were allowed. Carbamidomethylation (C) was set as a fixed modification, whereas oxidation (M) and acetylation (N-terminal) were used as variable modifications. Searches were performed using a peptide tolerance of 7 ppm, a product ion tolerance of 0.5 Da. Resulting data files were filtered for FDR < 5 %. Protein Top 3 areas were calculated with unique peptides per protein.

#### Data and model management

All omics data, modelling files as well as a backup of modelling pipeline and simulation outputs are available via the Seek data and model management platform for maximum reproducibility (http://doi.org/10.15490/FAIRDOMHUB.1.INVESTIGATION.133.3) ^29^,^30^. SBtab^31^, a tabular exchange format was used to add minimal information compliant with the Minimal Information Requirements In the Annotation of Models (MIRIAM)^32^ compliant annotation and to add Systems Biology Ontology (SBO) identifiers^33^ for metabolites, reactions and parameters in the model.

#### Data analysis

Relative metabolite measurements were log10-transformed following the recommendation of Jauhiainen et al ^34^. Pearson correlations between metabolites were computed using R v3.4.2^35^. To remove batch effects in the metabolites measurement, values were normalized by dividing them by the median measured metabolite value per batch^36^. Fold Change (FC) metabolite measurements for the 40 independent samples were analysed through Principal Component analysis using the prcomp package. Pearson correlation between samples and metabolites were calculated and used to generate heatmaps of sample correlations and metabolite correlations. We used the metabolite correlation matrix for [M+F]-ion data and filtered on correlations with at least p-value cut-off of 0.001. We calculated the Euclidean distance using the complete linkage method and used Hierarchical clustering and cut tree to identify 6 clusters in the metabolite correlation data. Absolute measurements needed for simulations with the model were obtained by multiplying relative metabolite values from [M-H]^-^ measurements with quantitative metabolite measurements at 24 h. Similarly, enzyme concentrations of samples with overexpression (OE) of enzymes were obtained by multiplying relative measurements for these mutants with absolute measurements of the wild type at their respective time point. These computations were performed using Python. Additionally, we calculated Pearson correlation between metabolite concentration and estimated growth of 24 time-series samples (P-value <0.05) applying Benjamin Hocheman Multiple testing correction.

#### Model construction and numerical implementation

A dynamic model was build of glycolysis an pyruvate metabolism. A base model containing all reactions in glycolysis, pyruvate metabolism from the MyMPN database^26^.Different additions to this model were tested such as individual and combined additions of 1) an ATPase reaction, 2) LDH inhibition by oxygen and 3) a NoxE reaction for NAD regeneration using oxygen. The tested models include i) the base model ii) the base model and the ATPase reaction iii) base model with NoxE reaction iv) base model with both ATPase and NoxE reaction v) based model with NoxE reaction and LDH inibhition by oxygen and vi) the base model with all three modifications. In case intermediate metabolites were not measurable, reactions were lumped in a single reaction. This was the case for Phosphoglycerate kinase (PGK), Glycerate phosphomutase (GMP) and enolase (ENO). These three reactions were combined in reaction re07 lumping the enzymatic reactions of PGK&GMP&ENO. Similarly, phosphotransacetylase (PTA) and acetate kinase (ACK) were combined in reaction re10 which lumps the enzymatic reactions of PTA&ACK.

Allosteric control was assumed to be similar to allosteric control in *Lactococcus lactis* as presented in the model by Costa *et al*^37^ due to the lack of *MPN* specific information on allosteric control. Allosteric control includes three activator and five inhibitor effects. Reactions were modelled using modular rate laws except for transport reactions for which Hill type kinetics were used. Enzyme concentrations were included as reaction parameter to allow model predictions at varying protein concentrations. The base model contains 10 equations and 72 parameters of which 10 represent experimentally determined enzyme concentrations, 5 represent equilibrium constants (Keq) and 1 is a Hill coefficient. The remaining 56 parameters represent Michaelis-Menten constants, activation constants and inhibition constants which are not known for *MPN*. The model was built using COPASI^38^.

#### Initial parameter values

Proteomics measurements for 6, 24 and 48-hour timepoints were used as estimates for enzyme concentrations at respective time points. In case of multi-subunit enzymes, the most abundant single copy subunit was chosen to represent the enzyme concentration. Many lower abundance subunits are only expressed under specific conditions and as such are not representative of the abundance of these glycolysis enzymes. We also tested using the average, but no major differences were found. Equilibrium constants were gathered from www.equilibrator.org assuming an ionic strength of 0.1 and a pH of 7^39^. Initial values for Monod constants and allosteric control constants were randomly selected between 0.01 and 100 fold of the observed metabolite concentrations at 24h. An overview of the six models reactions and equations can be found in Supplementary file 1 A.

#### Model selection and parameter estimation

The base model has 72 parameters of which 56 are unknown while model that includes ATPase, NoxE and O_2_ inhibition of LDH has 80 parameters of which 63 are unknown. Parameter estimation was performed training the models on metabolomics steady state data obtained from growth curve samples for 6h, 24h and 48h time points grown on medium containing 60 mM of glucose. These samples were selected as training data due to the completeness of the data available for these three times points. Only for these three samples, measurements for all 17 metabolites present in the model were available. In addition, protein copy number, glucose uptake, lactate secretion, and acetate secretion measurements were available for these three samples. Steady state concentration for 6h, 24h and 48h grown on a lower glucose concentration of 10 mmol were used as internal validation data. Internal validation data is used by COPASI to stop the parameter estimation algorithm from overfitting parameters to the training data. The large time interval between the samples means that metabolite concentrations in each sample can be assumed to be independent from the concentrations of the other samples. Therefore, each sample was treated as an independent steady state.

COPASI’s^38^ build in Genetic programming algorithm was used to estimate parameters using a maximum of 1000 generations with a population size of 500 models with normalized sum of squares as weights. 100 independent parameter estimations were run per model. Optimal parameters were searched within a range of 10^-2^-10^2^-fold of the observed metabolite concentration at 24h for Monod constants and allosteric control constants while maximum reaction velocity values were searched within a range of 10^-2^-10^3^. The performance of the six models were compared based on the distribution of the mean square error values for each of the 100 parameter estimations.

#### Simulations, local and global sensitivity analysis

The model was used to predict steady state concentrations for 40 independent samples comprised of OE and knock out (KO) mutants, perturbations, and time-series measurements in different growth conditions measured in triplicate. In these simulations, the input for the model was concentration data of 11 metabolites: acetyl coenzyme A (AcCoA), acetate (ACE), adenosine diphosphate (ADP), adenosine triphosphate (ATP), coenzyme A (CoA), diacylglycerol phosphate (DGP), fructose 6-phosphate (F6P), fructose-1,6-bisphosphatase (FBP), glucose 6-phosphate (G6P), glyceraldehyde-3-phosphate (GAP), lactate (LAC), nicotinamide adenine dinucleotide (NAD), reduced nicotinamide adenine dinucleotide (NADH), phosphoenolpyruvate (PEP), orthophosphate (Pi_Int), pyruvate (PYR), external glucose (GLC_Ext). Measurements for NAD and NADH are approximate. For each sample, 1000 steady state simulations were performed while sampling from the log normal distribution of metabolite measurements. By comparing sampled measurements and sampled simulation values, measurement error and its propagation are incorporated in model predictions.

Not all metabolites present in the model were measured for all independent samples. Reference values from measurements taken at 6h, 24h and 48h of growth on high glucose concentrations used to train the model were used to set the initial concentration of NAD, NADH and orthophosphate. Reference values were also used for CoA, Acetyl-CoA, and Lactic Acid (LAC) for some of the independent samples (Supplementary Material 2).

To compare the error between simulated and measured metabolite concentrations in a consistent manner, we used the symmetric Mean Absolute Percentage Error (sMAPE). sMAPE is a measure of prediction accuracy used for forecasting methods. This method has the advantage of providing an equal error to positive and negative errors for log normal distributed data such as metabolite measurements and predictions^40^.

We performed global sensitivity analysis for all kcat, Monod constants, activation and inhibition constants using a 100,000 Latin Hypercube sampling^41,42^. Samples were constructed by sampling from the log linear distribution of each parameter’s respective search range.

The above described operations were performed using Python 3.6.5 with the Tellurium 2.0.18 and Roadrunner 1.4.24 high performance SBML simulation and analysis libraries^43,44^. The pyDOE package was used for Latin Hypercube sampling. Conda version 4.3.21 was used for package management.

#### Modelling oxygen diffusion

Oxygen concentrations were calculated based on the initial oxygen concentration in the culture flasks and the acetate production rate which requires NAD to be regenerated from NADH by the oxygen dependent reaction catalysed by NoxE. The initial oxygen concentration was calculated with the ideal gas law, using the temperature used in cultivation (37 degrees Celsius) and atmospheric pressure. Volumes, surface area height of the medium were calculated based on the medium and inoculant volume and the specifications of the T300 cell culture flask^45^.

Diffusion of oxygen from the head space into the medium was calculated using the Wilke and Chang correlation^46^ while Fick’s law^47^ was used to calculate the diffusion of oxygen to the bottom of the flask at 6h, 24h, 48h and 96 hours of growth. The calculated oxygen concentrations were added to the metabolomics measurements for the 95 independent samples.

## Results

Datasets were collected growing *MPN* in a large number of conditions.

*M. pneumoniae* was grown in suspension until sedimenting after 6 h of incubation in rich medium in non-aerated, non-stirred conditions mimicking its host environment. At several time points during the growth of wild-type *MPN*, samples were taken for metabolomics, biomass, pH and acetate concentration measurements. In addition, relative metabolite concentrations were measured by untargeted metabolomics for 40 different samples from environmental perturbations, genetic mutations and at 6h, 24h, 48h and 96 hours of growth. The metabolomics data and targeted proteomics data for these samples is available in Supplementary file 2.

Among these 40 datasets, there were data corresponding to OE mutants for all glycolysis and pyruvate metabolism enzymes except for pyruvate dehydrogenase (PDH) of which the complex is large to clone and OE, as well as for the KO of LDH (Mpn674). The fold change in mRNA and or protein concentrations for genes targeted in each mutant were measured. Of the 40 datasets, 17 are mutants that target enzymes for which a reaction is present in the model. Of these mutants 2 are KO mutants and 14 are OE mutants and 1 is a combined KO and OE mutant (Table 1). Additionally, there are 6 mutants targeting enzymes in in the pentose phosphate pathway which is connected to the glycolysis via F6P.

**Table 1.**
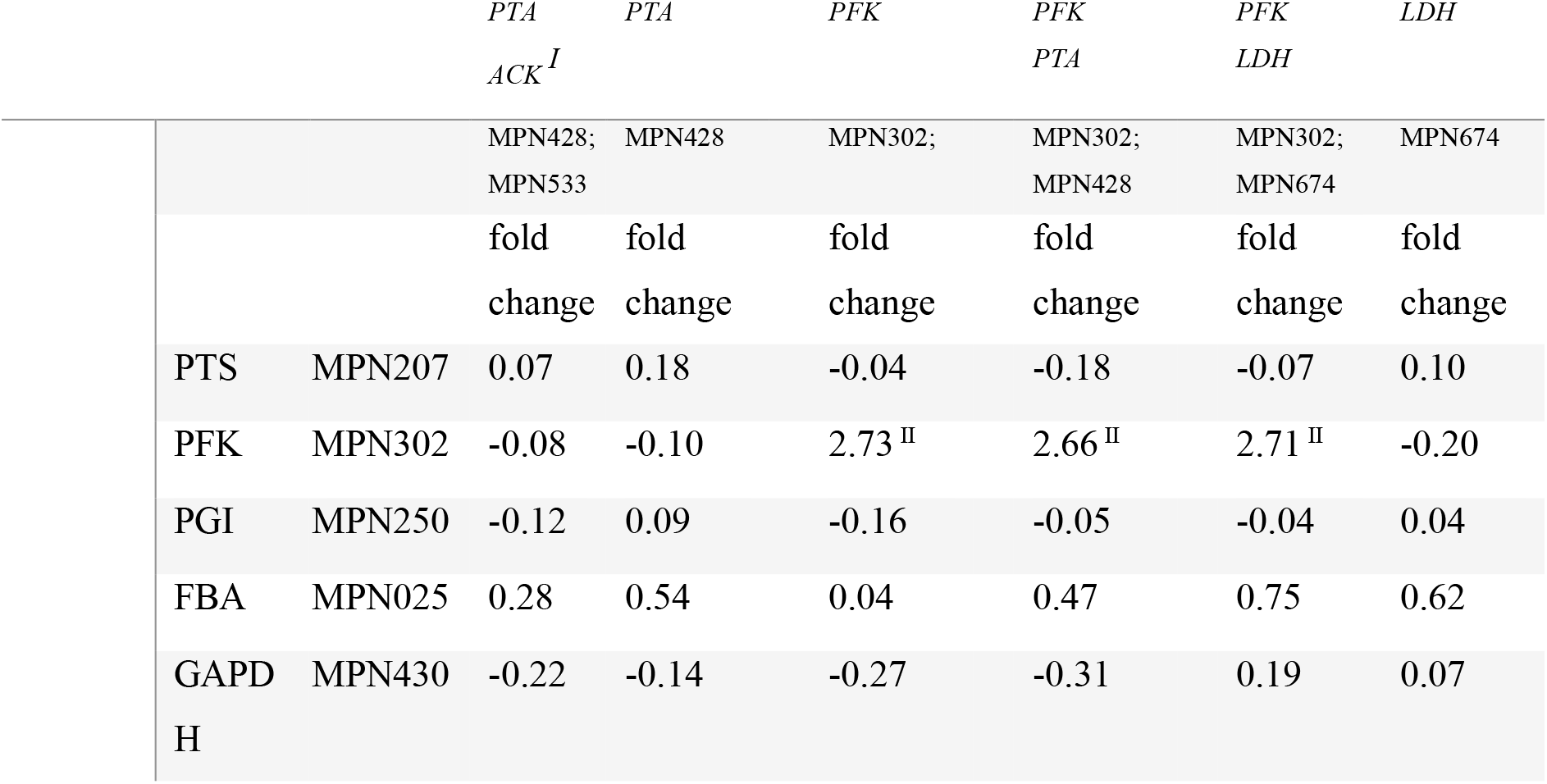
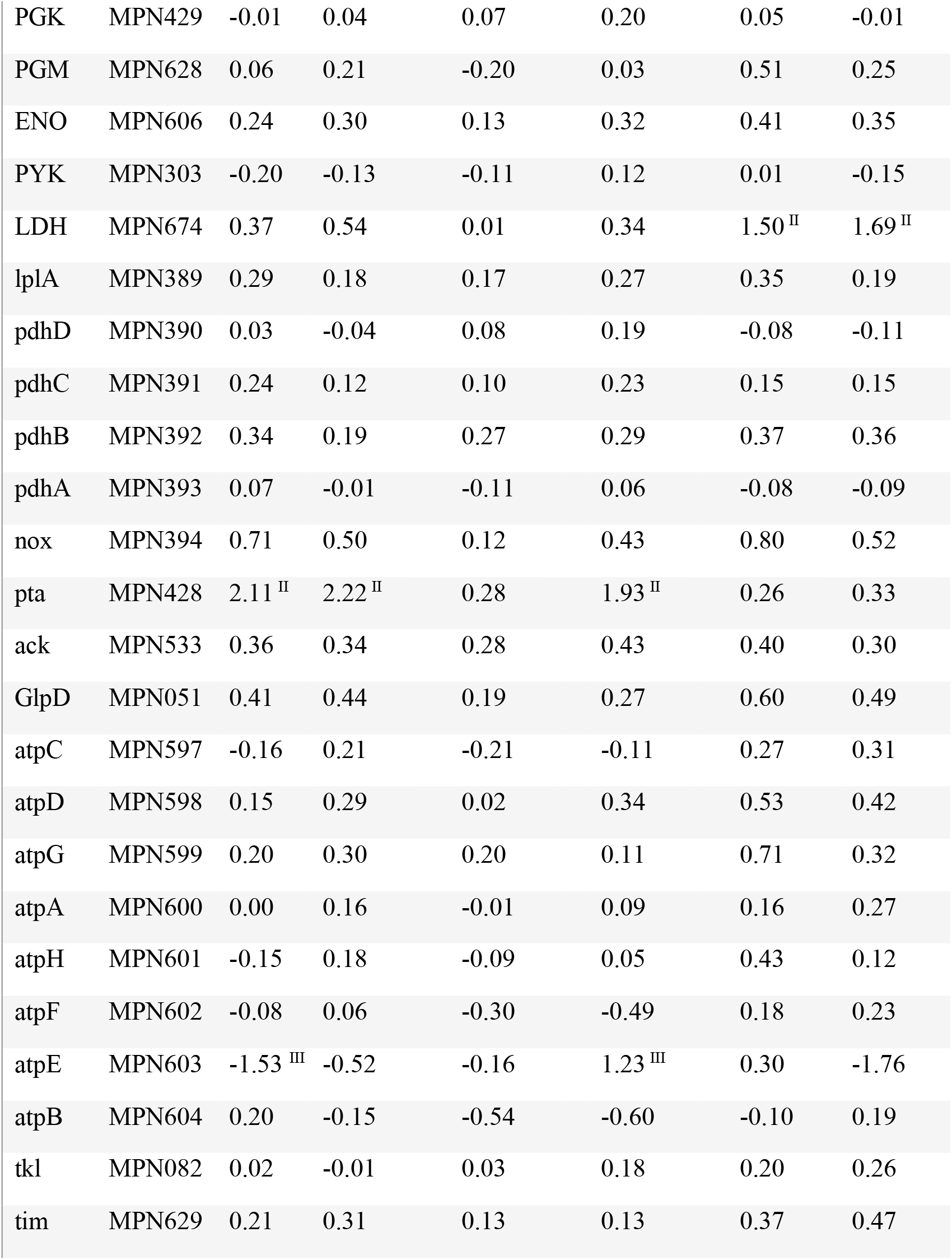
Log2 Fold change expression values of OE mutants. I: Mutant PTA ACK did not show any OE of ACK. II: OE are significantly different from the wild type. III: OE values are significant, but the changes are noisy.

Changes in mRNA and protein concentration of enzymes targeted in over expression (OE) and knock out (KO) mutants were also measured. Table 1 gives an overview of these conditions and mutants.

Unless stated otherwise, relative metabolite concentrations were measured at steady state in non-aerated conditions. In cases where different conditions were used, or where additional omics data were measured, this is indicated in Table 1. Targeted metabolomics were used to measure protein concentration for all OE mutants targeting central carbon metabolism enzymes.

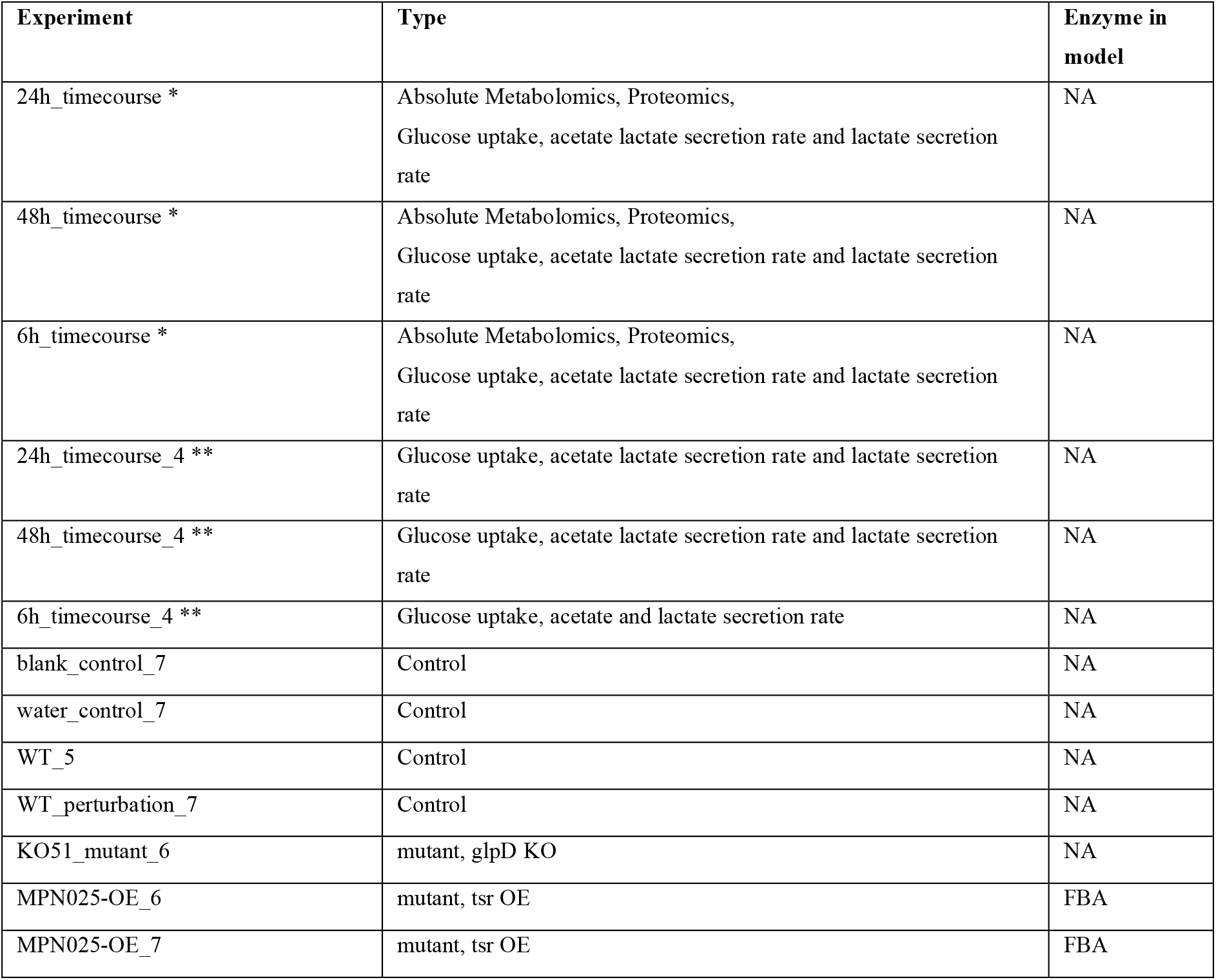
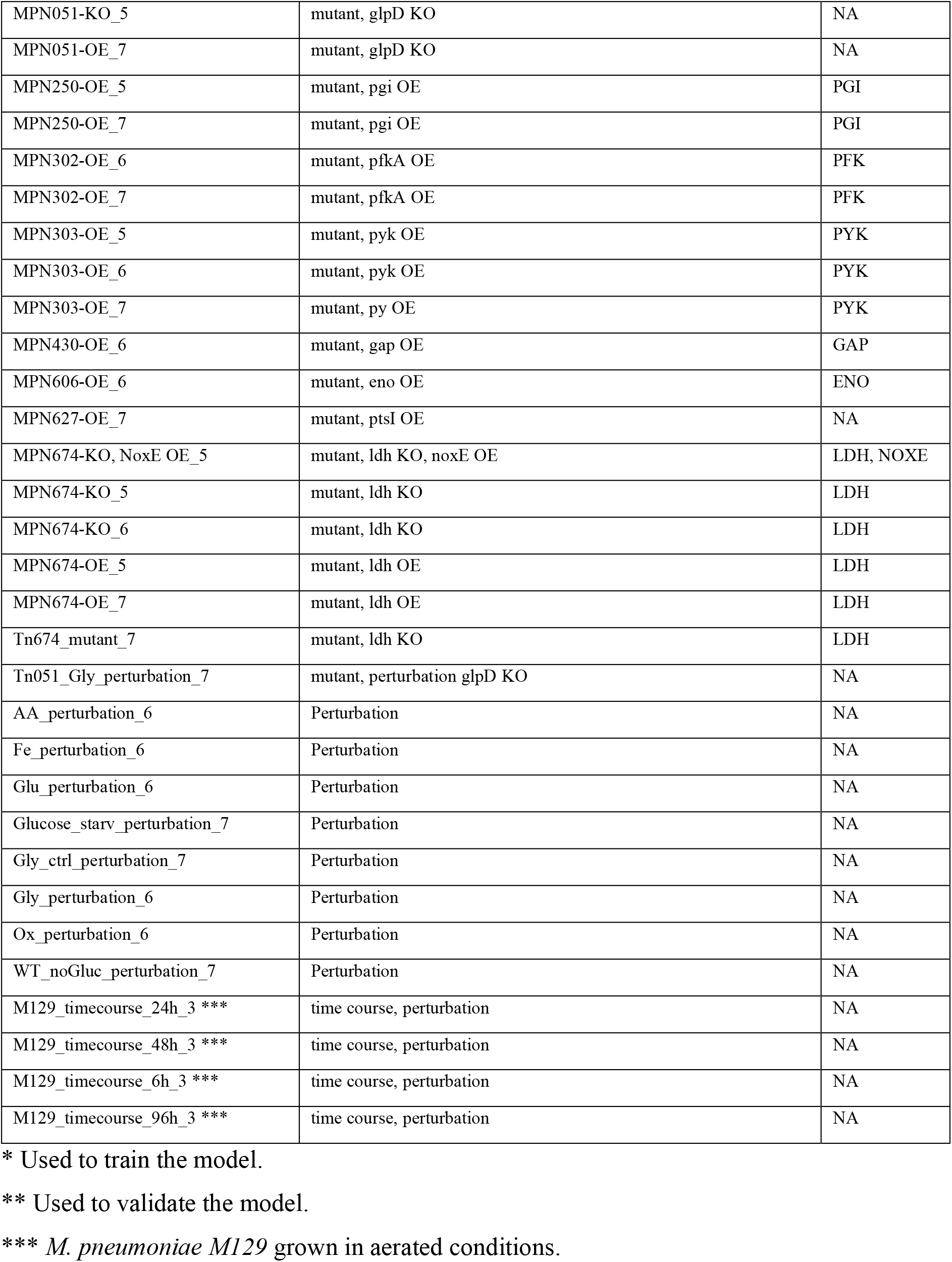

| Experiment | Type | Enzyme in model |
| --- | --- | --- |
| 24h_timecourse * | Absolute Metabolomics, Proteomics,<br>Glucose uptake, acetate lactate secretion rate and lactate secretion rate | NA |
| 48h_timecourse * | Absolute Metabolomics, Proteomics,<br>Glucose uptake, acetate lactate secretion rate and lactate secretion rate | NA |
| 6h_timecourse * | Absolute Metabolomics, Proteomics,<br>Glucose uptake, acetate lactate secretion rate and lactate secretion rate | NA |
| 24h_timecourse_4 ** | Glucose uptake, acetate lactate secretion rate and lactate secretion rate | NA |
| 48h_timecourse_4 ** | Glucose uptake, acetate lactate secretion rate and lactate secretion rate | NA |
| 6h_timecourse_4 ** | Glucose uptake, acetate and lactate secretion rate | NA |
| blank_control_7 | Control | NA |
| water_control_7 | Control | NA |
| WT_5 | Control | NA |
| WT_perturbation_7 | Control | NA |
| KO51_mutant_6 | mutant, glpD KO | NA |
| MPN025-OE_6 | mutant, tsr OE | FBA |
| MPN025-OE_7 | mutant, tsr OE | FBA |
| MPN051-KO_5 | mutant, glpD KO | NA |
| MPN051-OE_7 | mutant, glpD KO | NA |
| MPN250-OE_5 | mutant, pgi OE | PGI |
| MPN250-OE_7 | mutant, pgi OE | PGI |
| MPN302-OE_6 | mutant, pfkA OE | PFK |
| MPN302-OE_7 | mutant, pfkA OE | PFK |
| MPN303-OE_5 | mutant, pyk OE | PYK |
| MPN303-OE_6 | mutant, pyk OE | PYK |
| MPN303-OE_7 | mutant, py OE | PYK |
| MPN430-OE_6 | mutant, gap OE | GAP |
| MPN606-OE_6 | mutant, eno OE | ENO |
| MPN627-OE_7 | mutant, ptsI OE | NA |
| MPN674-KO, NoxE OE_5 | mutant, ldh KO, noxE OE | LDH, NOXE |
| MPN674-KO_5 | mutant, ldh KO | LDH |
| MPN674-KO_6 | mutant, ldh KO | LDH |
| MPN674-OE_5 | mutant, ldh OE | LDH |
| MPN674-OE_7 | mutant, ldh OE | LDH |
| Tn674_mutant_7 | mutant, ldh KO | LDH |
| Tn051_Gly_perturbation_7 | mutant, perturbation glpD KO | NA |
| AA_perturbation_6 | Perturbation | NA |
| Fe_perturbation_6 | Perturbation | NA |
| Glu_perturbation_6 | Perturbation | NA |
| Glucose_starv_perturbation_7 | Perturbation | NA |
| Gly_ctrl_perturbation_7 | Perturbation | NA |
| Gly_perturbation_6 | Perturbation | NA |
| Ox_perturbation_6 | Perturbation | NA |
| WT_noGluc_perturbation_7 | Perturbation | NA |
| M129_timecourse_24h_3 *** | time course, perturbation | NA |
| M129_timecourse_48h_3 *** | time course, perturbation | NA |
| M129_timecourse_6h_3 *** | time course, perturbation | NA |
| M129_timecourse_96h_3 *** | time course, perturbation | NA |

\* Used to train the model.
\*\* Used to validate the model.
\*\*\* *M. pneumoniae* M129 grown in aerated conditions.

We explored the measurements of metabolite concentrations for the various samples shown in Table 1. Similar conditions as well as KO of genes in the same pathway cluster together.

An example of this is the clustering of all M129 samples which are the only samples grown in aerated conditions (Figure 1).

**Figure 1.**
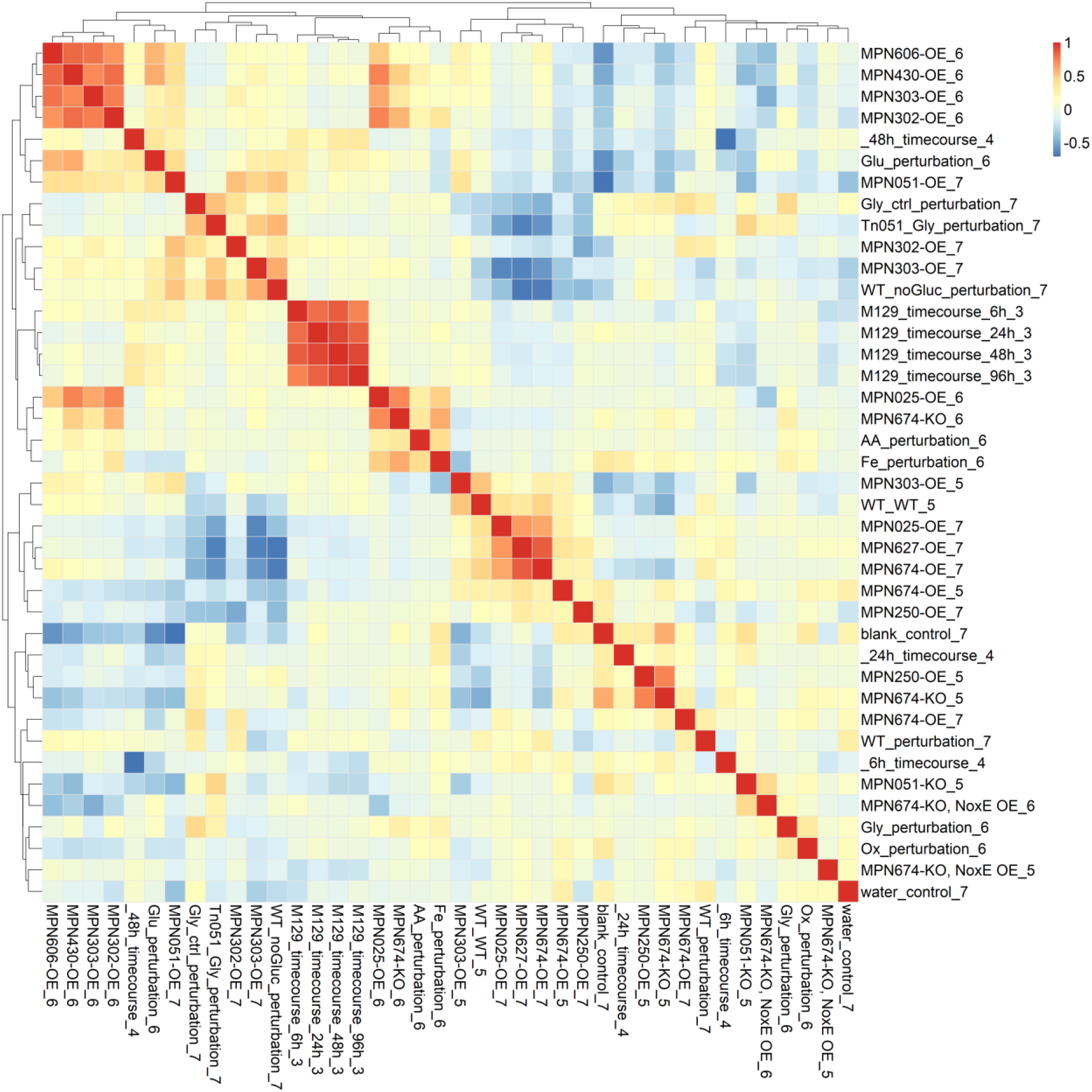
Heatmap of mutant and perturbation sample clustering based on their metabolite profile.

Two clusters exist of OE mutants targeting glycolysis. The first cluster contains an OE mutant of FBP, phosphotransferase MPN627 involved in mannitol and mannose uptake^8^ and LDH. This clustering corroborates the assumed allosteric activation of LDH by FBP. The second cluster contains OE mutants of ENO, GAP, PYK and PFKA.

Another interesting cluster contains *MPN* perturbation, growth without oxygen and growth without amino acids which cluster together with both an LDH KO and FBA OE. These four samples have in common that the conditions are growth inhibiting. The clustering of FBA OE which degrades FBP together with LDH KO can be explained by the positive allosteric control of FBP on LDH.

Two annotation techniques were used to measure metabolite abundance, [M+F]^-^ ion and [M-H]^-^ ion detection^48^. Some differences are present in the correlations between individual metabolites in the [M+F] ^-^ ion and [M-H]^-^ data, however, clustering of samples for both detection techniques is highly similar. A heatmap that compares both [M+F] ^-^ ion and [M-H] ^-^ data can be found in the Supplementary files 1B.

In addition to clustering of samples based on their relative metabolite concentrations we studied the clustering of metabolites of these samples. We use the Pearson correlations between metabolites to build a network of metabolite-metabolite interactions (Figure 2). We identified a neatly defined cluster structure. The largest cluster contains sn-glycero-3-phosphocholine, CDP-Choline, folate and methionine cycle metabolites, orthophosphate, all nucleotide three phosphates (ATP, CTP, UTP) and pentose phosphate metabolism (PPP) metabolites. This cluster suggest a link between sn-glycero-3-phosphocholine catabolism, energy production, and nucleobase salvaging by phosphorylation and de-oxidation. Higher concentrations of PRRP associated with these metabolites are needed to convert nucleobases into ribonucleotides, while increase in ATP is needed for phosphorylation of deoxyribonucleotides:

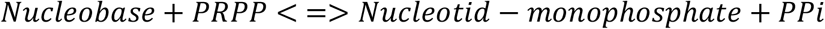

**Figure 2.**
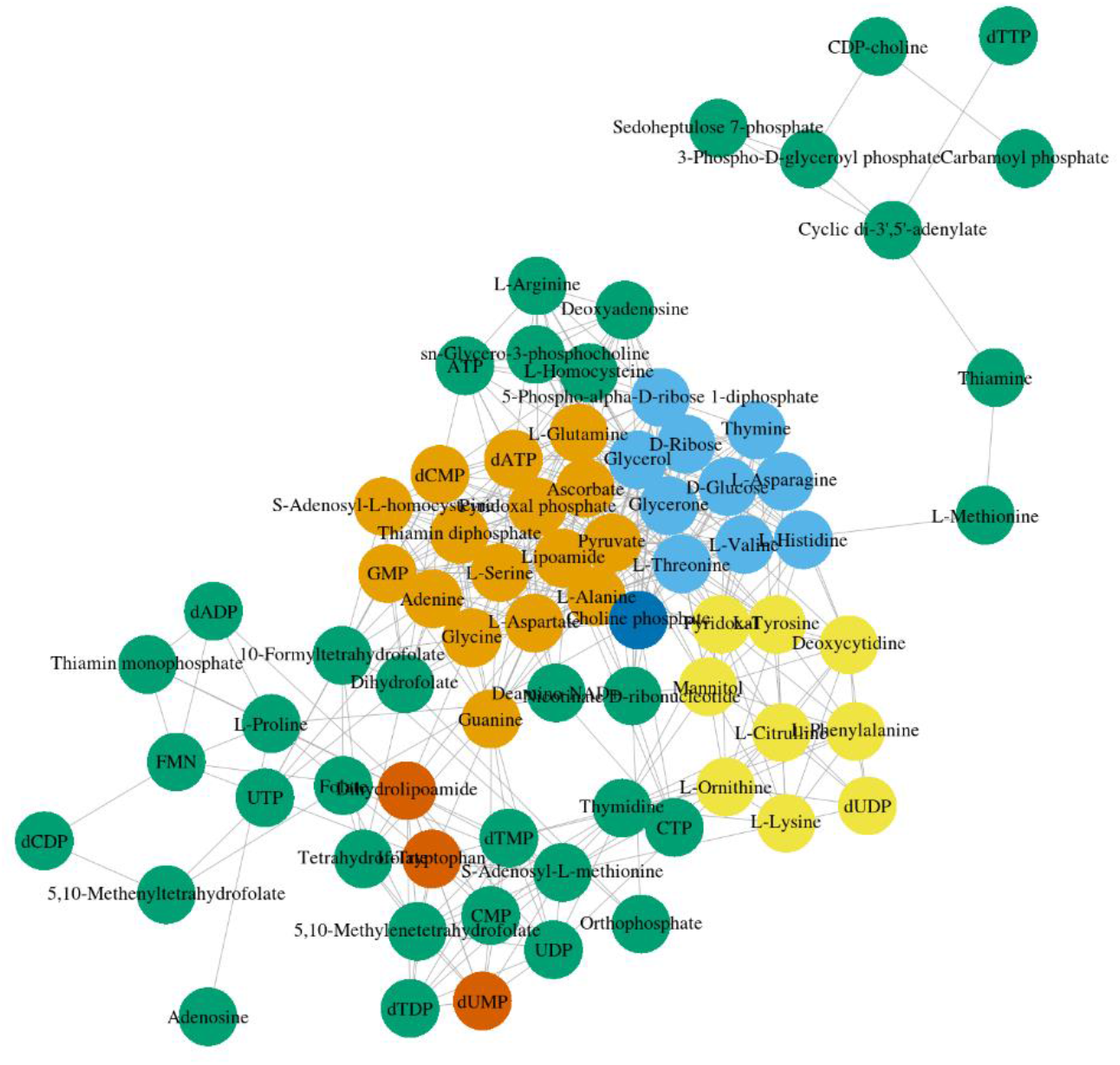
Metabolite correlation network based the Pearson correlation of metabolite measurements of the 40 independent samples. The 6 clusters were assigned based on hierarchical clustering of the correlation data.

In humans, phosphatidylcholine (lecithin) is used as main nutritional source for one carbon metabolism (CH_3_)^49^ One carbon metabolism and its relation to nucleotide synthesis in cancer cells has been extensively studied^48^. It has been argued that in human cancer cells, glycolysis can produce enough energy for growth by diverting its flux to other metabolic pathways including one-carbon metabolism. Indeed, several reactions of one-carbon metabolism contribute to ATP and NADPH production. Similarly, it has been argued that phosphatidyl choline plays a major role in the nutrition of *MPN* as it is by far the most abundant available carbon source in the lungs and for these reasons is also used as carbon and nitrogen source by pathogens like *P. aeruginosa*^50,51^ and claimed to be used as carbon source by *MPN*^52^.

The second largest cluster contains alanine, aspartate, glutamate, serine, pyridoxal phosphate (vitamin B6), S-adenosyl-L-homocysteine, glycine, lipoamide, pyruvate as well as adenine, guanine, GMP, dATP and dCTP. Pyruvate is positively correlated with dATP and dCTP which are needed for DNA synthesis. This cluster is strongly associated to the sub cluster of sn-Glycero-3-phosphocholine and tetrahydrofolate metabolites from the first cluster.

Based on the observed cluster, there are three main lessons to be learned. Firstly, clustering of sn-Glycero-3-phosphocholine with ATP, CTP and UTP metabolite correlation profiles supports the theory^50^ that glycerol-3-phosphate derived from sn-Glycero-3-phosphocholine functions as a carbon and energy source for *MPN*. Addition of phosphatidylcholine to a defined minimal medium for *MPN* indeed optimizes growth^22^. Secondly, sn-Glycero-3-phosphocholine clusters together with folate and methionine cycle one carbon metabolites and as such is likely the main one carbon donor in *MPN* metabolism. Thirdly, positive correlation between CDP-choline, 3-phospho-D-glyceroyl-phosphate and sedoheptulose-7 phosphate, as well as between sn-Glycero-3-phosphocholine and the cluster containing PRPP indicate a link between sn-Glycero-3-phosphocholine and pentose phosphate metabolism.

### Over Expression of glycolytic enzymes

To further study the control of different glycolytic enzymes on central carbon metabolism we, analysed the fold change of enzymes in glycolysis and pyruvate metabolism when OE single as well as some combinations of glycolytic enzymes (Table 1)

What we see is that when OE these other enzymes in the pathway don not widely change. These results suggests that allosteric regulation and circuit topology might play a great role on the control of central carbon metabolism of *MPN*.

### Exploration of metabolite’s concentration at steady state: study of associations

We build a dynamic model of central carbon metabolism including glycolysis and pyruvate metabolism. We trained this model with a limited subset of data and use the model to identify key regulatory elements in glycolysis.

Different additions to this model were tested such as individual and combined additions of 1) an ATPase reaction to account for varying ATP demand, 2) LDH inhibition by oxygen and 3) a NoxE reaction for NAD regeneration using oxygen. The tested models include i) the base model ii) the base model and the ATPase reaction iii) base model with NoxE reaction iv) base model with both ATPase and NoxE reaction v) based model with NoxE reaction and LDH inhibition by oxygen and vi) the base model with all three modifications.

We found the model with addition of NoxE to have the best fitting in multiple parameter estimations, therefore we kept the addition of the NoxE to the model we used for further simulations/

In case intermediate metabolites were not measurable, reactions were lumped in a single reaction. This was the case for Phosphoglycerate kinase (PGK), Glycerate phosphomutase (GMP) and enolase (ENO). These three reactions were combined in reaction re07 lumping the enzymatic reactions of PGK&GMP&ENO. Similarly, phosphotransacetylase (PTA) and acetate kinase (ACK) were combined in reaction re10 which lumps the enzymatic reactions of PTA&ACK.

Allosteric control was assumed to be similar to allosteric control in *Lactococcus lactis* as presented in the model by Costa *et al*^37^ due to the lack of *MPN* specific information on allosteric control. Allosteric control includes three activator and five inhibitor effects. Reactions were modelled using modular rate laws except for transport reactions for which Hill type kinetics were used. Enzyme concentrations were included as reaction parameter to allow model predictions at varying protein concentrations. The base model contains 10 equations and 72 parameters of which 10represent experimentally determined enzyme concentrations, 5 represent equilibrium constants (Keq) and 1 is a Hill coefficient. The remaining 56 parameters represent Michaelis-Menten constants, activation constants and inhibition constants which are not known for *MPN*. The model was built using COPASI^38^.

An overview of the model’s reactions and equations and the additions tested can be found in in Supplementary material 1A.

We compared the mean square error, for all parameter sets (Figure 4). Models that include ATPase (models 2, 4 and 6) have on average the largest error and took the most iterations to reach a stable solution. The addition of NoxE was shown to reduce the error in model predictions (see Figure 4). A zoomed in version of Figure 4 as well as correlation analysis of parameter sets is available in Supplementary file 1B. Since the model with addition of NoxE performed best, further analyses were continued using this model. This model loaded with the best performing parameter set was deposited in BioModels^54^ and assigned the identifier MODEL1911200003.

**Figure 3:**
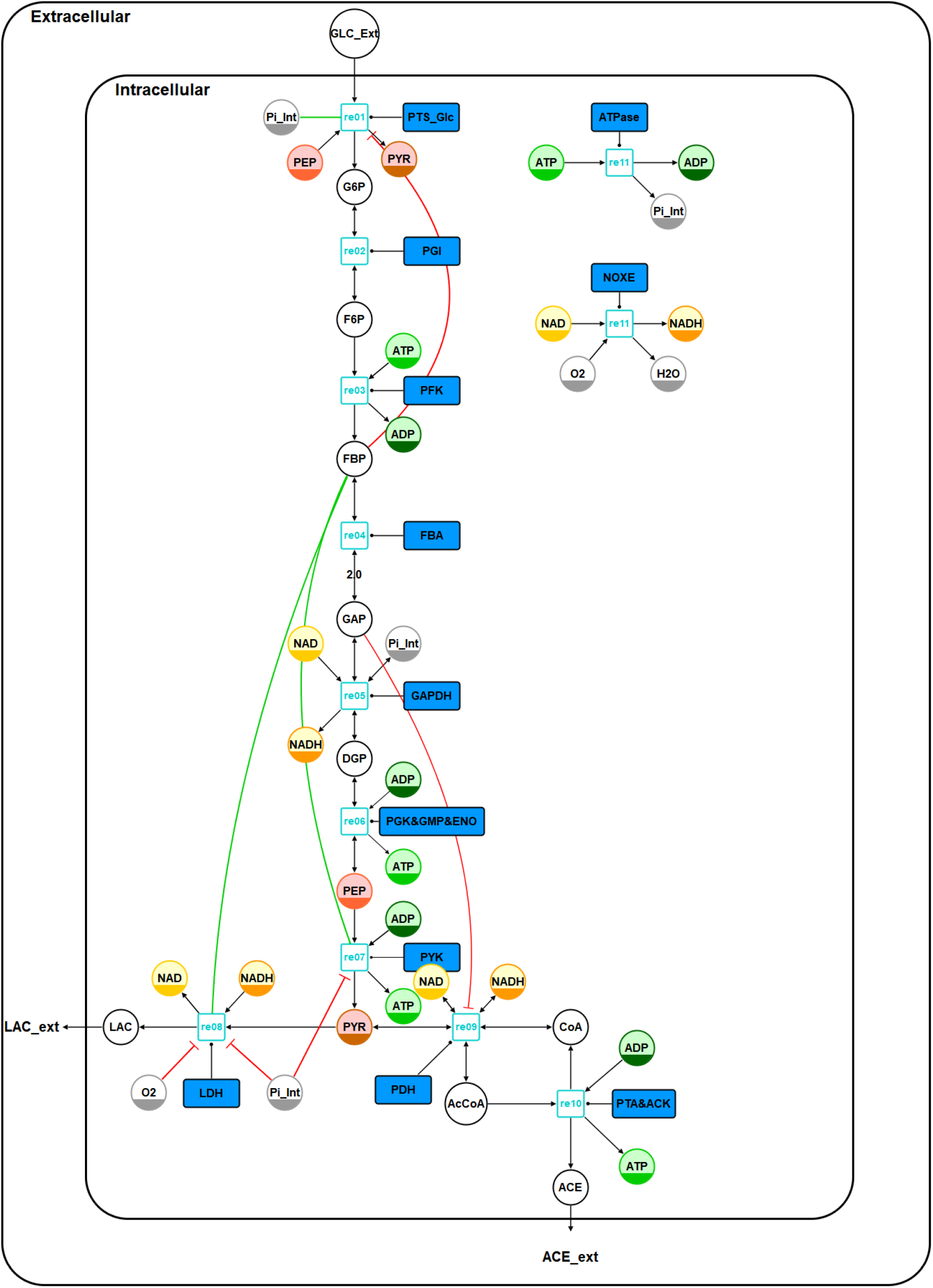
Schema of the model. The diagram meets the Systems Biology Graphical Notation (SBGN) standard^53^ with the exception of the LAC and ACE export arrows which in the model are present as syncs for LAC and ACE: Arrowheads represent reactions (black arrowhead end), catalysis (open circle end), activation (green) and inhibition (red). Circles indicate metabolites. Half-filled circles are clone makers to indicate the metabolites appears multiple times in the diagram (green=adenine nucleotides, yellow=redox equivalents, red=PEP/PYR). Blue filled rectangles describe macromolecules (enzymes, transporters) that catalyse a reaction. In case the reaction stoichiometry is different to 1, the reaction stoichiometry is given as text at the reaction arrow. Blue empty rectangles indicate reaction identifiers in the model. Metabolite abbreviations: AcCoA = acetyl coenzyme A, ACE = acetate, ADP = adenosine diphosphate, ATP = adenosine triphosphate, CoA = coenzyme A, DGP = diacylglycerol phosphate, F6P = fructose 6-phosphate, FBP = fructose-1,6-bisphosphatase, G6P = glucose 6-phosphate, GAP = glyceraldehyde-3-phosphate, LAC = lactate, NAD = nicotinamide adenine dinucleotide, NADH = reduced nicotinamide adenine dinucleotide, PEP = phosphoenolpyruvate, Pi_Int = orthophosphate, PYR = pyruvate (PYR), GLC_Ext = external glucose.

**Figure 4.**
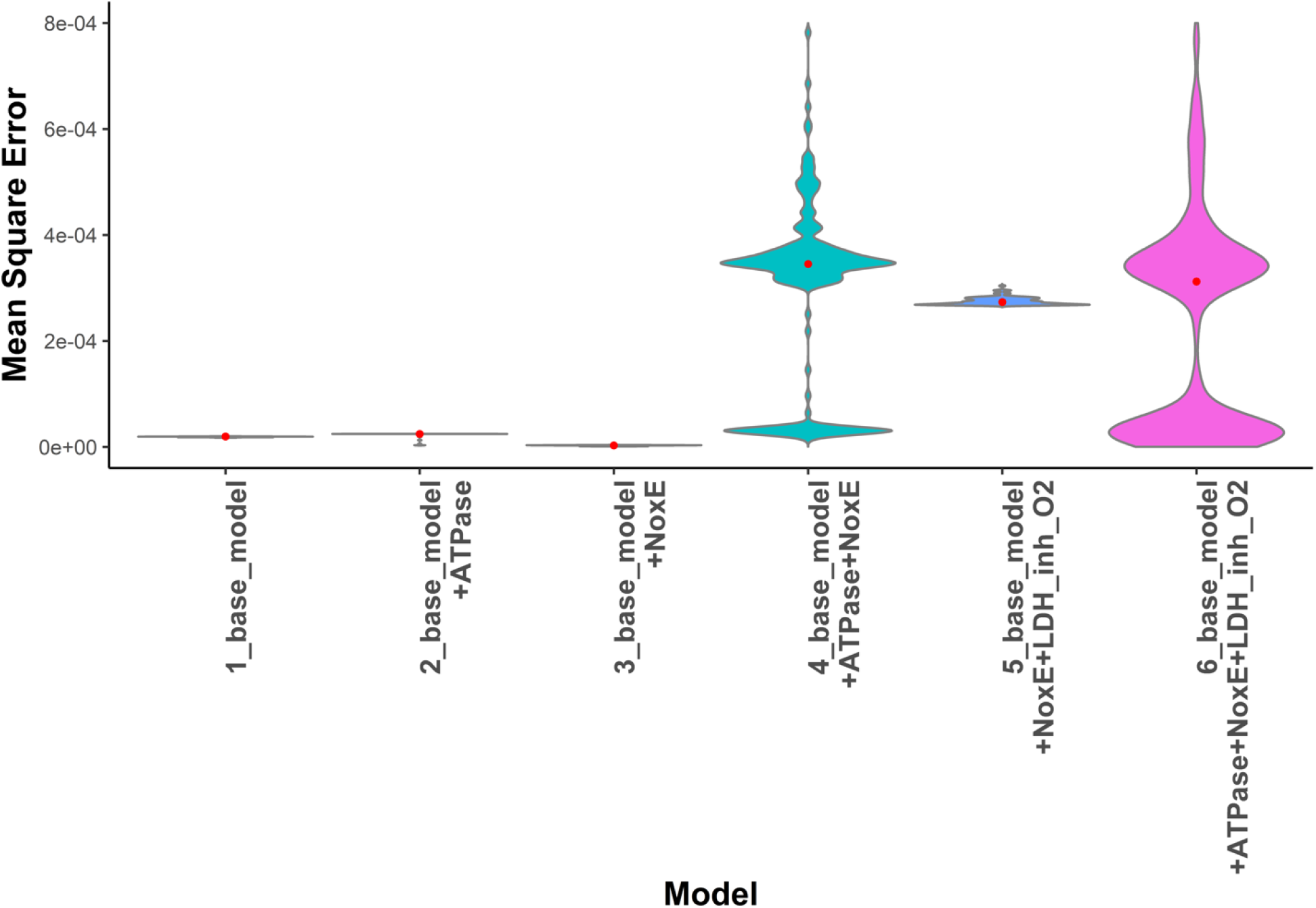
Violin plot comparing the mean square error, for all 100 parameter sets for six models. The red dot indicates the median value.

### Simulating perturbations, KO and OE mutants

We used the trained model to predict steady state metabolite concentrations for 40 independent samples. Sample’s metabolite concentration mean and standard deviation values were determined by measuring samples in triplicate. In addition to being measured in triplicates, biological replicates were available for all OE targeting glycolysis and pyruvate metabolism.

For each of the 40 samples, 1000 independent simulations were performed with slightly different initial concentrations to explore the impact of biological variability and uncertainty associated to measurement error rates. On average the model predicted these samples with reasonable accuracy (Figure 4).

The largest error in predicted metabolite concentrations occur for the samples of MPN129 growth in aerated conditions. These results were to be expected as these samples are clearly forming an outlier since the growth conditions are so vastly different from the other samples. From samples corresponding to perturbations in growth condition, the oxidative stress perturbation (0.15% H_2_O_2_) had the largest prediction error.

The relatively simple model here presented reproduces the states attained under a broad range of perturbations such as glucose concentrations up to a factor 10 lower as compared to the training data. The use of proteomics data is most likely one of the main reasons the model simulates OE and KO mutants of enzymes in glycolysis relatively well. The calculated metabolite concentrations are, on average within a factor 3 of measured values for all mutants and perturbation samples, which is comparable to the accuracy of the training set. An example of measured and simulated values of sampled simulations can be seen in Figure 6.

**Figure 5.**
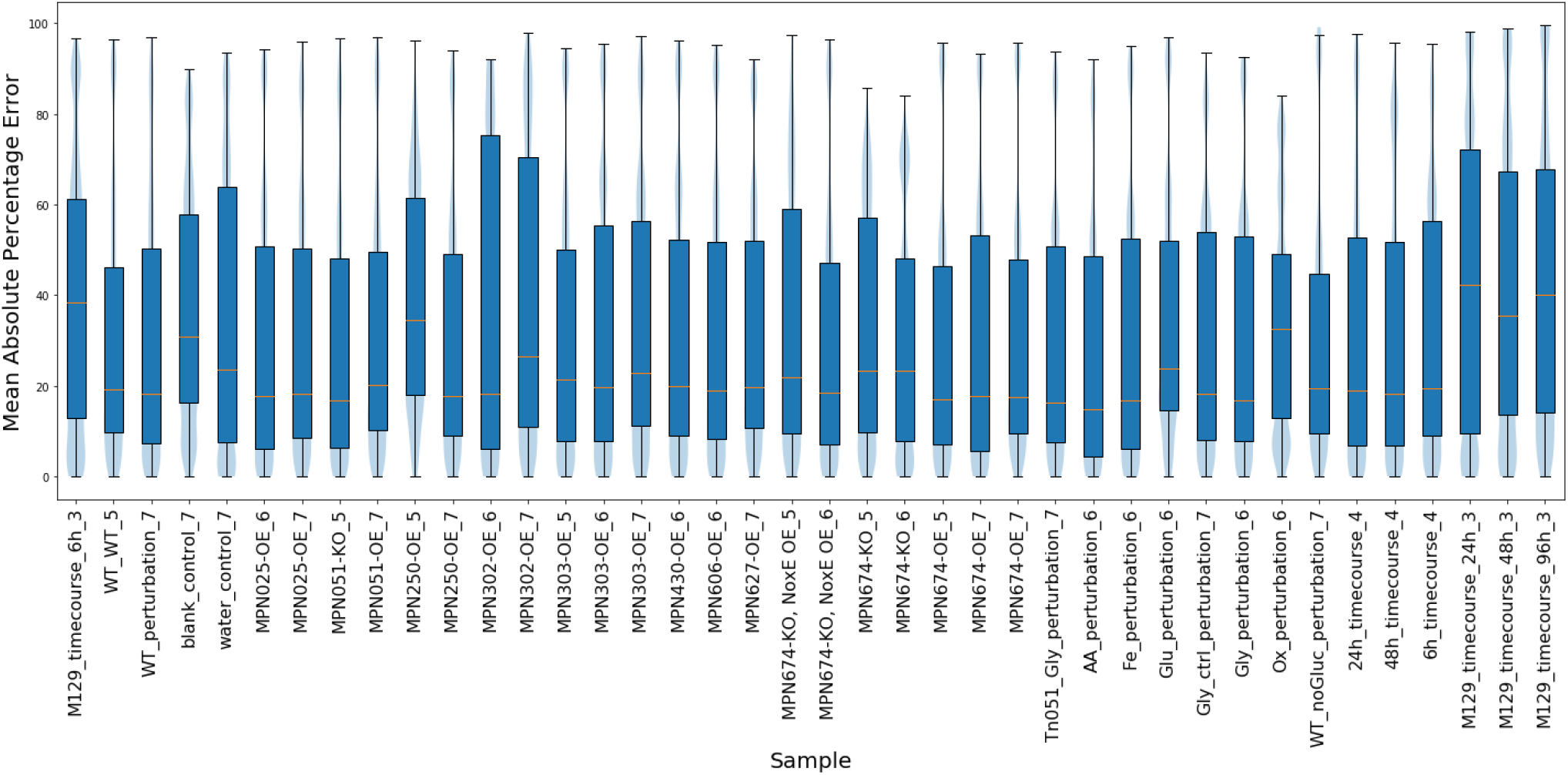
Symmetric Mean Absolute Percentage Error between simulated and measured values using a 1000x times sampling. sMAPE for all metabolites per sample are combined.

**Figure 6.**
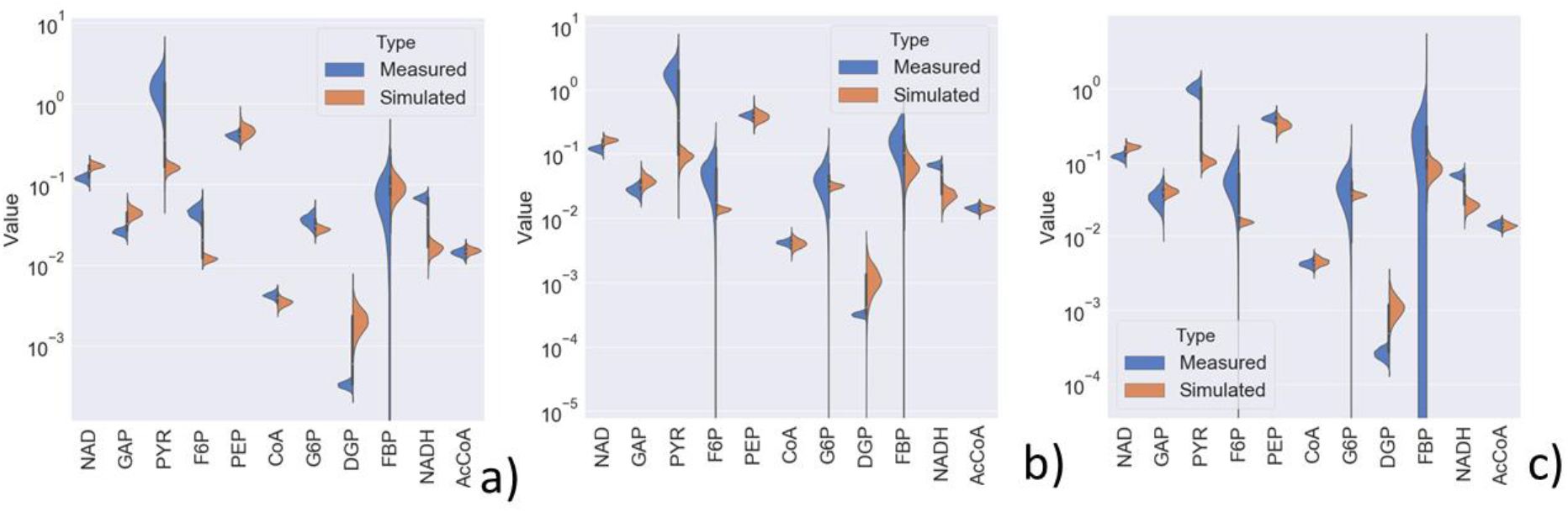
Measured and simulated steady state concentrations at a) 6h of growth, b) 24h of growth and c) 48h of growth on low glucose concentration. Measurement for DGP, NAD and NADH are reference values from samples grown on high glucose concentration.

Additionally, concentrations of for most metabolites are predicted within the 95% confidence interval of the *in-vitro* data (Supplementary material 1C). Exceptions to this rule are PYR and ATP and NADH. Simulated concentrations for ATP and NADH are systematically lower than measured values while PYR is systematically predicted to have a higher than measured concentration. Pyruvate is hard to quantify since it easily is degraded causing variation in its measurement due to for example different times between measurement and sampling. As such we cannot quantify the accuracy of the predictions for pyruvate. For the metabolites such as DGP, NAD and NADH, reference values from high glucose conditions were used to set initial concentrations. As such deviation from these reference values in the simulations in steady state was to be expected.

### Metabolic control

We used two types of metabolic control analysis to understand which enzymes and metabolites effort the greatest control on glycolysis: Local Sensitivity analysis and Global Sensitivity analysis. Control coefficients are unit less measures of the relative steady state change in a system variable, in our case the flux through PFK, in response to a relative change in a parameters value. Global sensitivity provides information on parameters that exert control independent of a specific parameter value. We achieved this by sampling parameter sets uniformly from the parameter search space. These parameters sets are not specific for *MPN* since they are not fitted to any *MPN* data. Local sensitivity analysis on the other hand is based on control coefficients derived at the steady state using parameter sets fitted to *MPN* specific data. As such the approaches are fundamentally different and complementary in the information they provide. For our local sensitivity analysis, we use the best performing10 parameter sets to calculate metabolic control coefficients (**Error! Reference source not found.**). Since these parameter sets are independent of one another, if metabolic control is higher for certain parameters based on multiple parameter sets, we can conclude the control to be relevant since it is a result of the fitting to *MPN* data. Additionally, we also performed local sensitivity analysis while sampling from the measurement distribution of metabolites to investigate the effect of measurement error on control coefficients. The steady state changes for each sampled simulation, as such we can see how the uncertainty in metabolite concentrations propagates and creates uncertainty in the control coefficients calculated at these steady states. An overview of our metabolic control analyses can be found in Supplementary file 1D. We found two main control hubs PTS_Glc + PFK and LDH+PDH+PYK. The first control hub consists of parameters associated to PTS_Glc and PFK and represents metabolism in the upper part of glycolysis, the second hub consists of parameter associated to LDH, PDH and PYK and part of pyruvate metabolism.

### Simulating combined OE and KO mutants

We used the model to simulate the combined effect of genetic perturbations targeting glycolysis enzymes (OE, KO) combined with a second perturbation, either genetic or environmental. This analysis can identify bottlenecks in central carbon metabolism consisting of combinations of enzymes. Such bottlenecks cannot be identified through local sensitivity analysis or when simulating single over expressions. The expected variations in the flux through glycolysis is shown in Figure 7. For most of these combined perturbations, only minor changes were observed. However, simulations of OE of PFK show greatly increased flux through glycolysis while oxygen stress, iron limitation and growth of MPN129 in aerated conditions lead to greatly reduced flux.

**Figure 7.**
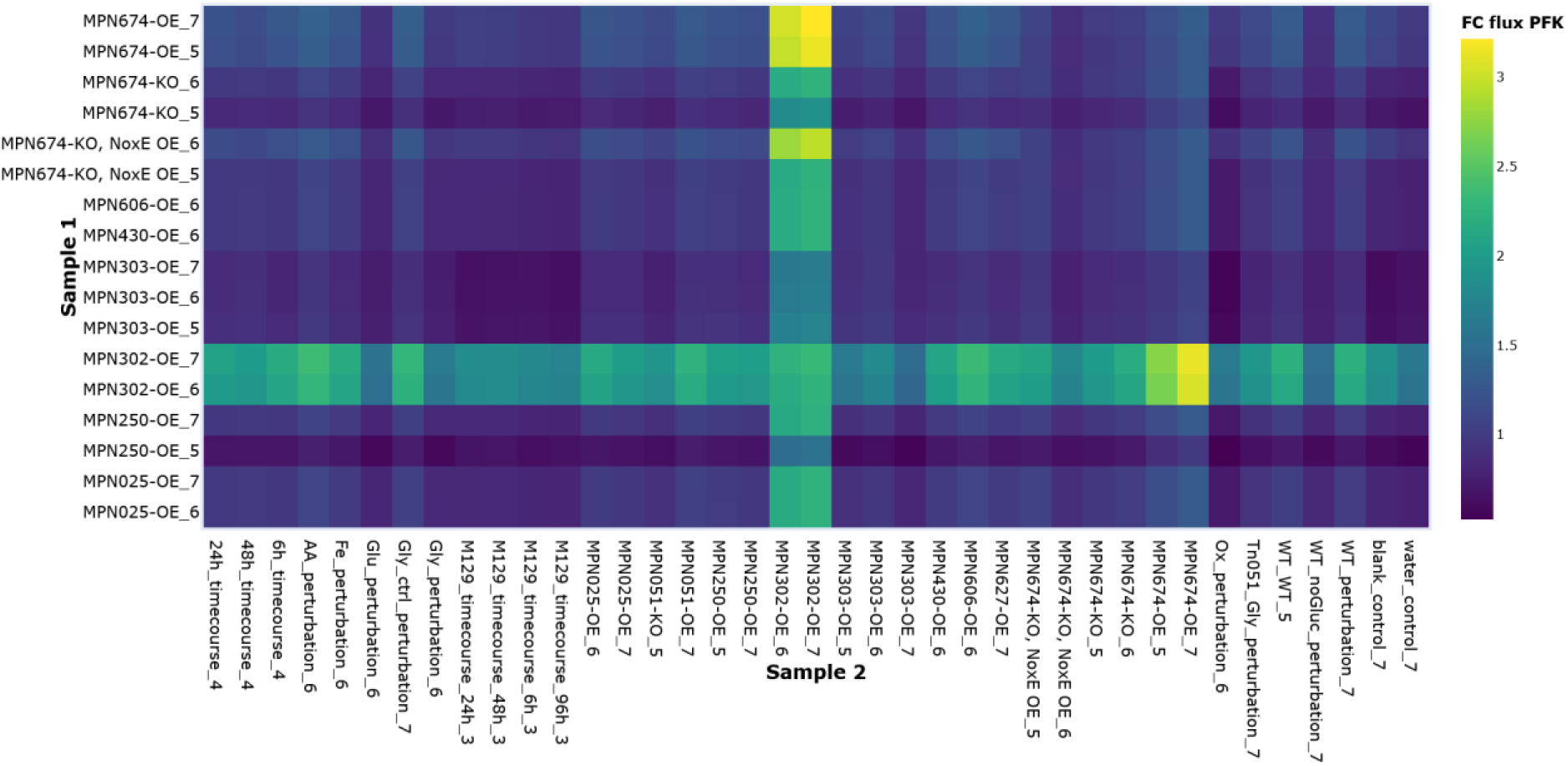
Fold change flux through PFK for combinatorial mutants and perturbations in steady state, relative to the wild type at 24h. The primary sample, of which only the protein OE values are used, are shown on the y-axis while the secondary sample, of which both changes to parameters and metabolite concentrations are used, are shown on the x-axis. The colour represents the fold change in flux through glycolysis compared to the wild type.

However, combination of PFK OE with OE of lactate dehydrogenase (LDH) or phosphotransacetylase (PTA) with acetate kinase (ACK), increase the flux nearly as much and is realistic to obtain *in vivo* since OE mutants for each of these enzymes individually are available.

Based on these simulations, combined OE mutants were suggested for lab validation: PFK+LDH, PFK+NOXE. Growth curves, protein and metabolite concentrations were measured for the PFK+LDH OE mutant but not for combined PFK+NOXE OE. Combined PFK+PDH OE did not increase flux through glycolysis. However, some positive epistasis on the growth for the combined OE of PFK+LDH was observed, with higher biomass and higher acidification than the individual mutants. None of the combined OE mutants increased the growth rate of *MPN* with respect to the wild type as could eb expected since energy metabolism is not growth limiting in *MPN*^9^.

The simulation results of the double OE mutants show that glycolysis in robust. Meaning that the network structure that includes feed forward and feed backward allosteric control, makes the glycolysis of *MPN* inherently robust.

## Discussion

Metabolomics data for a large number of perturbations, OE and KO mutants were collected. We observe that metabolite concentration of samples in general do not vary that much and that samples of similar conditions as well samples of enzymes OE and KO mutants of enzymes close to other in the metabolic network cluster together. In glycolysis we observed no clear clustering between metabolites. We also see that there is relatively little difference in the expression of glycolytic enzymes when OE enzymes in glycolysis and pyruvate metabolism. These results suggest that central carbon metabolism of *MPN* is robust against perturbation. Literature research revealed that glycolysis and pyruvate metabolism in many species is observed to be robust against perturbations ^16,55,56^. Arguably robustness is an even more important property for a minimal organism such as *MPN* where biological noise can be expected to have much larger effect than for many organisms with a larger cell volume. The property of robustness agrees with model results that show only minor changes occur even when OE multiples glycolysis enzymes in silico. We tested the effect of adding a reaction for additions of 1) ATPase, 2) O_2_ inhibition by LDH and 3) a NoxE reaction. Only the the addition of addition of a NoxE reaction resulted in a much better fit to the training data. Arguably, addition of NoxE improves parameter identifiability since the models without this reaction use a fixed reference values for NAD and NADH. The models that include NoxE allow NAD and NADH concentrations to change, this added flexibility and improves parameter identifiability since NAD and NADH associated parameter can now be used to account for differences in metabolite concentration opposed to being fixed values based on reference values. Additionally, NAD/NADH are known to have control over central carbon metabolism in other organisms such as *L. lactis* ^57,58^ therefore we argue it is likely they also have control over central carbon metabolism in *MPN*.

Steady state simulations showed the model to be flexible since it can predict metabolite concentrations well for all mutant samples. Part of this flexibility is the result of including parameters representing the enzyme concentration, therefore genetic perturbations are accounted for in simulations. Similarly, by using measurements of cofactors from these conditions, the model can approximate the effects of these simulations on central carbon metabolism. Additionally, we argue the flexibility of the model might partly be the result of the inherent robustness of central carbon metabolism in *MPN*.

## Conclusion

In this study we integrated experimental data and model simulations and analysis to explore the robustness of central carbon metabolism of *MPN* in steady state.

Firstly, we analyzed metabolomics data. We observed that samples from similar conditions as well as samples of OE and KO mutants of enzymes close to each other in the metabolic network cluster together. Samples from vastly varying conditions, such as aerated conditions, do not cluster with the other samples. Secondly, we build a model to simulate samples of various single or combined perturbations. The simple model presented in this study can predict metabolite concentration with reasonable accuracy for a wide range of conditions and OE and KO mutants.

Two control hubs were identified using our dynamic model a) upper glycolysis (PTS_Glc + PFK) and b) lower pyruvate metabolism (LDH+PDH+PYK). No single or combined OE mutant of glycolysis and pyruvate metabolism enzymes resulted in a higher growth rate although OE of PFK and LDH resulted in somewhat higher acidification indicating there might be higher flux through glycolysis. These results are in in agreement with studies that show that glycolysis and pyruvate metabolism in *MPN* is not growth limiting. Both the results from the analysis of our samples as well as the model results, suggest robustness to be a central property of *MPN* glycolysis and pyruvate metabolism.

## Supporting information

Supplementary Table 1 - equations

## Abbreviation list

AcCoA: Acetyl coenzyme A
ACE: Acetate
ACK: Acetate kinase
ADP: Adenosine diphosphate
ATP: Adenosine triphosphate
ATPase: Adenylpyrophosphatase
CoA: Coenzyme A dehydrogenase
DGP: Diacylglycerol phosphate
ENO: Enolase
F6P: Fructose 6-phosphate
FBA: Fructose-bisphosphate aldolase
FBP: Fructose-1,6-bisphosphatase
G6P: Glucose 6-phosphate
GAP: Glyceraldehyde-3-phosphate
GAPDH: Glyceraldehyde 3-phosphate
GLC_Ext: Extracellular glucose
GMP: Glycerate phosphomutase
Kcat: Enzyme catalytic rate
Keq: Equilibrium constant
Km: Michaelis-Menten constant
KO: Knock out
LAC: Lactate
LDH: Lactate dehydrogenase
*MPN*: Mycoplasma pneumonia
NAD: Nicotinamide adenine dinucleotide
NADH: Reduced nicotinamide adenine dinucleotide
OE: Over-expression
PDH: Pyruvate dehydrogenase
PEP: Phosphoenolpyruvate
PFK: Phosphofructokinase
PGI: Phosphoglucose isomerase
PGK: Phosphoglycerate kinase
Pi_Int: Orthophosphate
PTA: Phosphotransacetylase
PTS_Glc: Phosphotransferase system
PYK: Pyruvate kinase
PYR: Pyruvate

## Acknowledgement

We acknowledge support of the Spanish Ministry of Economy, Industry and Competitiveness (MEIC) to the EMBL partnership, the Spanish Ministry of Economy and Competitiveness, ‘Centro de Excelencia Severo Ochoa 2013-2017’, the European Research Council (ERC) under the European Union’s Horizon 2020 research and innovation program under agreement No 670216 (MYCOCHASSIS), the European Union’s Horizon 2020 research and innovation programme under grant agreement No 634942 (MycoSynVac), the CERCA Programme / Generalitat de Catalunya, FEDER project from Instituto Carlos III (ISCIII, Acción Estratégica en Salud 2016) (reference CP16/00094) and “Secretaria d’Universitats i Recerca del Departament d’Economia i Coneixement de la Generalitat de Catalunya” (2014SGR678). We would like to thank dr.ir. A. Rinzema for his expert advice on calculating the various oxygen diffusion rates in our experimental setup.

## Contributions

LS, VMdS and MS-D conceived the study. NZ developed the model and performed the simulations with guidance and input from PS, VMdS, LS and MS-D. EY, DS, SM and CG performed the experiments with guidance and input from ML-S and LS. NZ wrote the manuscript with input from VMdS, PS, ML-S, LS and MS-D.

