## Supplementary Table 1 - equations for "Exploring the adaptability and robustness of the central carbon metabolism of *Mycoplasma pneumoniae*"

#### Supplementary file 1 A: Model equations

| **Reaction nr** | **reaction** | **Function** | **Present in model(s)** |
|---|---|---|---|
| 1 PTS_Glc Glucose proton symporter | GLC_Ext + PEP -> G6P + PYR; FBP Pi_Int | $\frac{PTS\_Glc \cdot \frac{Pi\_Int}{Pi\_Int + kaPi\_Int\_PTS\_Glc} \cdot \frac{kiFBP\_PTS\_Glc}{FBP + kiFBP\_PTS\_Glc} \cdot kcat\_PTS\_Glc \cdot \frac{GLC\_Ext}{kmGLC\_PTS\_Glc} \cdot \frac{PEP}{kmPEP\_PTS\_Glc}}{\left(1 + \frac{GLC\_Ext}{kmGLC\_PTS\_Glc}\right) \cdot \left(1 + \frac{PEP}{kmPEP\_PTS\_Glc}\right) + \left(1 + \frac{G6P}{kmG6P\_PTS\_Glc}\right) \cdot \left(1 + \frac{PYR}{kmPYR\_PTS\_Glc}\right) - 1}$ | 1,2,3,4,5,6 |
| 2 PGI Glucose-6-phosphate isomerase | G6P = F6P | $\frac{PGI \cdot \left(kcat\_PGI \cdot \frac{G6P}{kmG6P\_PGI} - \frac{kcat\_PGI}{Keq\_PGI} \cdot \frac{F6P}{kmG6P\_PGI}\right)}{1 + \frac{G6P}{kmG6P\_PGI} + \frac{F6P}{kmF6P\_PGI}}$ | 1,2,3,4,5,6 |
| 3 PFK Phosphofructokinase | F6P + ATP -> FBP + ADP | $\frac{PFK \cdot kcat\_PFK \cdot \frac{ATP}{kmF6P\_PFK} \cdot \frac{F6P}{kmATP\_PFK}}{\left(1 + \frac{ATP}{kmF6P\_PFK}\right) \cdot \left(1 + \frac{F6P}{kmATP\_PFK}\right) + \left(1 + \frac{ADP}{kmFBP\_PFK}\right) \cdot \left(1 + \frac{FBP}{kmADP\_PFK}\right) - 1}$ | 1,2,3,4,5,6 |
| 4 FBA Fructose-bisphosphate aldolase | GAP + Pi_Int + NAD = DGP + NADH | $\frac{FBA \cdot \left(kcat\_FBA \cdot \frac{FBP}{kmFBP\_FBA} - \frac{kcat\_FBA}{Keq\_FBA} \cdot \frac{GAP^2}{kmFBP\_FBA}\right)}{1 + \frac{FBP}{kmFBP\_FBA} + \frac{GAP}{kmGAP\_FBA} + \left(\frac{GAP}{kmGAP\_FBA}\right)^2}$ | 1,2,3,4,5,6 |
| 5 GAPDH Glyceraldehyde-3-phosphate dehydrogenase | DGP + ADP = PEP + ATP | $\frac{GAPDH \cdot \left(kcat\_GAPDH \cdot \frac{GAP}{kmGAP\_GAPDH} \cdot \frac{NAD}{kmNAD\_GAPDH} \cdot \frac{Pi\_Int}{kmPi\_Int\_GAPDH} - \frac{kcat\_GAPDH}{Keq\_GAPDH} \cdot \frac{DGP}{kmGAP\_GAPDH} \cdot \frac{NADH}{kmNAD\_GAPDH}\right.}{\left(1 + \frac{GAP}{kmGAP\_GAPDH}\right) \cdot \left(1 + \frac{Pi\_Int}{kmPi\_Int\_GAPDH}\right) \cdot \left(1 + \frac{NAD}{kmNAD\_GAPDH}\right) + \left(1 + \frac{DGP}{kmDGP\_GAPDH}\right) \cdot \left(1 + \frac{NADH}{kmNADH\_GAP[illegible]}\right.}$ | 1,2,3,4,5,6 |

| | | | |
|---|---|---|---|
| 6 ENO<br>Enolase | PEP + ADP -> PYR + ATP; FBP Pi_Int | $$\frac{\mathrm{ENO}\cdot\left(\mathrm{kcat\_ENO}\cdot\frac{\mathrm{DGP}}{\mathrm{kmDGP\_ENO}}\cdot\frac{\mathrm{ADP}}{\mathrm{kmADP\_ENO}}-\frac{\mathrm{kcat\_ENO}}{\mathrm{Keq\_ENO}}\cdot\frac{\mathrm{PEP}}{\mathrm{kmDGP\_ENO}}\cdot\frac{\mathrm{ATP}}{\mathrm{kmADP\_ENO}}\right)}{\left(1+\frac{\mathrm{DGP}}{\mathrm{kmDGP\_ENO}}\right)\cdot\left(1+\frac{\mathrm{ADP}}{\mathrm{kmADP\_ENO}}\right)+\left(1+\frac{\mathrm{PEP}}{\mathrm{kmPEP\_ENO}}\right)\cdot\left(1+\frac{\mathrm{ATP}}{\mathrm{kmATP\_ENO}}\right)-1}$$ | 1,2,3,4,5,6 |
| 7 PYK<br>Pyruvate kinase | PYR + CoA + NAD = AcCoA + NADH; GAP | $$\frac{\frac{\mathrm{PYK}\cdot\mathrm{FBP}}{\mathrm{FBP}+\mathrm{kaFBP\_PYK}}\cdot\frac{\mathrm{kiPi\_Int\_PYK}^{\mathrm{nPYK}}}{\mathrm{Pi\_Int}^{\mathrm{nPYK}}+\mathrm{kiPi\_Int\_PYK}^{\mathrm{nPYK}}}\cdot\mathrm{kcat\_PYK}\cdot\frac{\mathrm{ADP}}{\mathrm{kmADP\_PYK}}\cdot\frac{\mathrm{PEP}}{\mathrm{kmPEP\_PYK}}}{\left(1+\frac{\mathrm{ADP}}{\mathrm{kmADP\_PYK}}\right)\cdot\left(1+\frac{\mathrm{PEP}}{\mathrm{kmPEP\_PYK}}\right)+\left(1+\frac{\mathrm{ATP}}{\mathrm{kmATP\_PYK}}\right)\cdot\left(1+\frac{\mathrm{PYR}}{\mathrm{kmPYR\_PYK}}\right)-1}$$ | 1,2,3,4,5,6 |
| 8 LDH<br>Lactate dehydrogenase | PYR + NADH -> LAC + NAD; FBP Pi_Int* O2 | $$\frac{\frac{\mathrm{LDH}\cdot\mathrm{FBP}}{\mathrm{FBP}+\mathrm{kaFBP\_LDH}}\cdot\frac{\mathrm{kiPi\_Int\_LDH}}{\mathrm{Pi\_Int}+\mathrm{kiPi\_Int\_LDH}}\cdot\frac{\mathrm{kiO2\_LDH}}{\mathrm{O2}+\mathrm{kiO2\_LDH}}\cdot\mathrm{kcat\_LDH}\cdot\frac{\mathrm{NADH}}{\mathrm{kmPYR\_LDH}}\cdot\frac{\mathrm{PYR}}{\mathrm{kmNADH\_LDH}}}{\left(1+\frac{\mathrm{NADH}}{\mathrm{kmPYR\_LDH}}\right)\cdot\left(1+\frac{\mathrm{PYR}}{\mathrm{kmNADH\_LDH}}\right)+\left(1+\frac{\mathrm{LAC}}{\mathrm{kmLAC\_LDH}}\right)\cdot\left(1+\frac{\mathrm{NAD}}{\mathrm{kmNAD\_LDH}}\right)-1}$$ | 1*,2*,3*,4*,5, 6 |
| 9 PDH<br>Pyruvate dehydrogenase | PYR + CoA + NAD = AcCoA + NADH; GAP | $$\frac{\mathrm{PDH}\cdot\frac{\mathrm{kiGAP\_PDH}}{\mathrm{GAP}+\mathrm{kiGAP\_PDH}}\cdot\left(\mathrm{kcat\_PDH}\cdot\frac{\mathrm{NAD}}{\mathrm{kmPYR\_PDH}}\cdot\frac{\mathrm{PYR}}{\mathrm{kmCoA\_PDH}}\cdot\frac{\mathrm{CoA}}{\mathrm{kmNAD\_PDH}}-\frac{\mathrm{kcat\_PDH}}{\mathrm{Keq\_PDH}}\cdot\frac{\mathrm{AcCoA}}{\mathrm{kmPYR\_PDH}}\cdot\frac{\mathrm{NADH}}{\mathrm{kmCoA\_PDH}}\right)}{\left(1+\frac{\mathrm{NAD}}{\mathrm{kmPYR\_PDH}}\right)\cdot\left(1+\frac{\mathrm{PYR}}{\mathrm{kmCoA\_PDH}}\right)\cdot\left(1+\frac{\mathrm{CoA}}{\mathrm{kmNAD\_PDH}}\right)+\left(1+\frac{\mathrm{AcCoA}}{\mathrm{kmAcCoA\_PDH}}\right)\cdot\left(1+\frac{\mathrm{NADH}}{\mathrm{kmNADH\_PDH}}\right)-1}$$ | 1,2,3,4,5,6 |
| 10 PTA_ACK<br>Phosphotransacetylase | AcCoA + ADP -> ACE + ATP + CoA | $$\frac{\mathrm{PTA\_ACK}\cdot\mathrm{kcat\_PTA\_ACK}\cdot\frac{\mathrm{AcCoA}}{\mathrm{kmAcCoA\_PTA\_ACK}}\cdot\frac{\mathrm{ADP}}{\mathrm{kmADP\_PTA\_ACK}}}{\left(1+\frac{\mathrm{AcCoA}}{\mathrm{kmAcCoA\_PTA\_ACK}}\right)\cdot\left(1+\frac{\mathrm{ADP}}{\mathrm{kmADP\_PTA\_ACK}}\right)+\left(1+\frac{\mathrm{ACE}}{\mathrm{kmACE\_PTA\_ACK}}\right)\cdot\left(1+\frac{\mathrm{ATP}}{\mathrm{kmATP\_PTA\_ACK}}\right)\cdot(1+\frac{\mathrm{CoA}}{\mathrm{kmCoA\_PTA\_ACK}}}$$ | 1,2,3,4,5,6 |
| 11 ATPase | ATP -> ADP + Pi_Int | $$\mathrm{ATPase}\cdot\mathrm{kcat\_ATPase}\cdot\frac{\left(\frac{\mathrm{ATP}}{\mathrm{kmATP\_ATPase}}\right)^{\mathrm{nATPase}}}{\left(\frac{\mathrm{ATP}}{\mathrm{kmATP\_ATPase}}\right)^{\mathrm{nATPase}}+1}$$ | 2,4,6 |
| 12 NOXE<br>NADH oxidase | O2 + 2 * NADH -> 2 * NAD | $$\frac{\mathrm{NOXE}\cdot\mathrm{kcat\_NOXE}\cdot\left(\frac{\mathrm{O2}}{\mathrm{kmNADH\_NOXE}}\right)^{2}\cdot\frac{\mathrm{NADH}}{\mathrm{kmO2\_NOXE}}}{\left(1+\frac{\mathrm{O2}}{\mathrm{kmNADH\_NOXE}}\right)^{2}\cdot\left(1+\frac{\mathrm{NADH}}{\mathrm{kmO2\_NOXE}}\right)+1+\frac{\mathrm{NAD}}{\mathrm{kmNAD\_NOXE}}-1}$$ | 3,4,6 |

*Models marked with an asterisk contain the function for Lactate Dehydrogenase without the oxygen inhibition term $\frac{\text{kiO2\_LHD}}{\text{O2+kiO2\_LDH}}$

### Supplementary file 1 B comparison of model additions

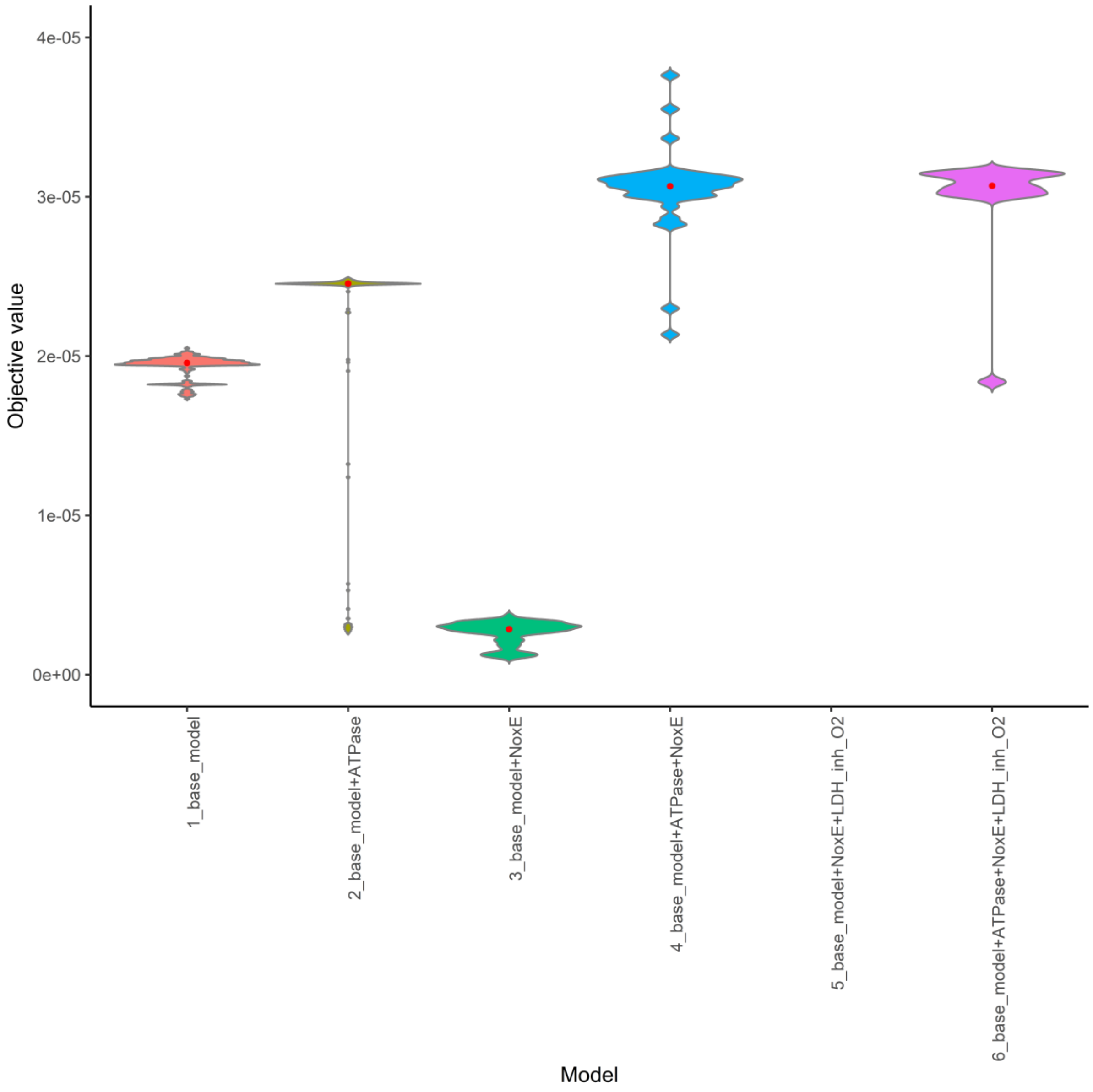


*Figure 1 Zoomed in version of comparison of 100 parameter sets for the model with and without additions. Outliers on the top were removed to better show overall performance of the models.*

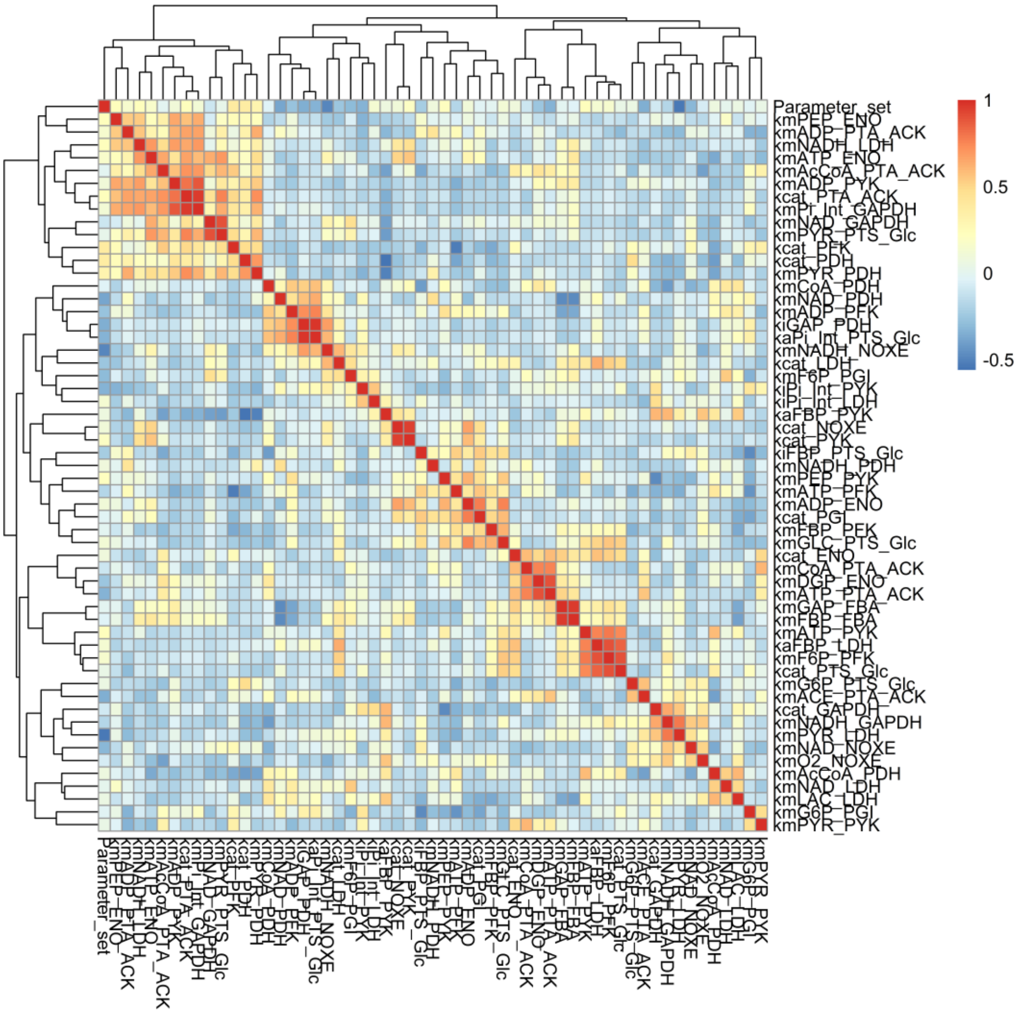


*Figure 2 Correlation of the parameter sets for model 3 that includes NoxE for the best 10 performing parameter sets.*

Parameter distributions for each of the 6 models over 100 parameter sets as well as heatmaps showing the correlation between parameters over all 100 fittings for each of the six model configurations are available upon request.

#### Supplementary file 1 C Simulating 40 independent samples.

Metabolite concentrations measured and predicted using a 1000x sampling.

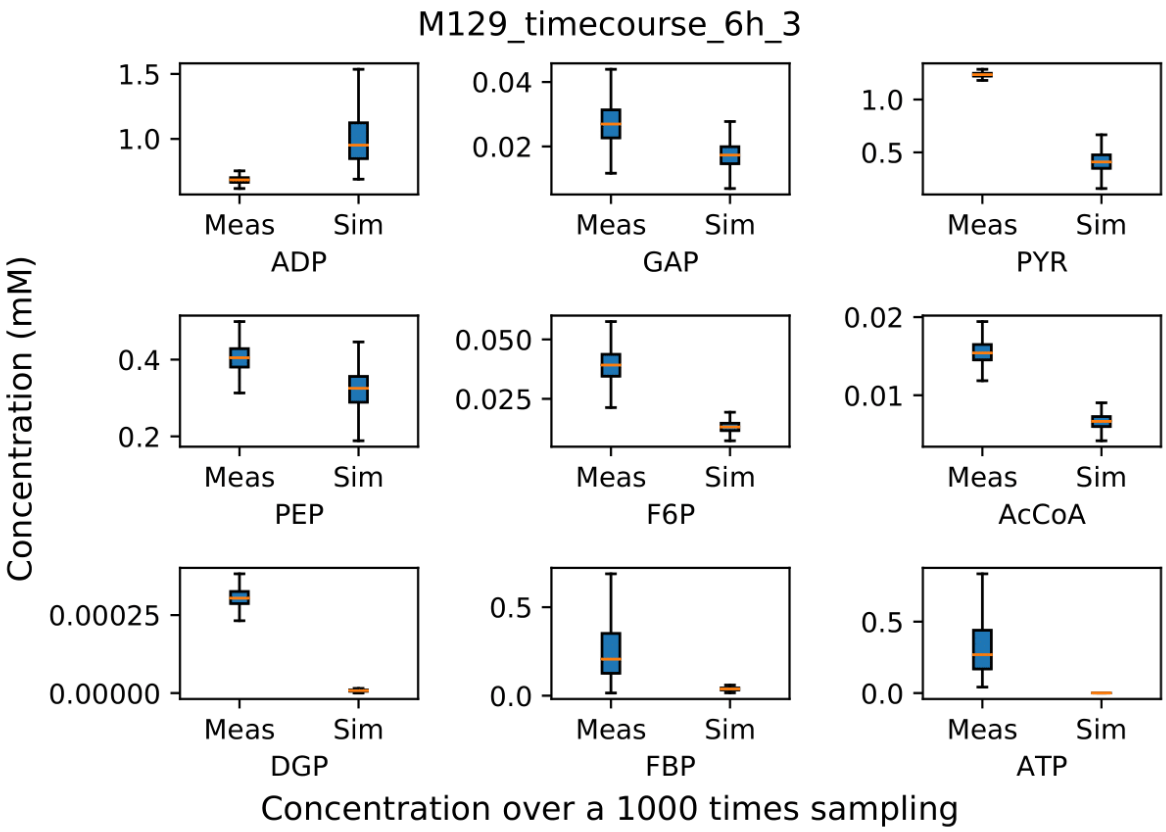


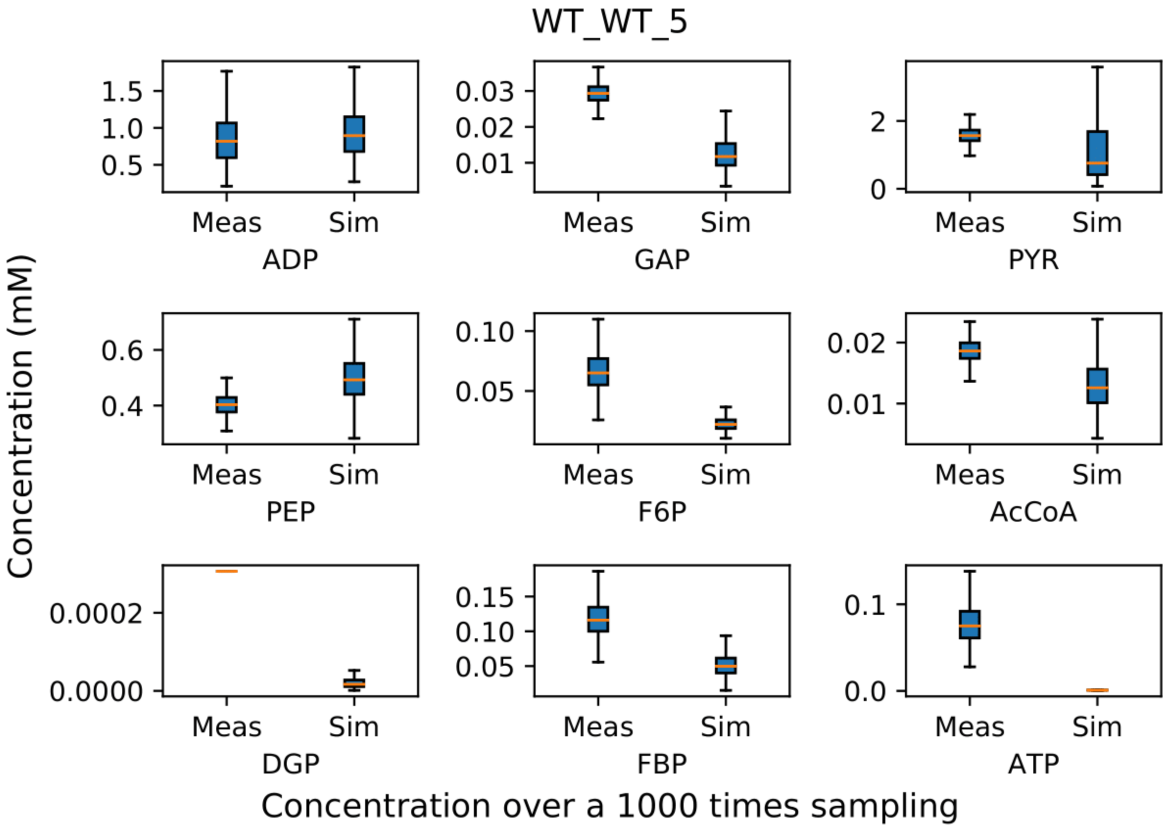

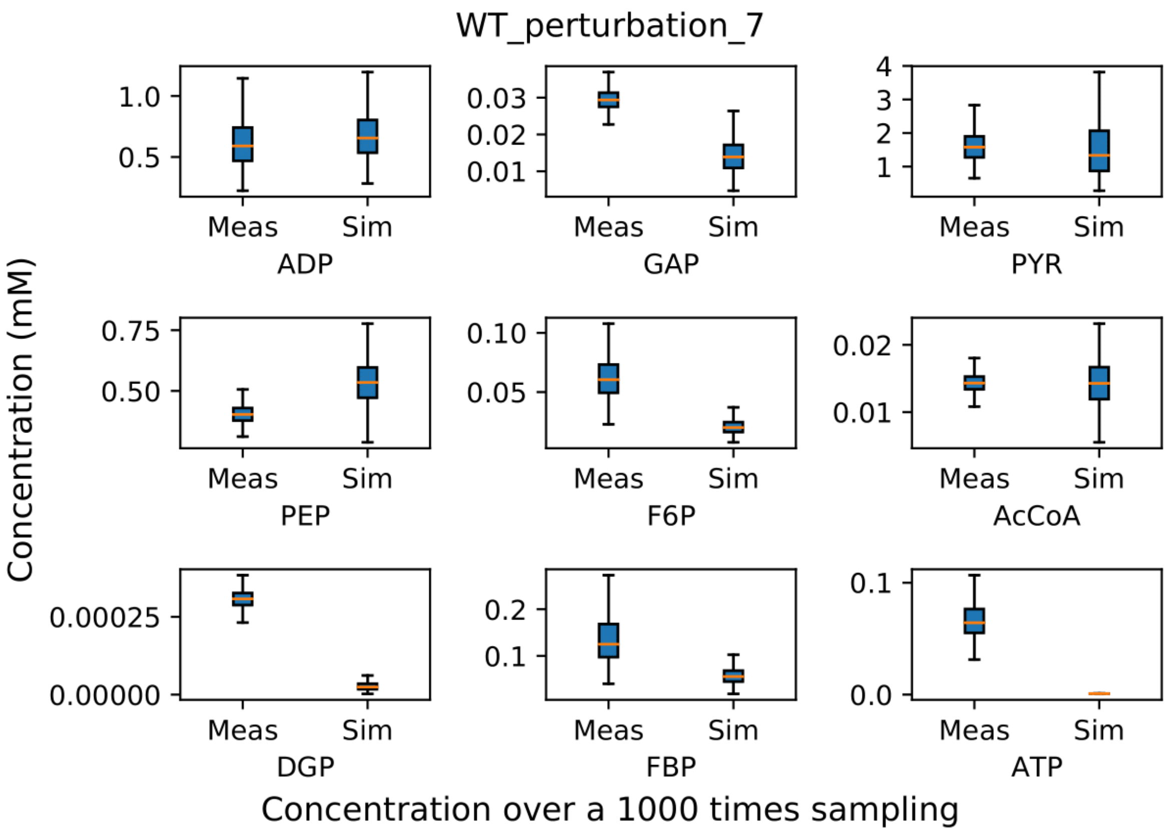
WT_perturbation_7
Concentration (mM)
ADP
GAP
PYR
PEP
F6P
AcCoA
DGP
FBP
ATP
Meas
Sim
Concentration over a 1000 times sampling

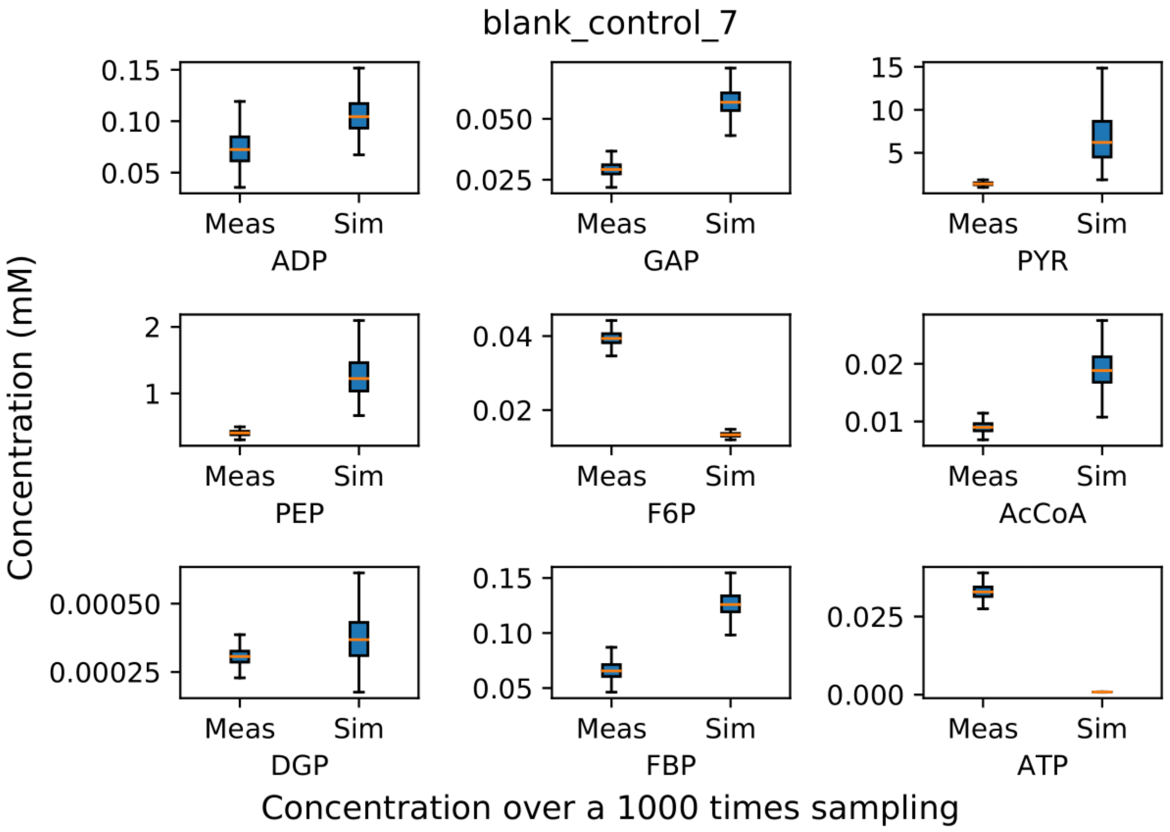
blank_control_7
Concentration (mM)
ADP
GAP
PYR
PEP
F6P
AcCoA
DGP
FBP
ATP
Meas
Sim
Concentration over a 1000 times sampling

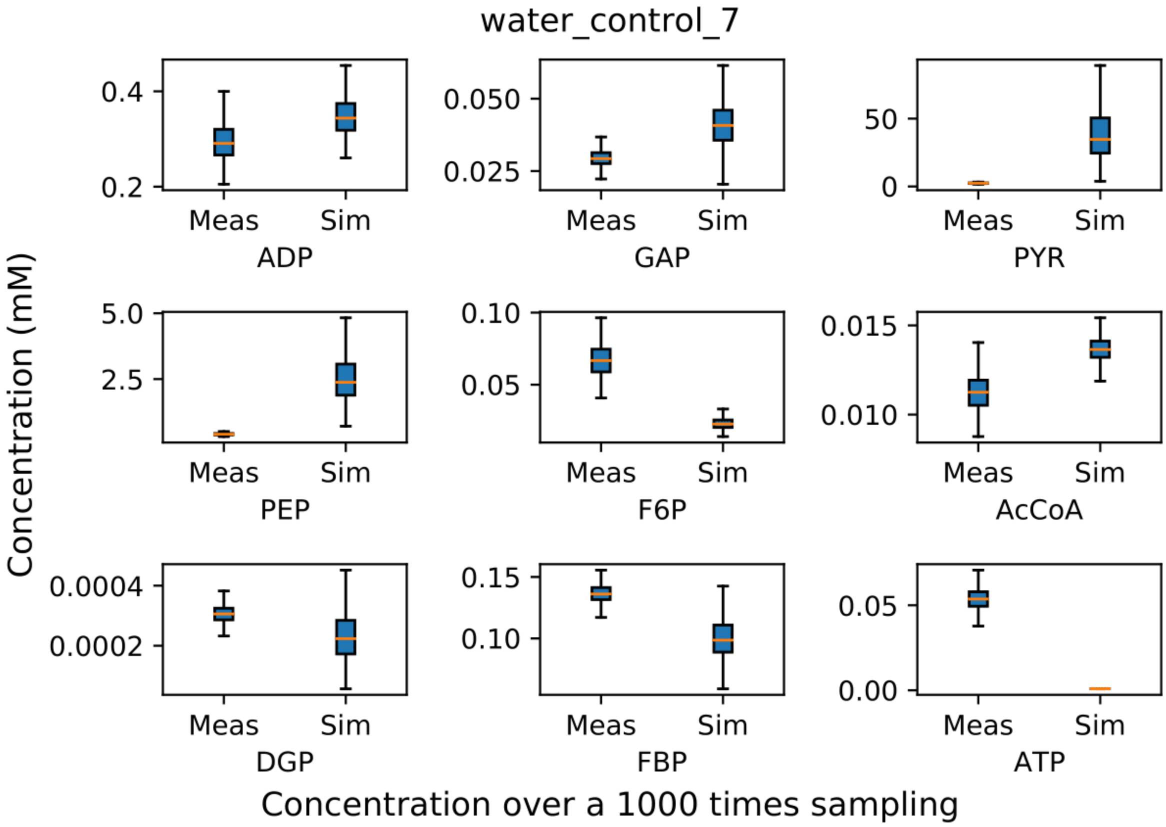
water_control_7
Concentration (mM)
0.4
0.2
Meas
Sim
ADP
0.050
0.025
Meas
Sim
GAP
50
0
Meas
Sim
PYR
5.0
2.5
Meas
Sim
PEP
0.10
0.05
Meas
Sim
F6P
0.015
0.010
Meas
Sim
AcCoA
0.0004
0.0002
Meas
Sim
DGP
0.15
0.10
Meas
Sim
FBP
0.05
0.00
Meas
Sim
ATP
Concentration over a 1000 times sampling

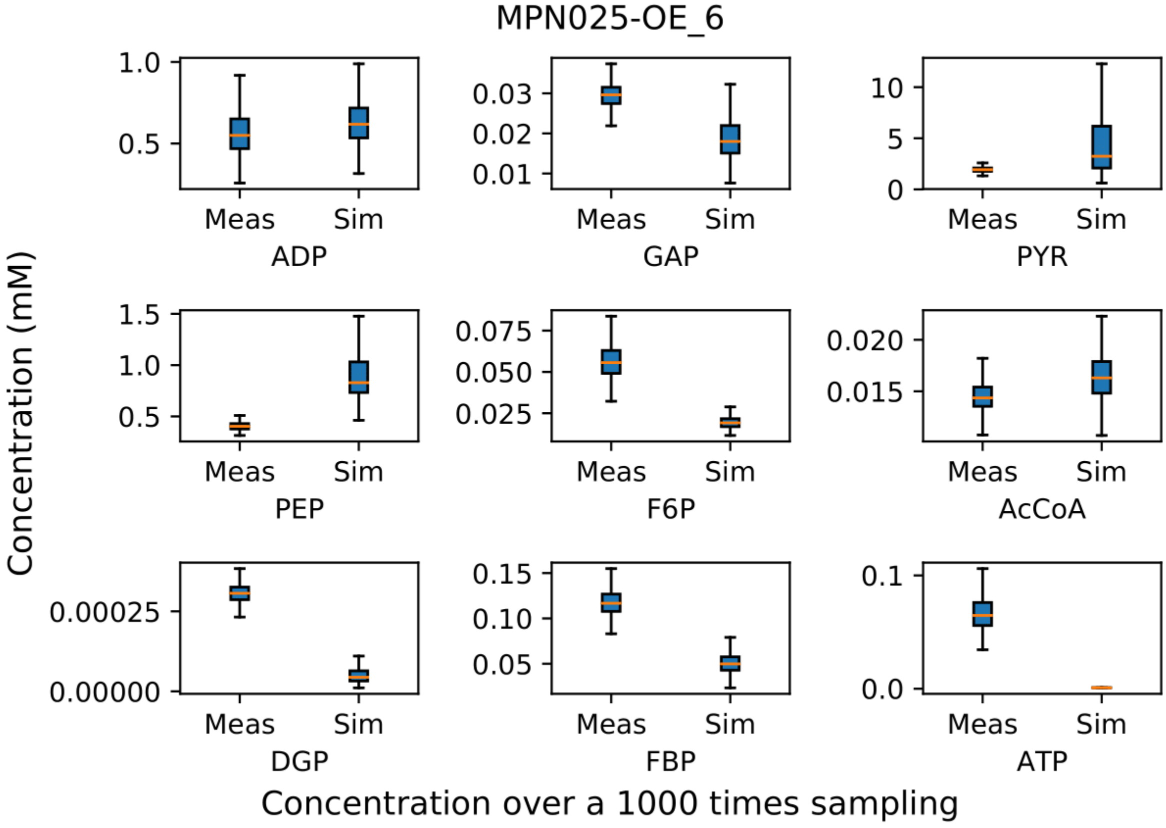
MPN025-OE_6
Concentration (mM)
1.0
0.5
Meas
Sim
ADP
0.03
0.02
0.01
Meas
Sim
GAP
10
5
0
Meas
Sim
PYR
1.5
1.0
0.5
Meas
Sim
PEP
0.075
0.050
0.025
Meas
Sim
F6P
0.020
0.015
Meas
Sim
AcCoA
0.00025
0.00000
Meas
Sim
DGP
0.15
0.10
0.05
Meas
Sim
FBP
0.1
0.0
Meas
Sim
ATP
Concentration over a 1000 times sampling

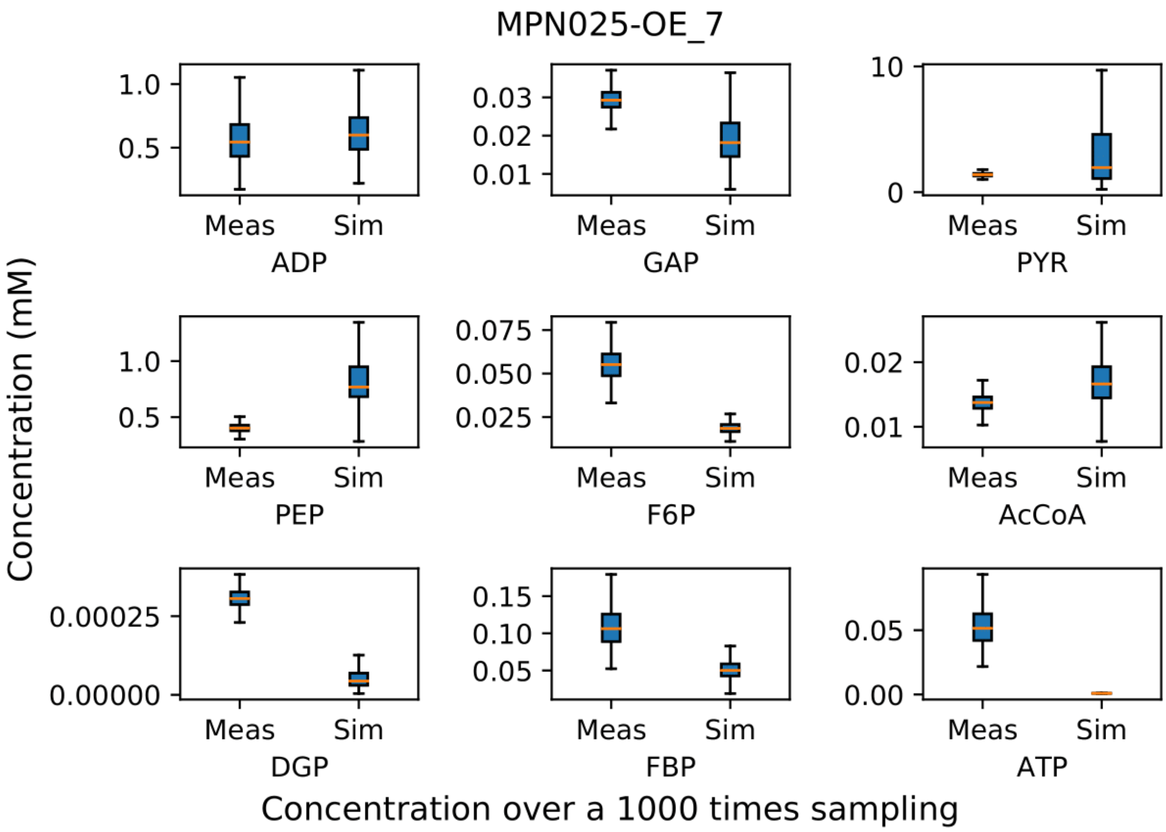
MPN025-OE_7
Concentration (mM)
ADP
GAP
PYR
PEP
F6P
AcCoA
DGP
FBP
ATP
Meas
Sim
Concentration over a 1000 times sampling

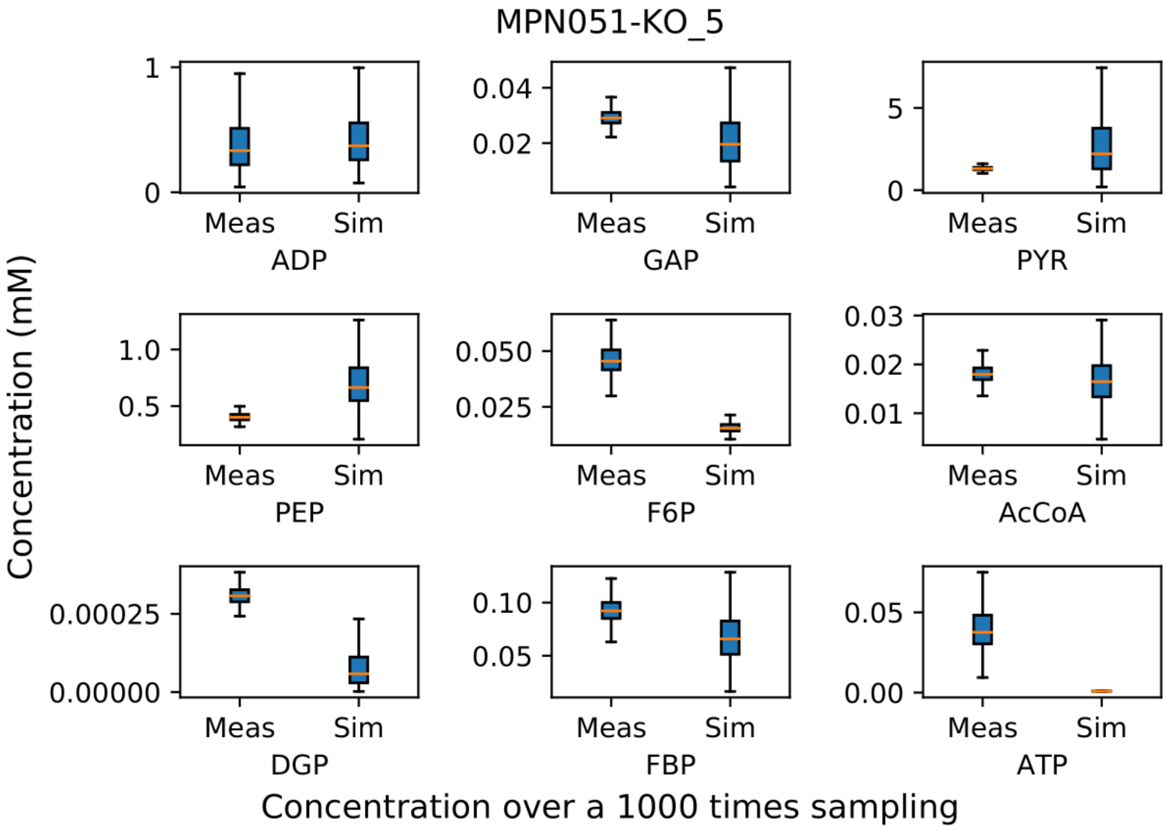
MPN051-KO_5
Concentration (mM)
ADP
GAP
PYR
PEP
F6P
AcCoA
DGP
FBP
ATP
Meas
Sim
Concentration over a 1000 times sampling

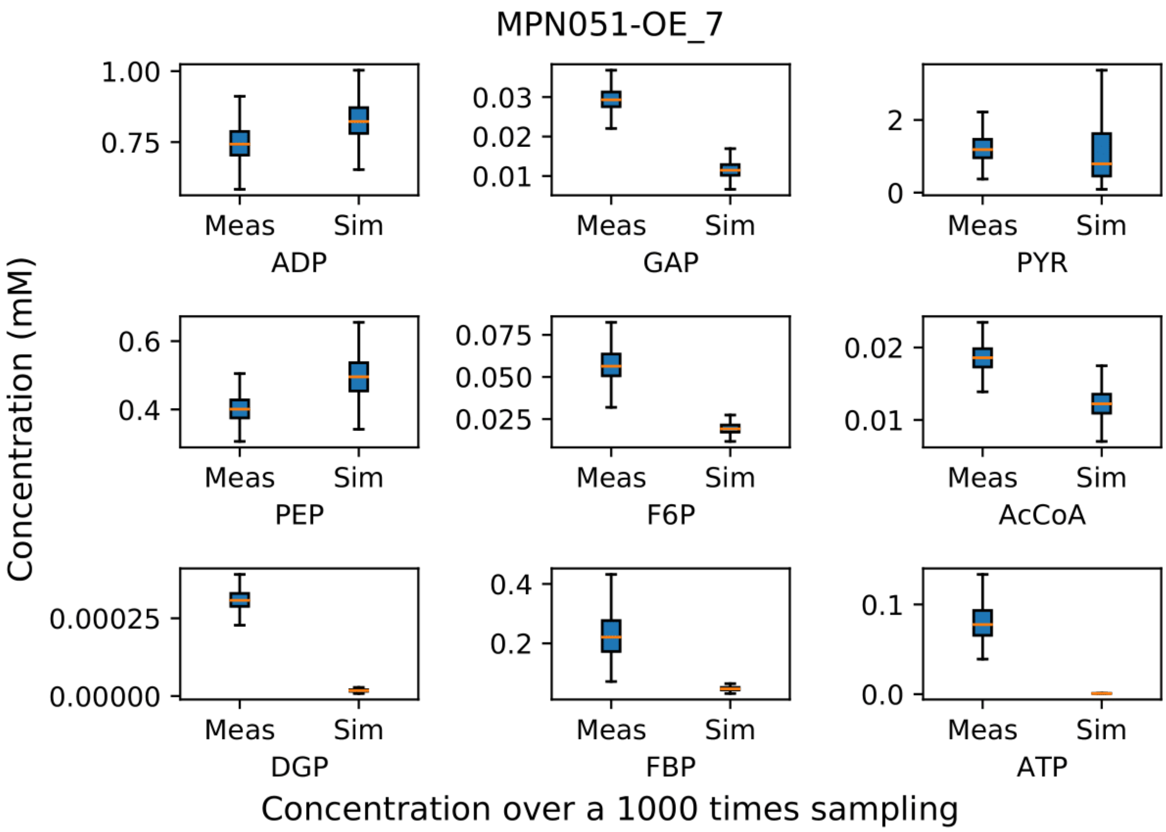
MPN051-OE_7
Concentration (mM)
ADP
GAP
PYR
PEP
F6P
AcCoA
DGP
FBP
ATP
Meas
Sim
Concentration over a 1000 times sampling

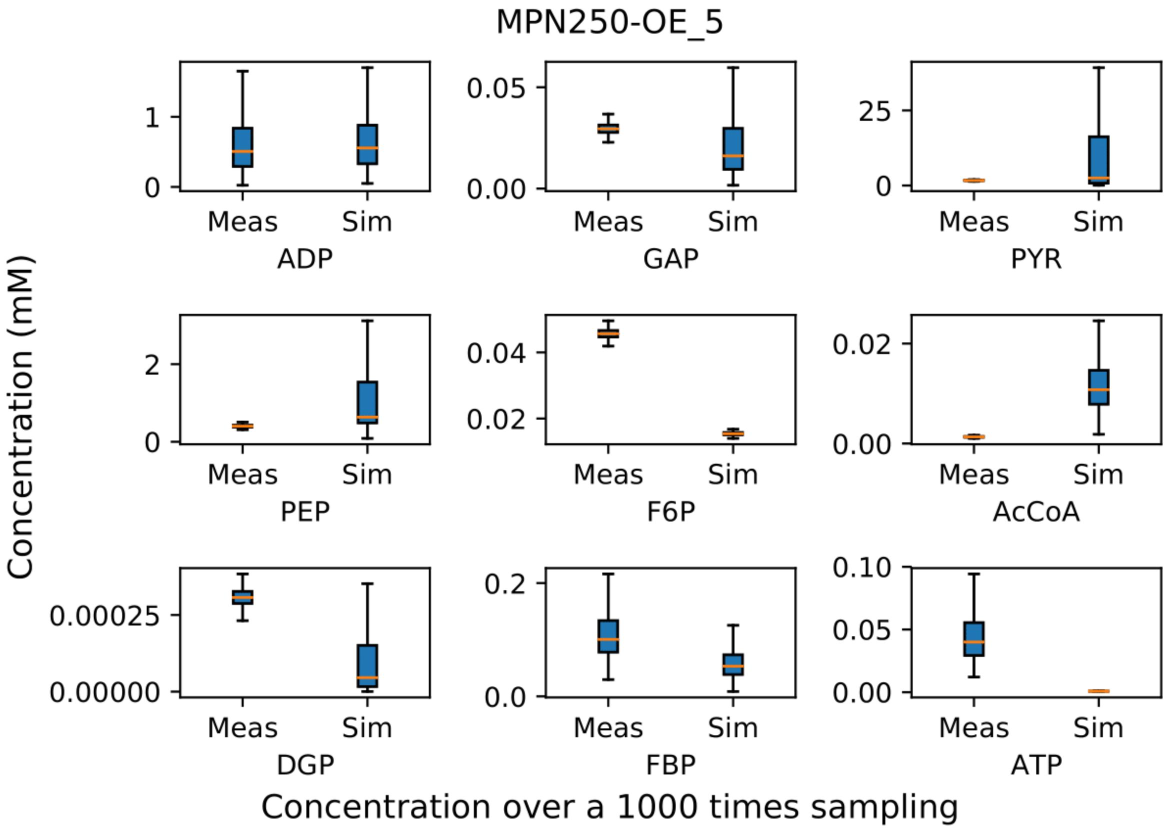
MPN250-OE_5
Concentration (mM)
ADP
GAP
PYR
PEP
F6P
AcCoA
DGP
FBP
ATP
Meas
Sim
Concentration over a 1000 times sampling

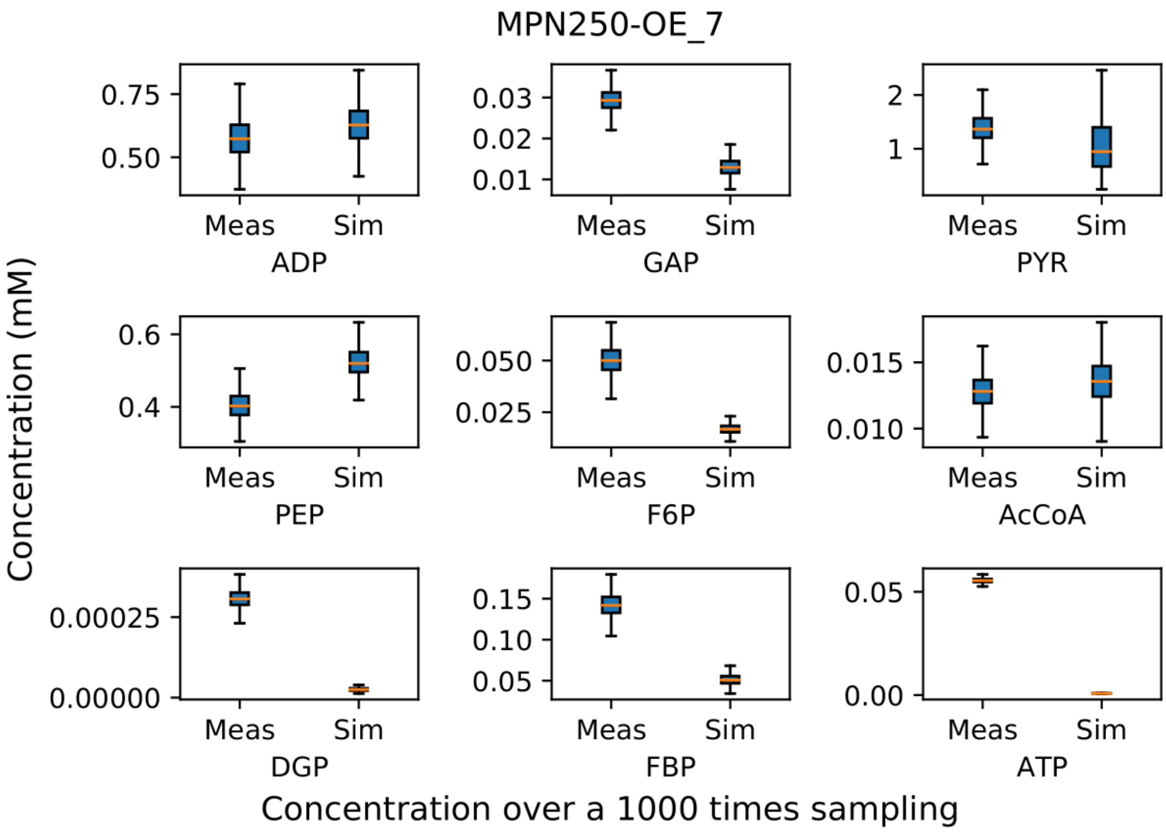
MPN250-OE_7
Concentration (mM)
0.75
0.50
Meas
Sim
ADP
0.03
0.02
0.01
GAP
2
1
PYR
0.6
0.4
PEP
0.050
0.025
F6P
0.015
0.010
AcCoA
0.00025
0.00000
DGP
0.15
0.10
0.05
FBP
0.05
0.00
ATP
Concentration over a 1000 times sampling

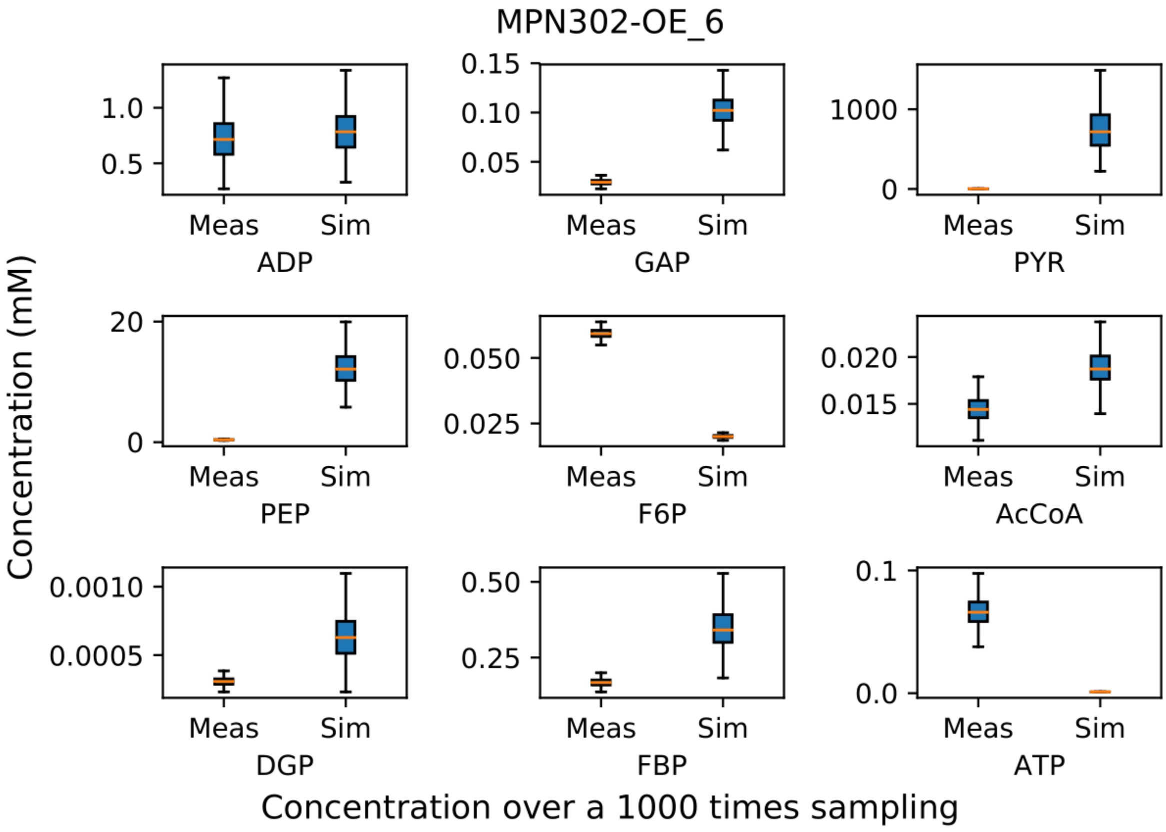
MPN302-OE_6
Concentration (mM)
1.0
0.5
Meas
Sim
ADP
0.15
0.10
0.05
GAP
1000
0
PYR
20
0
PEP
0.050
0.025
F6P
0.020
0.015
AcCoA
0.0010
0.0005
DGP
0.50
0.25
FBP
0.1
0.0
ATP
Concentration over a 1000 times sampling

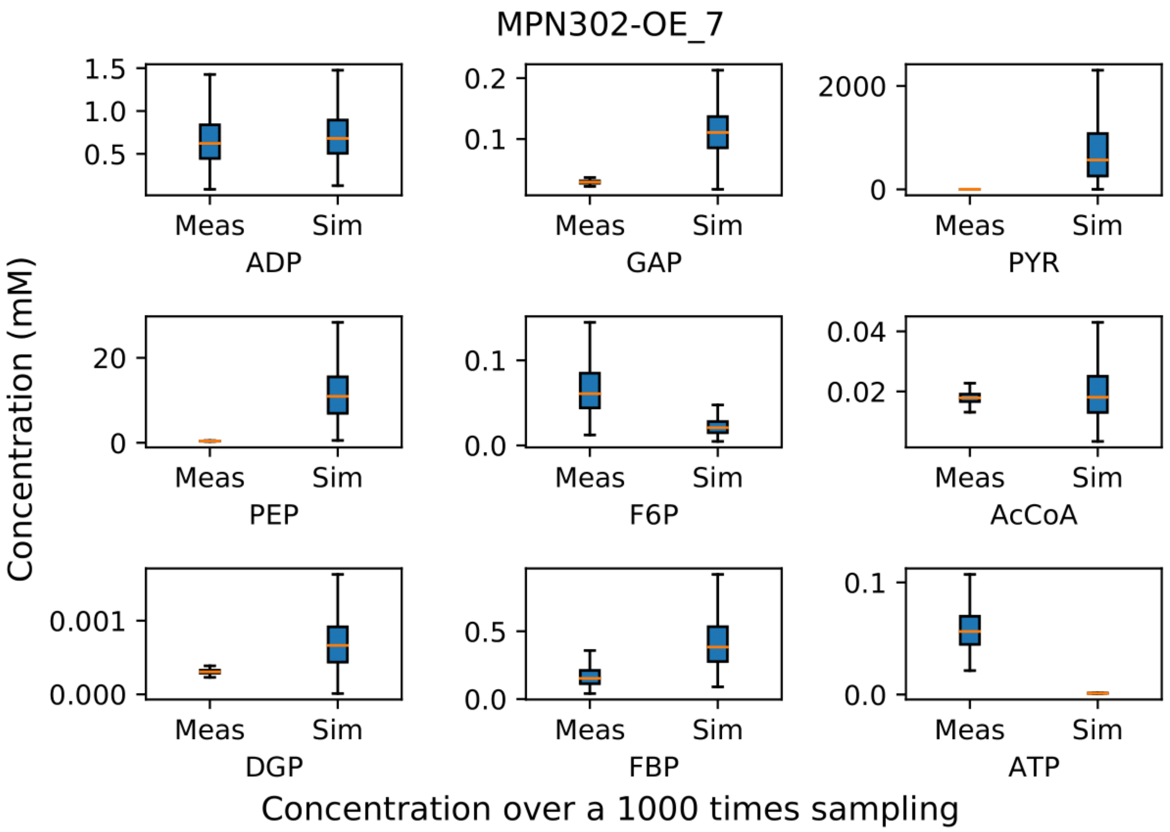
MPN302-OE_7
Concentration (mM)
ADP
GAP
PYR
PEP
F6P
AcCoA
DGP
FBP
ATP
Meas
Sim
Concentration over a 1000 times sampling

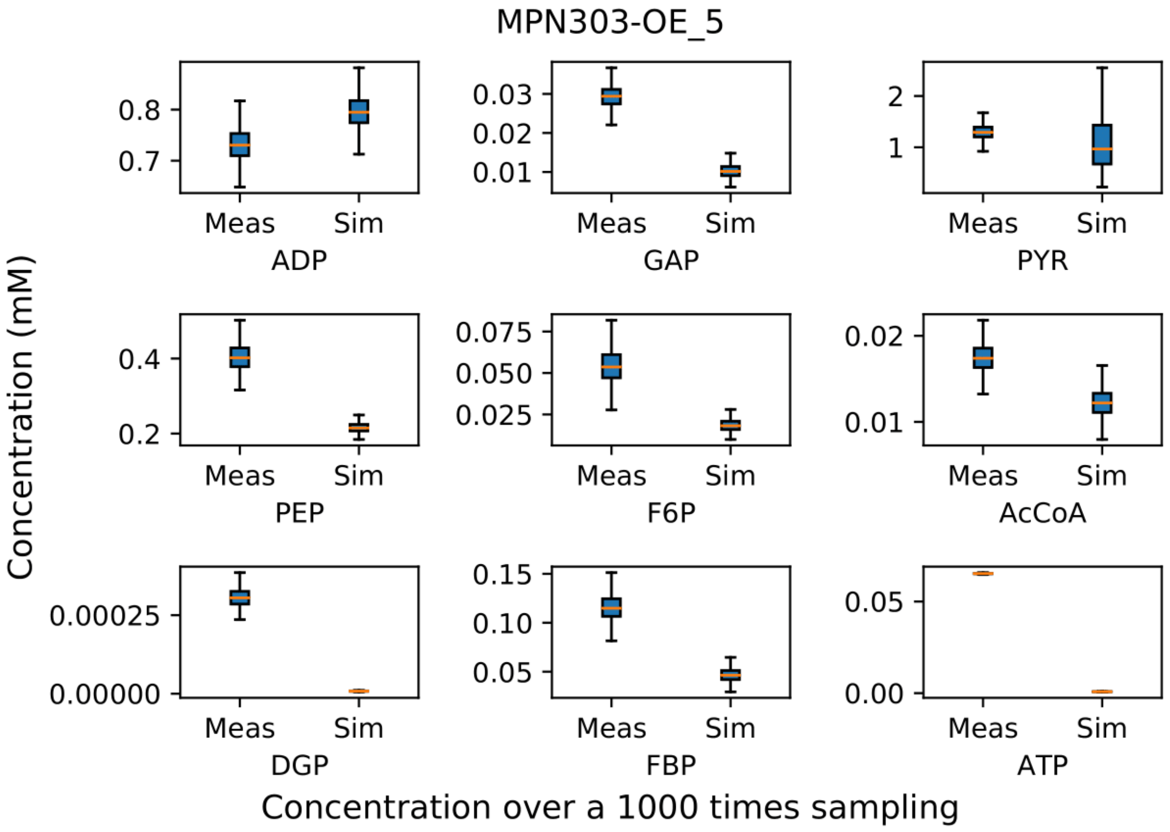
MPN303-OE_5
Concentration (mM)
ADP
GAP
PYR
PEP
F6P
AcCoA
DGP
FBP
ATP
Meas
Sim
Concentration over a 1000 times sampling

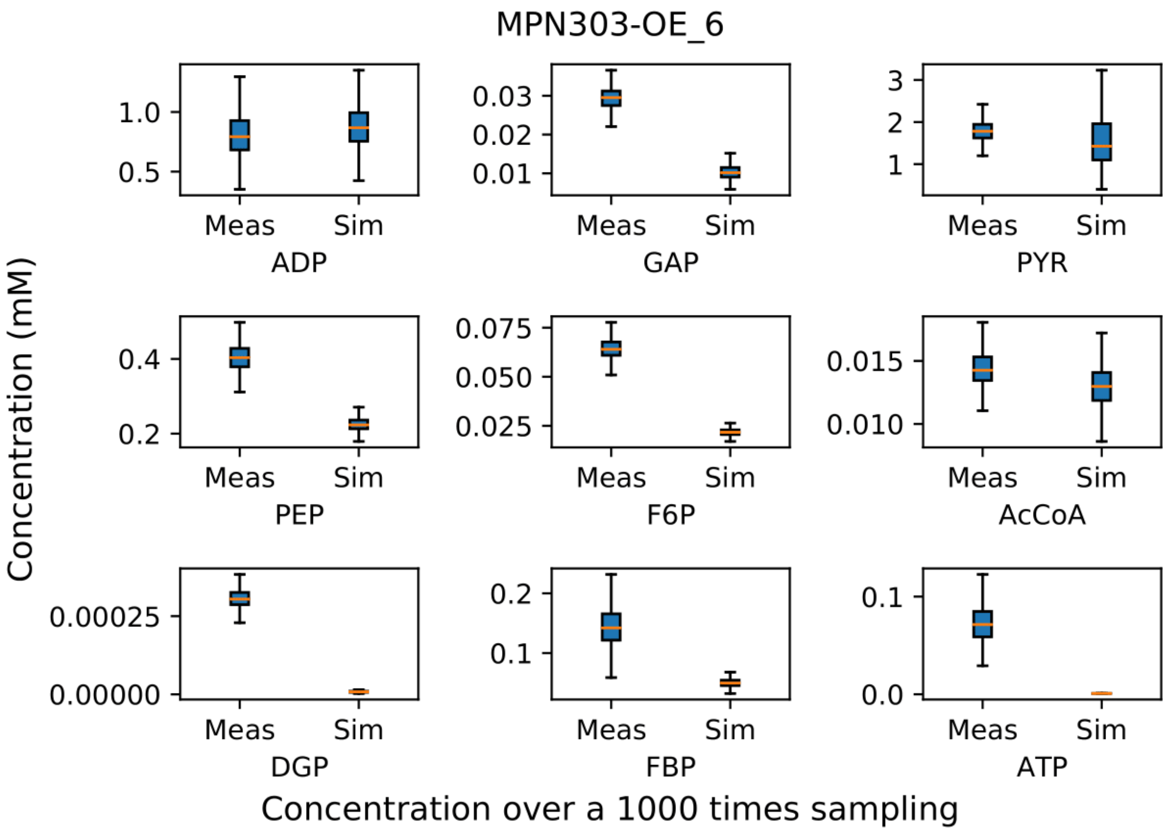
MPN303-OE_6
Concentration (mM)
1.0
0.5
Meas
Sim
ADP
0.03
0.02
0.01
GAP
3
2
1
PYR
0.4
0.2
PEP
0.075
0.050
0.025
F6P
0.015
0.010
AcCoA
0.00025
0.00000
DGP
0.2
0.1
FBP
0.1
0.0
ATP
Concentration over a 1000 times sampling

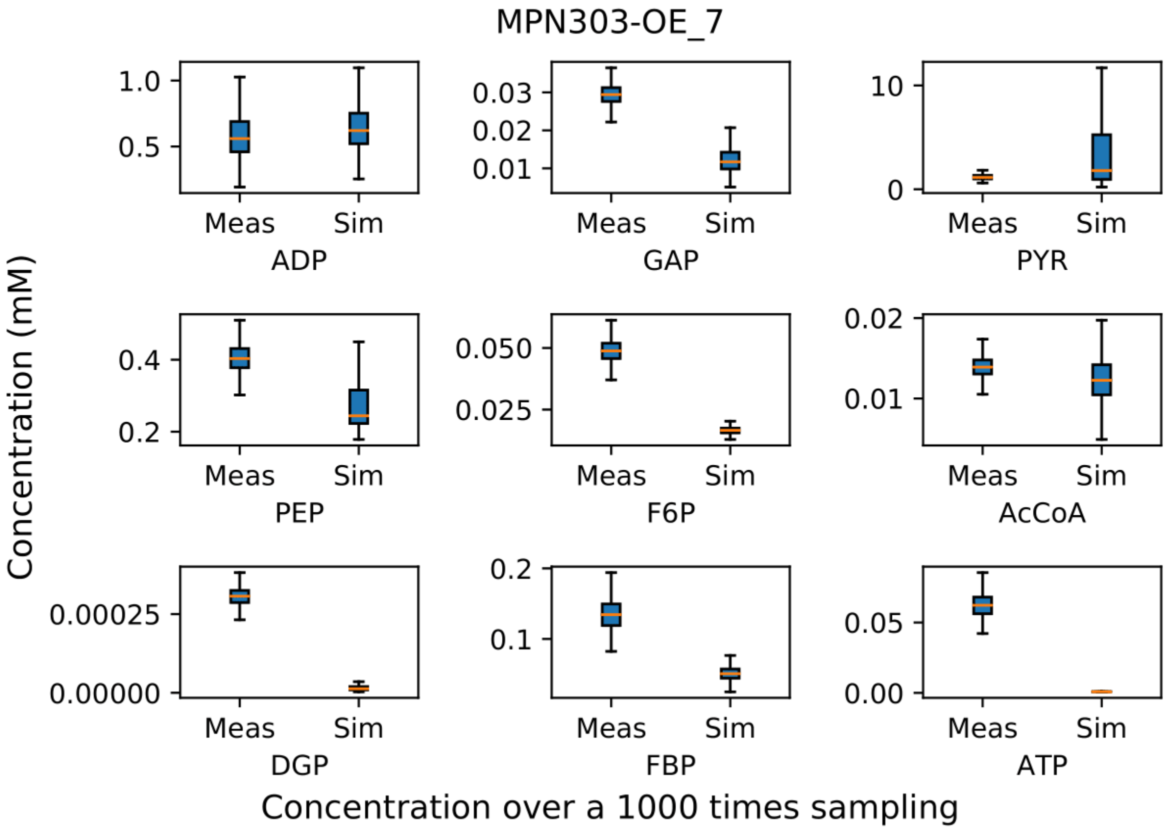
MPN303-OE_7
Concentration (mM)
1.0
0.5
Meas
Sim
ADP
0.03
0.02
0.01
GAP
10
0
PYR
0.4
0.2
PEP
0.050
0.025
F6P
0.02
0.01
AcCoA
0.00025
0.00000
DGP
0.2
0.1
FBP
0.05
0.00
ATP
Concentration over a 1000 times sampling

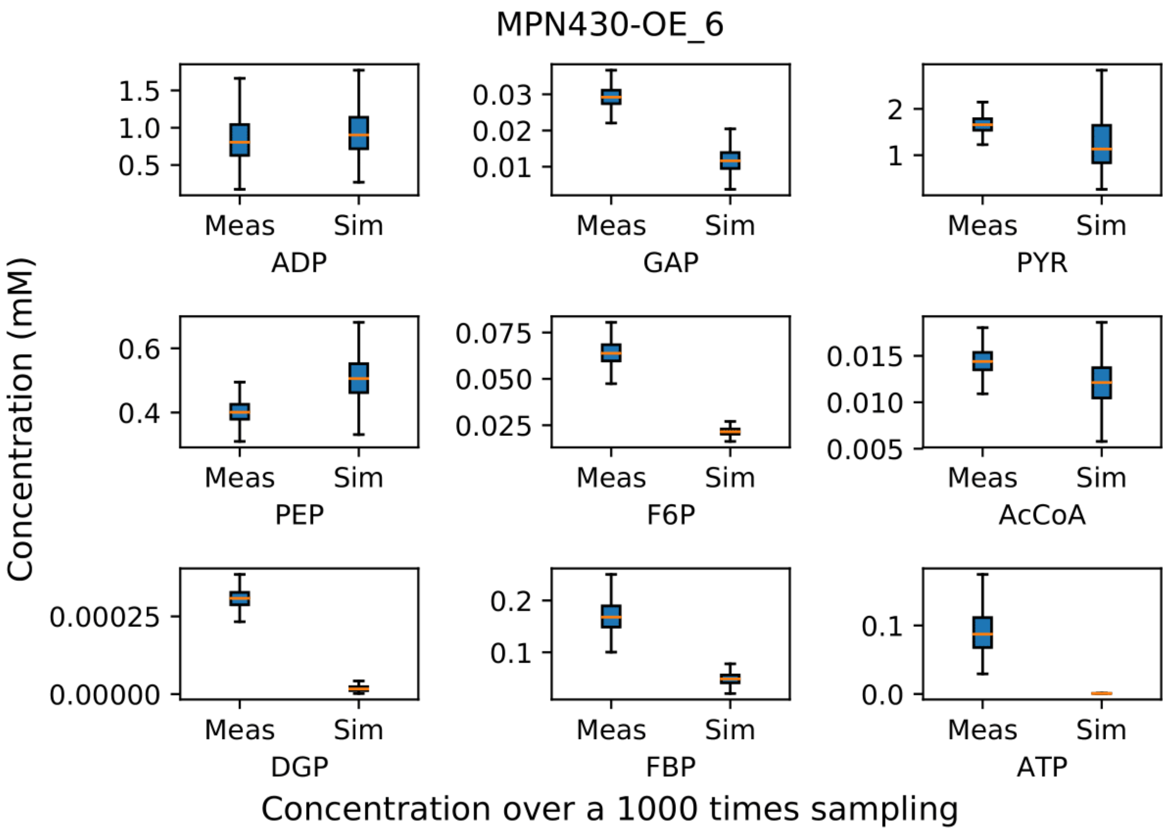
MPN430-OE_6
Concentration (mM)
ADP
GAP
PYR
PEP
F6P
AcCoA
DGP
FBP
ATP
Meas
Sim
Concentration over a 1000 times sampling

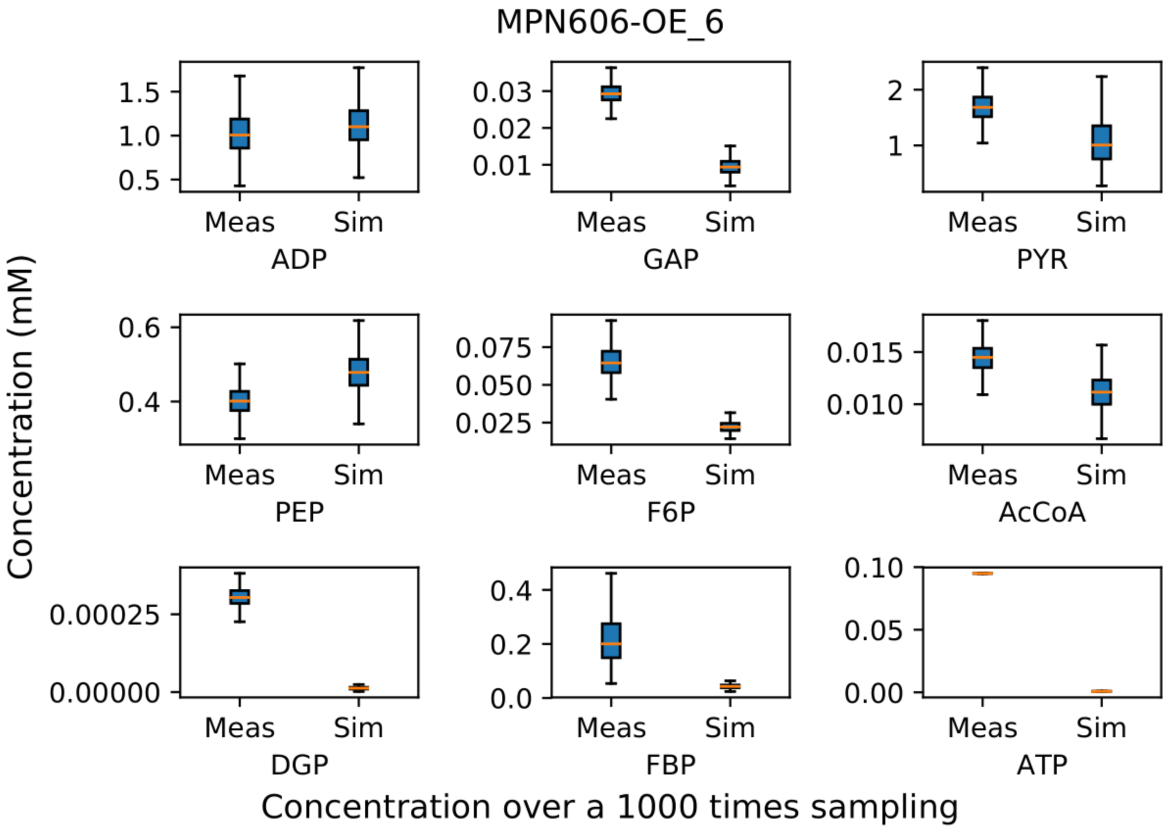
MPN606-OE_6
Concentration (mM)
ADP
GAP
PYR
PEP
F6P
AcCoA
DGP
FBP
ATP
Meas
Sim
Concentration over a 1000 times sampling

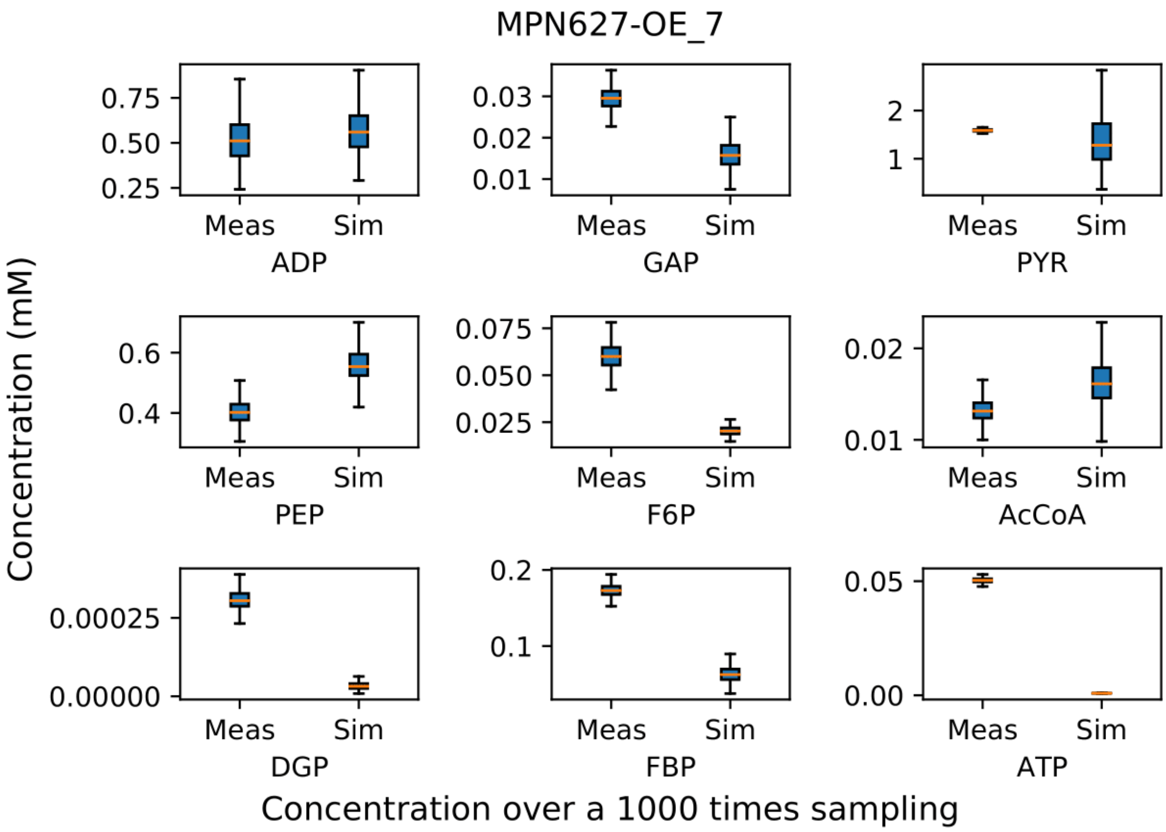
MPN627-OE_7
Concentration (mM)
Meas
Sim
ADP
GAP
PYR
PEP
F6P
AcCoA
DGP
FBP
ATP
Concentration over a 1000 times sampling

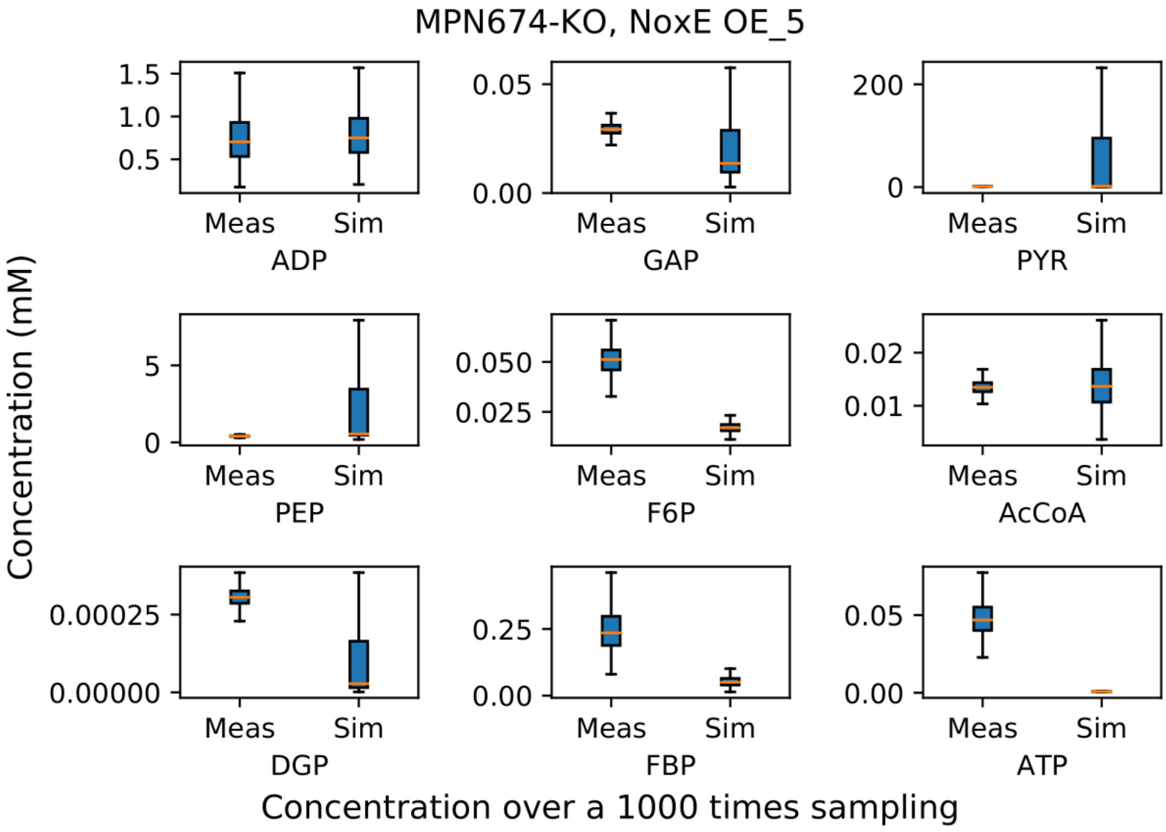
MPN674-KO, NoxE OE_5
Concentration (mM)
Meas
Sim
ADP
GAP
PYR
PEP
F6P
AcCoA
DGP
FBP
ATP
Concentration over a 1000 times sampling

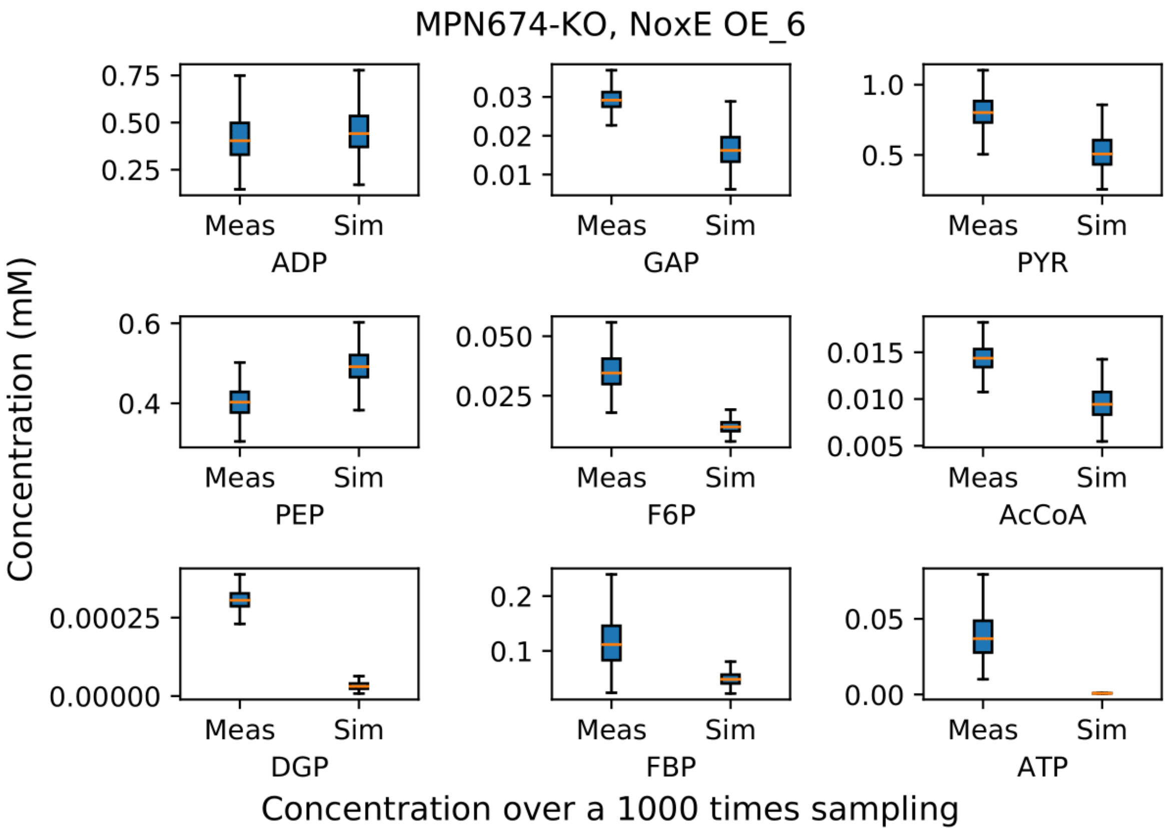
MPN674-KO, NoxE OE_6
Concentration (mM)
Meas
Sim
ADP
GAP
PYR
PEP
F6P
AcCoA
DGP
FBP
ATP
Concentration over a 1000 times sampling

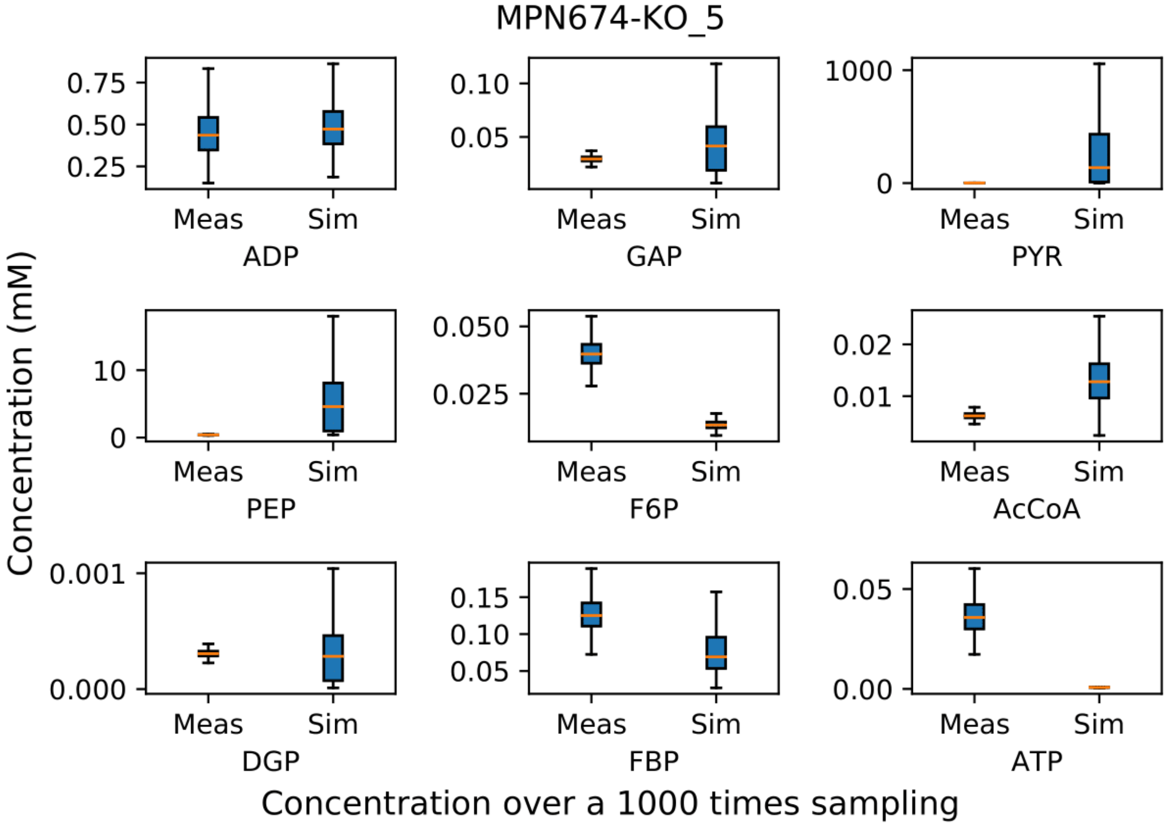
MPN674-KO_5
Concentration (mM)
Meas
Sim
ADP
GAP
PYR
PEP
F6P
AcCoA
DGP
FBP
ATP
Concentration over a 1000 times sampling

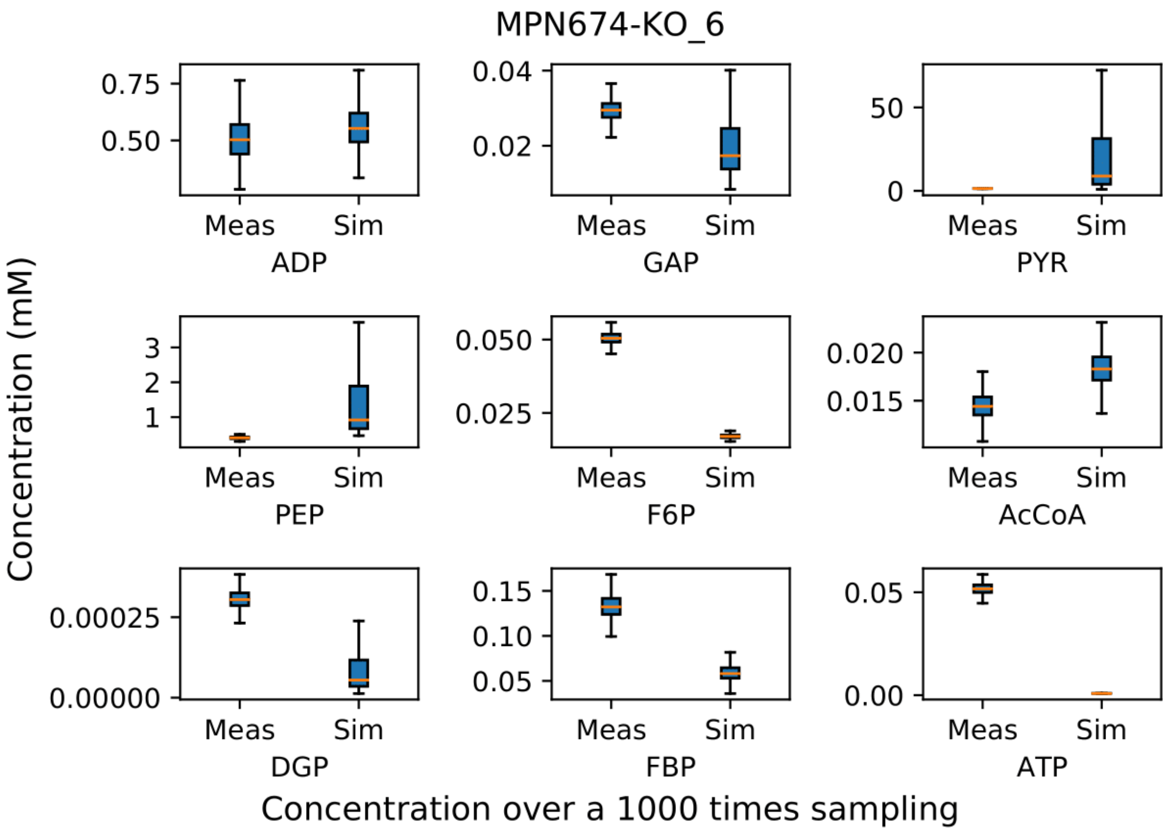
MPN674-KO_6
Concentration (mM)
0.75
0.50
Meas
Sim
ADP
0.04
0.02
Meas
Sim
GAP
50
0
Meas
Sim
PYR
3
2
1
Meas
Sim
PEP
0.050
0.025
Meas
Sim
F6P
0.020
0.015
Meas
Sim
AcCoA
0.00025
0.00000
Meas
Sim
DGP
0.15
0.10
0.05
Meas
Sim
FBP
0.05
0.00
Meas
Sim
ATP
Concentration over a 1000 times sampling

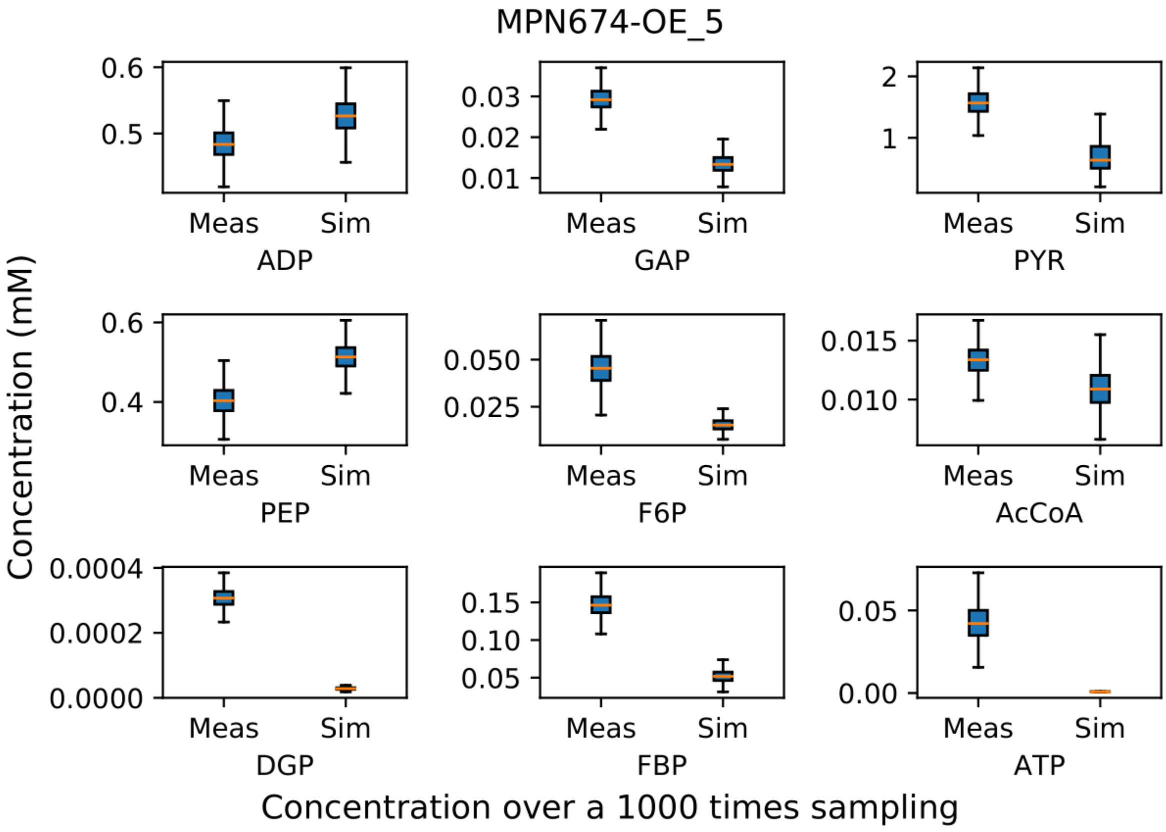
MPN674-OE_5
Concentration (mM)
0.6
0.5
Meas
Sim
ADP
0.03
0.02
0.01
Meas
Sim
GAP
2
1
Meas
Sim
PYR
0.6
0.4
Meas
Sim
PEP
0.050
0.025
Meas
Sim
F6P
0.015
0.010
Meas
Sim
AcCoA
0.0004
0.0002
0.0000
Meas
Sim
DGP
0.15
0.10
0.05
Meas
Sim
FBP
0.05
0.00
Meas
Sim
ATP
Concentration over a 1000 times sampling

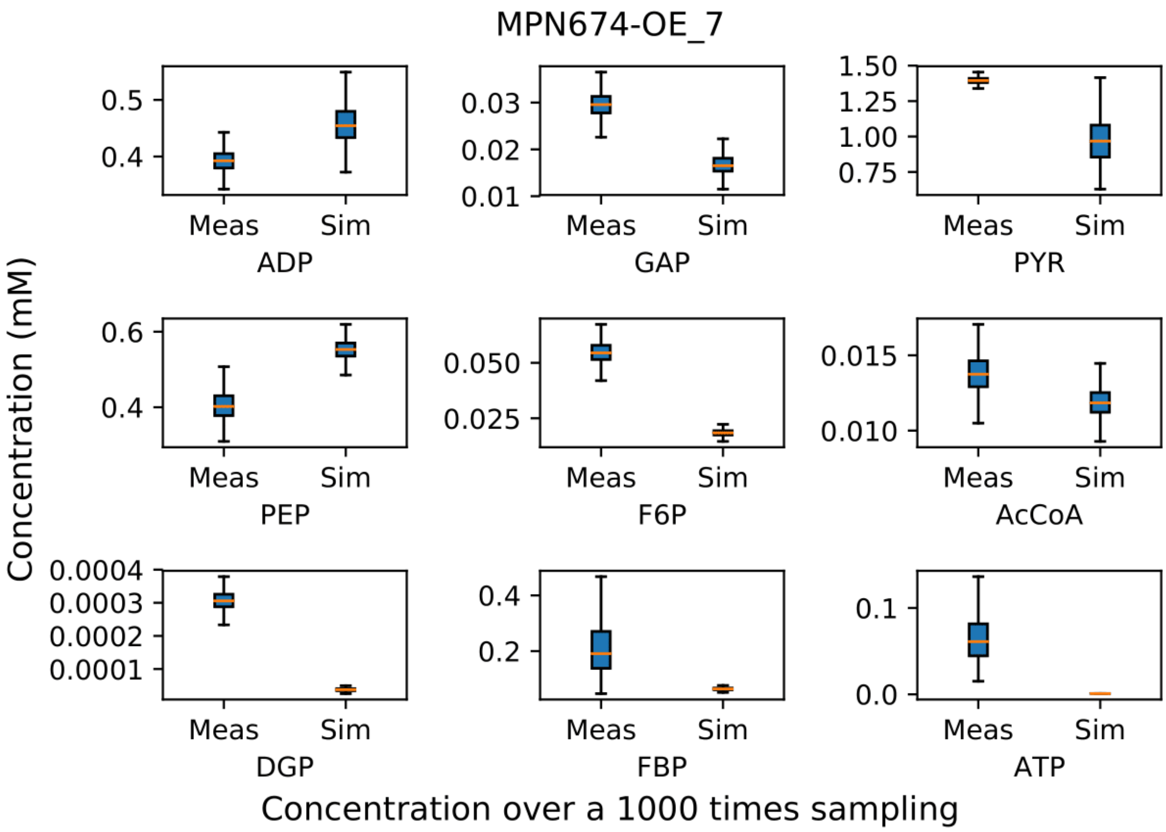
MPN674-OE_7
Concentration (mM)
ADP
GAP
PYR
PEP
F6P
AcCoA
DGP
FBP
ATP
Meas
Sim
0.5
0.4
0.03
0.02
0.01
1.50
1.25
1.00
0.75
0.6
0.4
0.050
0.025
0.015
0.010
0.0004
0.0003
0.0002
0.0001
0.4
0.2
0.1
0.0
Concentration over a 1000 times sampling

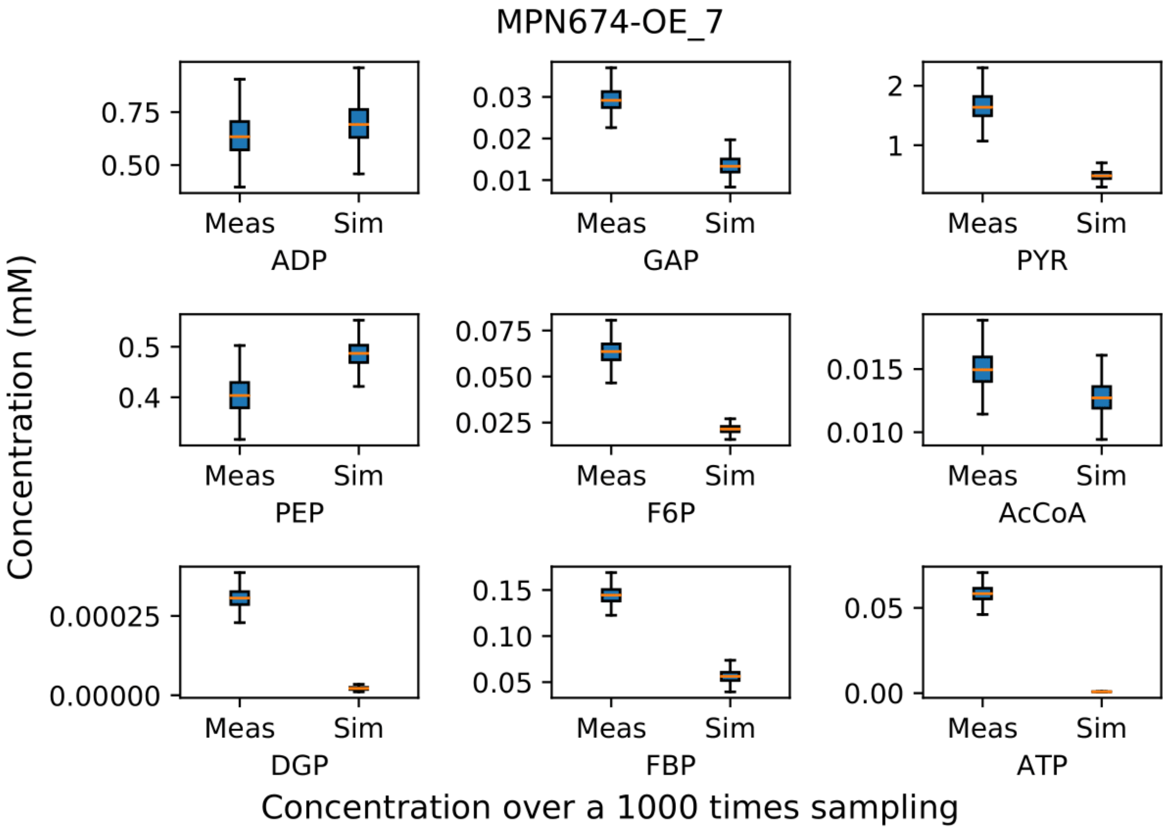
MPN674-OE_7
Concentration (mM)
ADP
GAP
PYR
PEP
F6P
AcCoA
DGP
FBP
ATP
Meas
Sim
0.75
0.50
0.03
0.02
0.01
2
1
0.5
0.4
0.075
0.050
0.025
0.015
0.010
0.00025
0.00000
0.15
0.10
0.05
0.05
0.00
Concentration over a 1000 times sampling

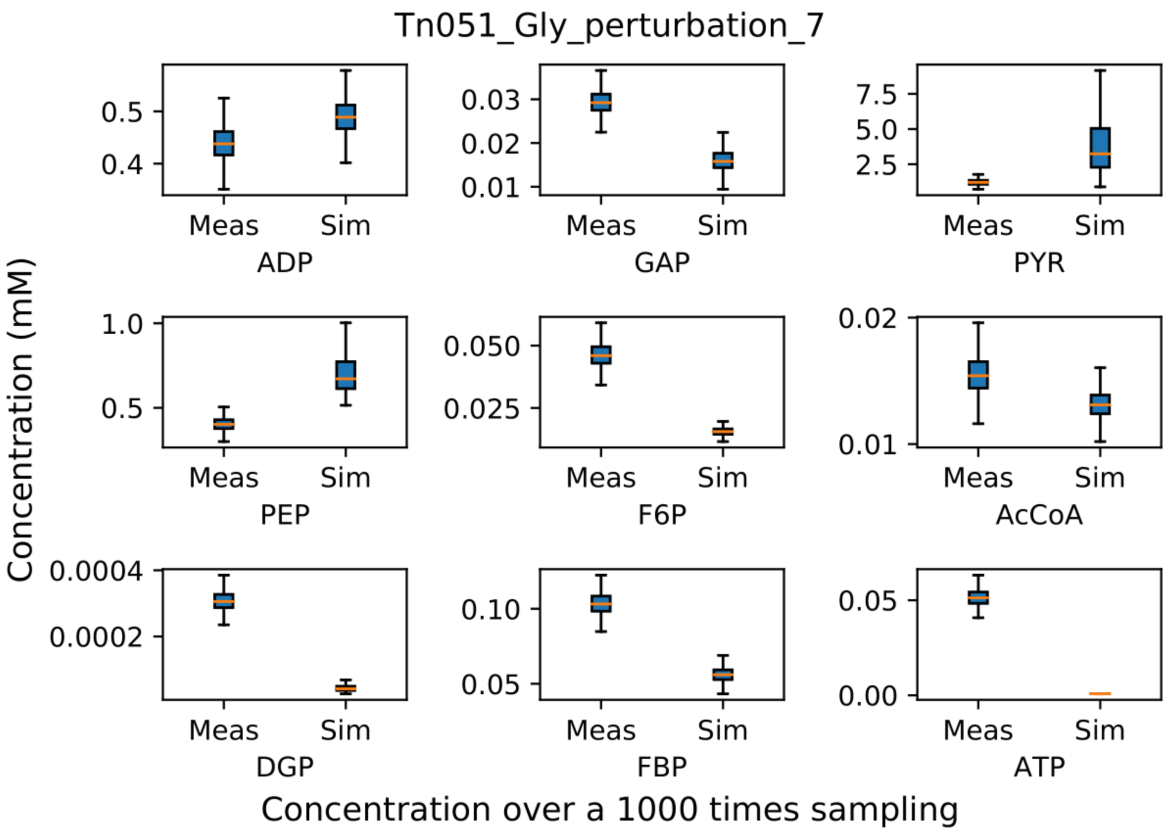
Tn051_Gly_perturbation_7
Concentration (mM)
ADP
GAP
PYR
PEP
F6P
AcCoA
DGP
FBP
ATP
Meas
Sim
Concentration over a 1000 times sampling

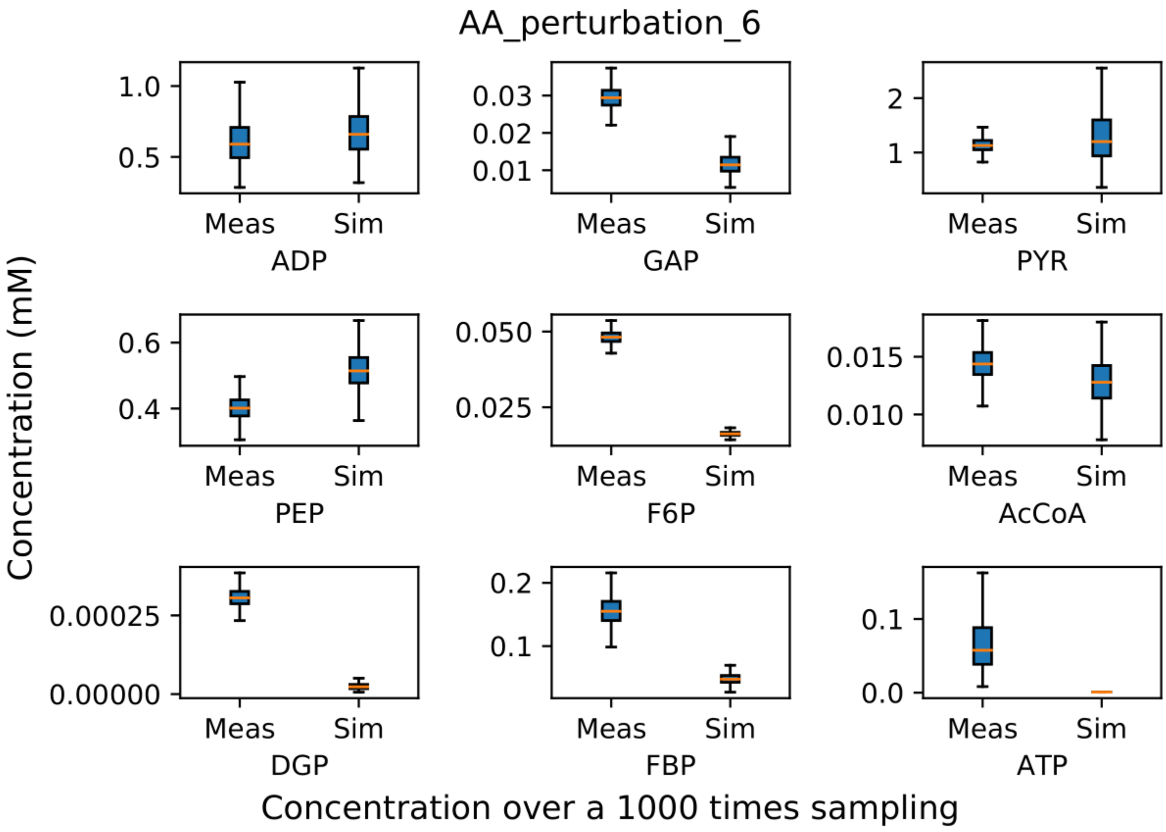
AA_perturbation_6
Concentration (mM)
ADP
GAP
PYR
PEP
F6P
AcCoA
DGP
FBP
ATP
Meas
Sim
Concentration over a 1000 times sampling

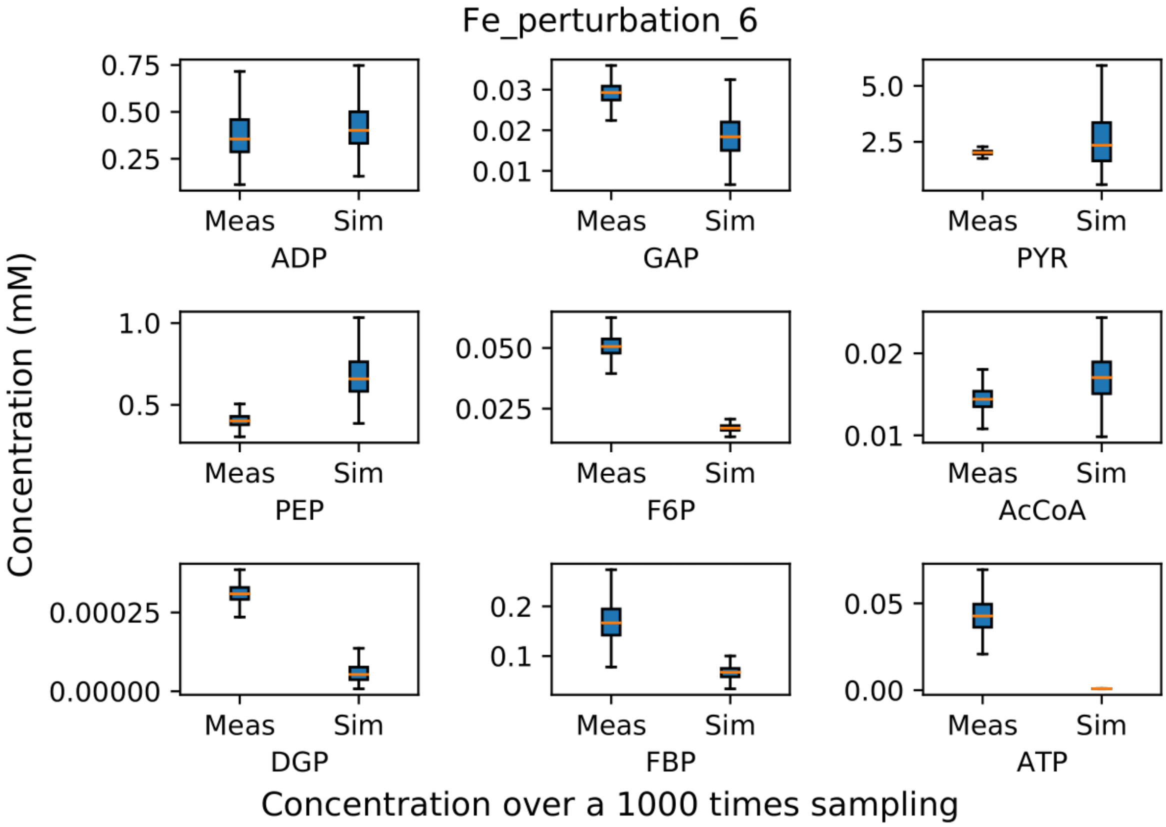
Fe_perturbation_6
Concentration (mM)
ADP
GAP
PYR
PEP
F6P
AcCoA
DGP
FBP
ATP
Meas
Sim
Concentration over a 1000 times sampling

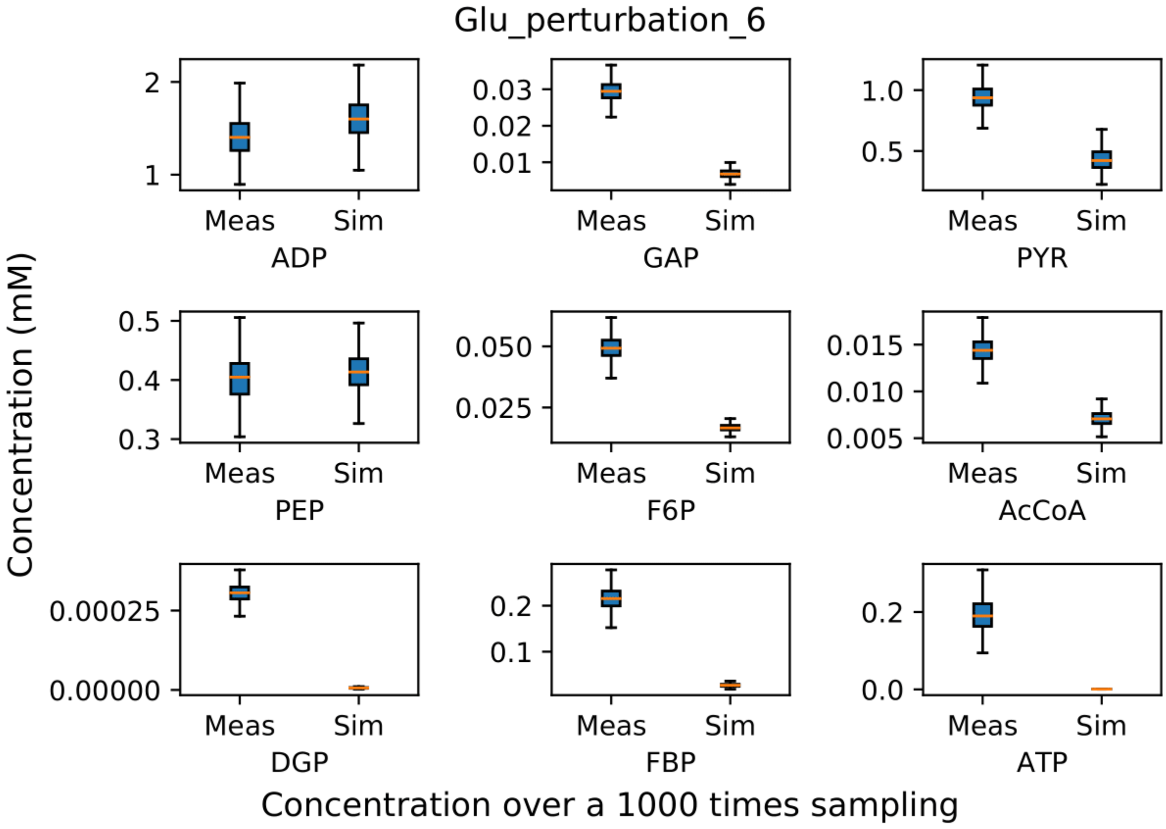
Glu_perturbation_6
Concentration (mM)
ADP
GAP
PYR
PEP
F6P
AcCoA
DGP
FBP
ATP
Meas
Sim
Concentration over a 1000 times sampling

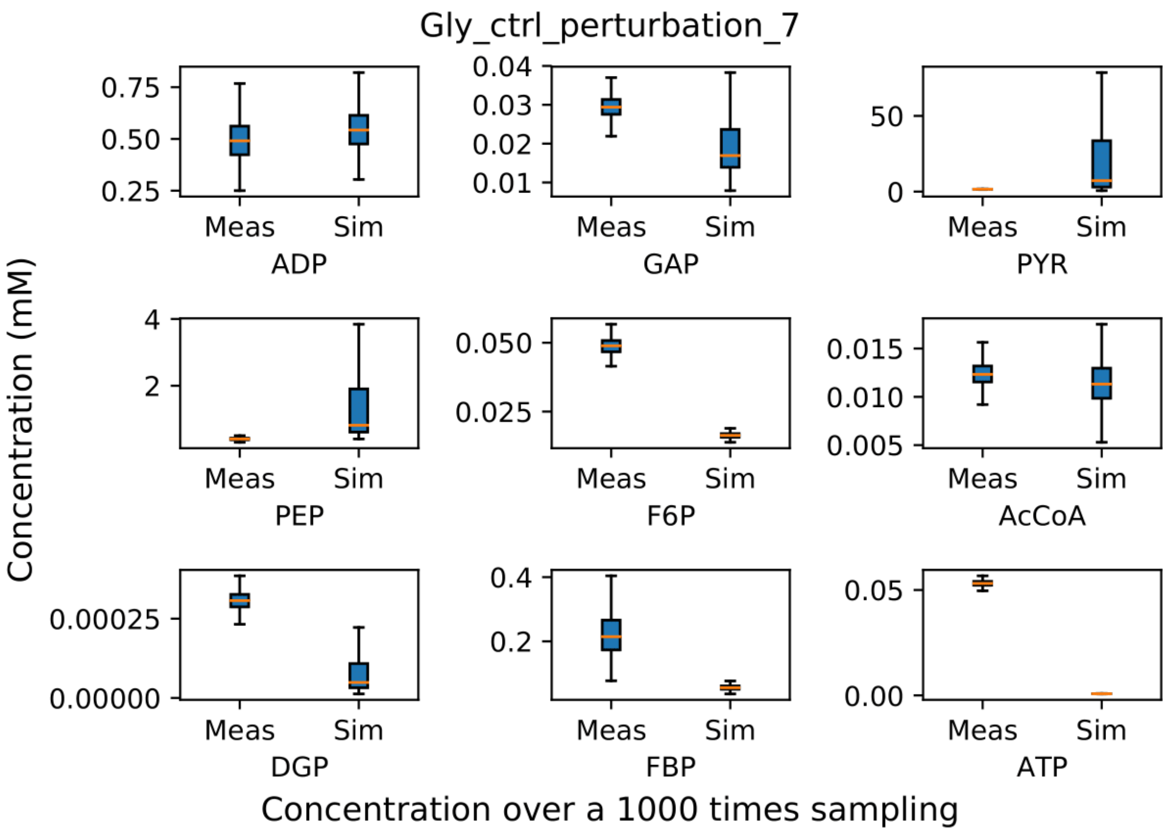
Gly_ctrl_perturbation_7
Concentration (mM)
ADP
GAP
PYR
PEP
F6P
AcCoA
DGP
FBP
ATP
Meas
Sim
Concentration over a 1000 times sampling

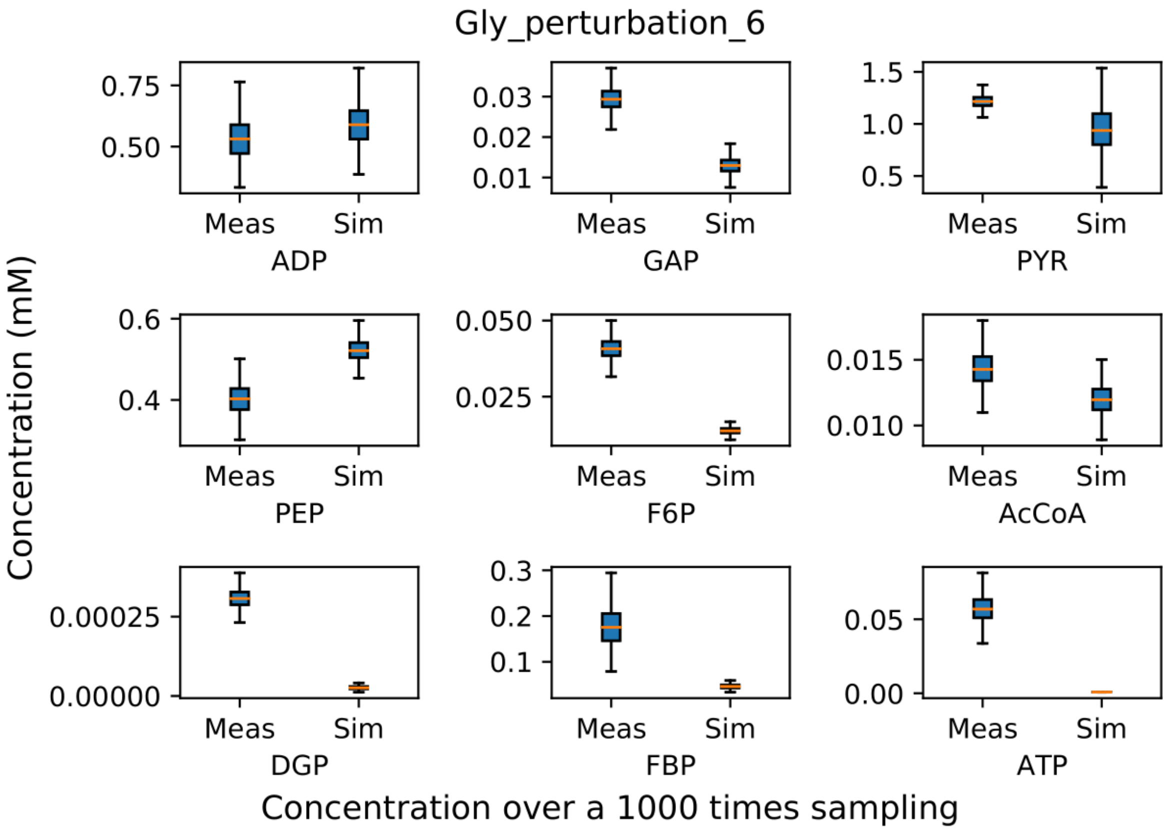
Gly_perturbation_6
Concentration (mM)
ADP
GAP
PYR
PEP
F6P
AcCoA
DGP
FBP
ATP
Meas
Sim
Concentration over a 1000 times sampling

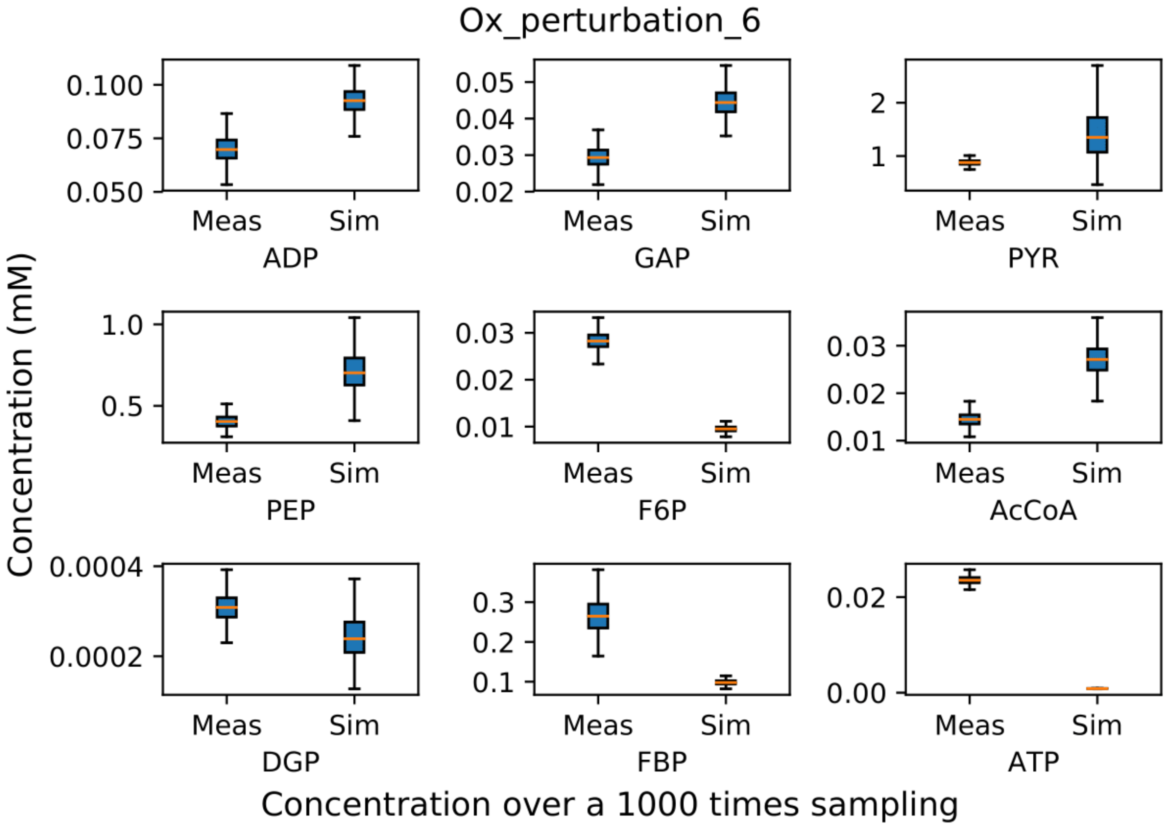
Ox_perturbation_6
Concentration (mM)
0.100
0.075
0.050
Meas
Sim
ADP
0.05
0.04
0.03
0.02
Meas
Sim
GAP
2
1
Meas
Sim
PYR
1.0
0.5
Meas
Sim
PEP
0.03
0.02
0.01
Meas
Sim
F6P
0.03
0.02
0.01
Meas
Sim
AcCoA
0.0004
0.0002
Meas
Sim
DGP
0.3
0.2
0.1
Meas
Sim
FBP
0.02
0.00
Meas
Sim
ATP
Concentration over a 1000 times sampling

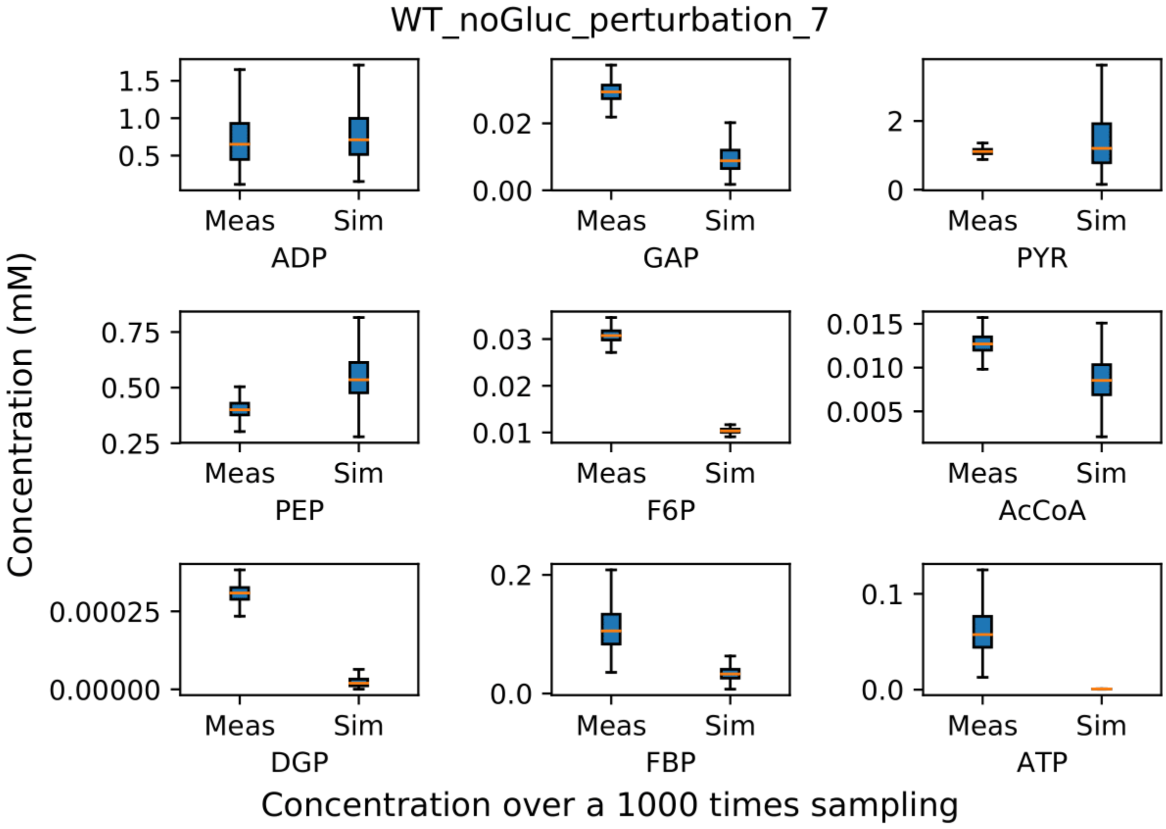
WT_noGluc_perturbation_7
Concentration (mM)
1.5
1.0
0.5
Meas
Sim
ADP
0.02
0.00
Meas
Sim
GAP
2
0
Meas
Sim
PYR
0.75
0.50
0.25
Meas
Sim
PEP
0.03
0.02
0.01
Meas
Sim
F6P
0.015
0.010
0.005
Meas
Sim
AcCoA
0.00025
0.00000
Meas
Sim
DGP
0.2
0.0
Meas
Sim
FBP
0.1
0.0
Meas
Sim
ATP
Concentration over a 1000 times sampling

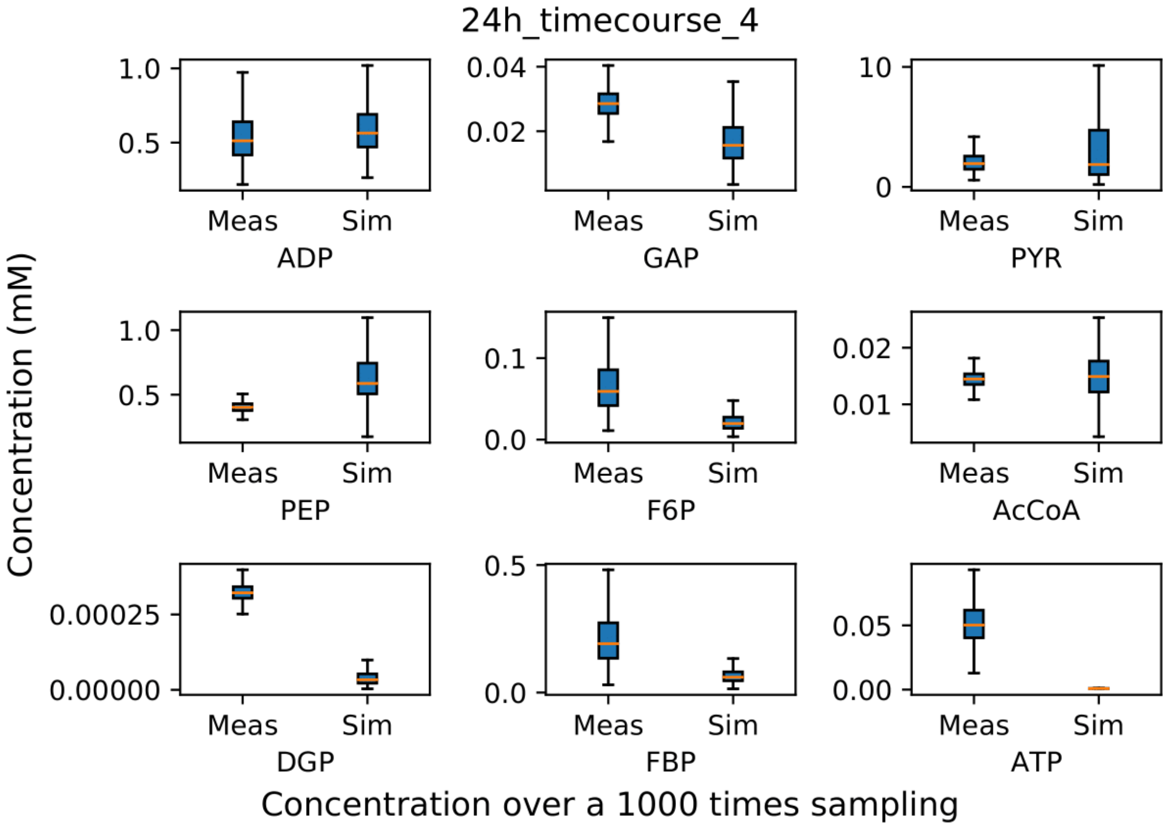

24h_timecourse_4
Concentration (mM)
Meas
Sim
ADP
GAP
PYR
PEP
F6P
AcCoA
DGP
FBP
ATP
Concentration over a 1000 times sampling


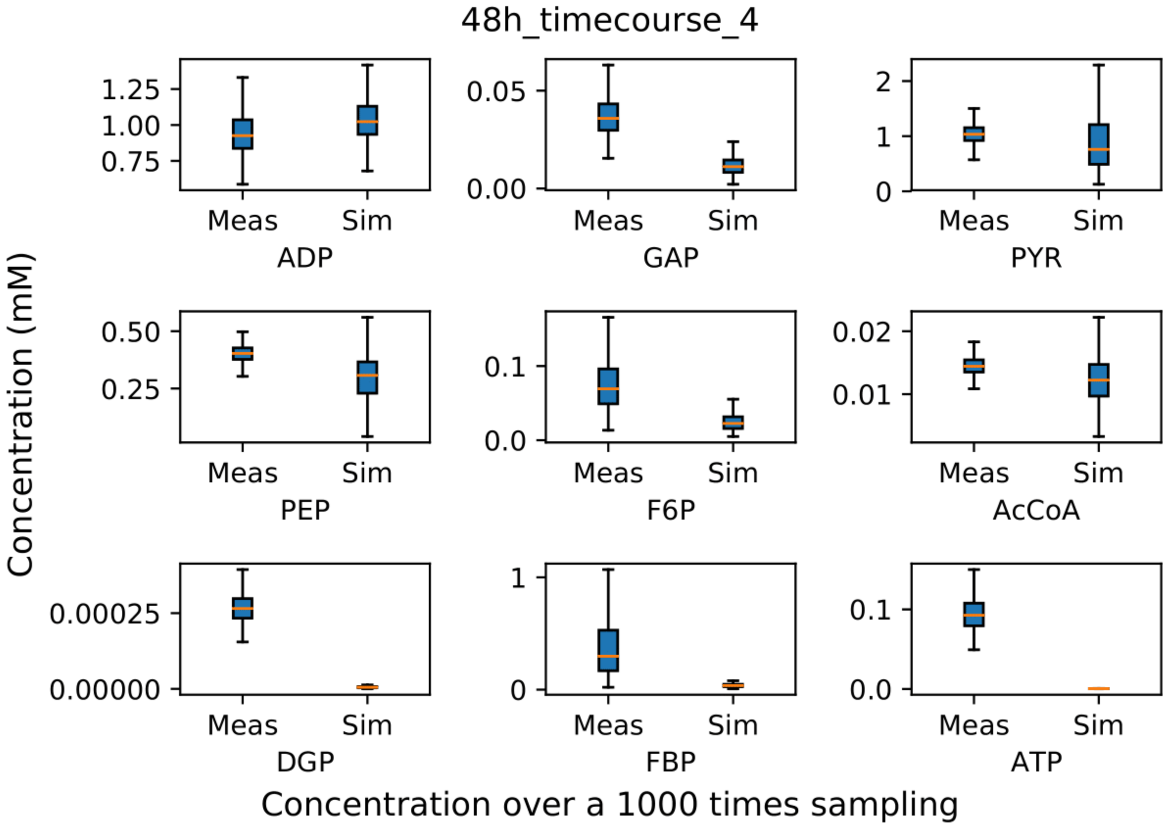

48h_timecourse_4
Concentration (mM)
Meas
Sim
ADP
GAP
PYR
PEP
F6P
AcCoA
DGP
FBP
ATP
Concentration over a 1000 times sampling

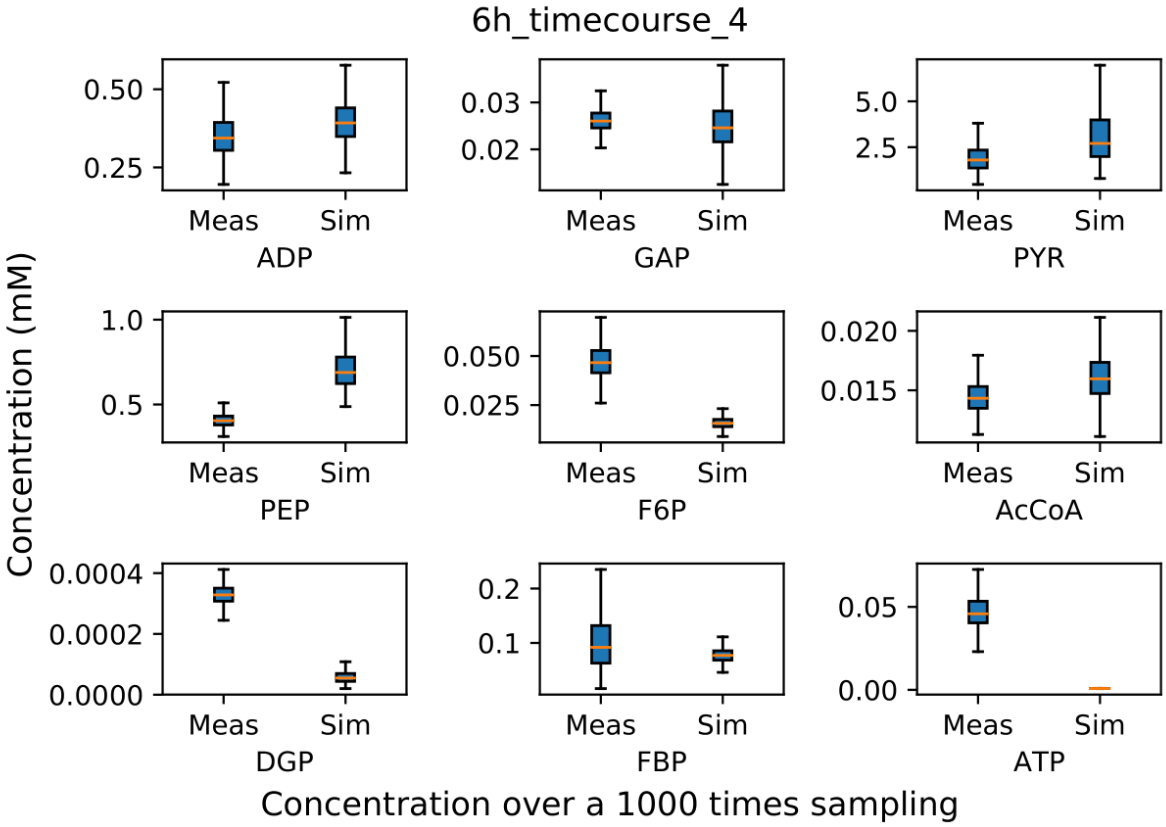
6h_timecourse_4
Concentration (mM)
ADP
GAP
PYR
PEP
F6P
AcCoA
DGP
FBP
ATP
Meas
Sim
Concentration over a 1000 times sampling

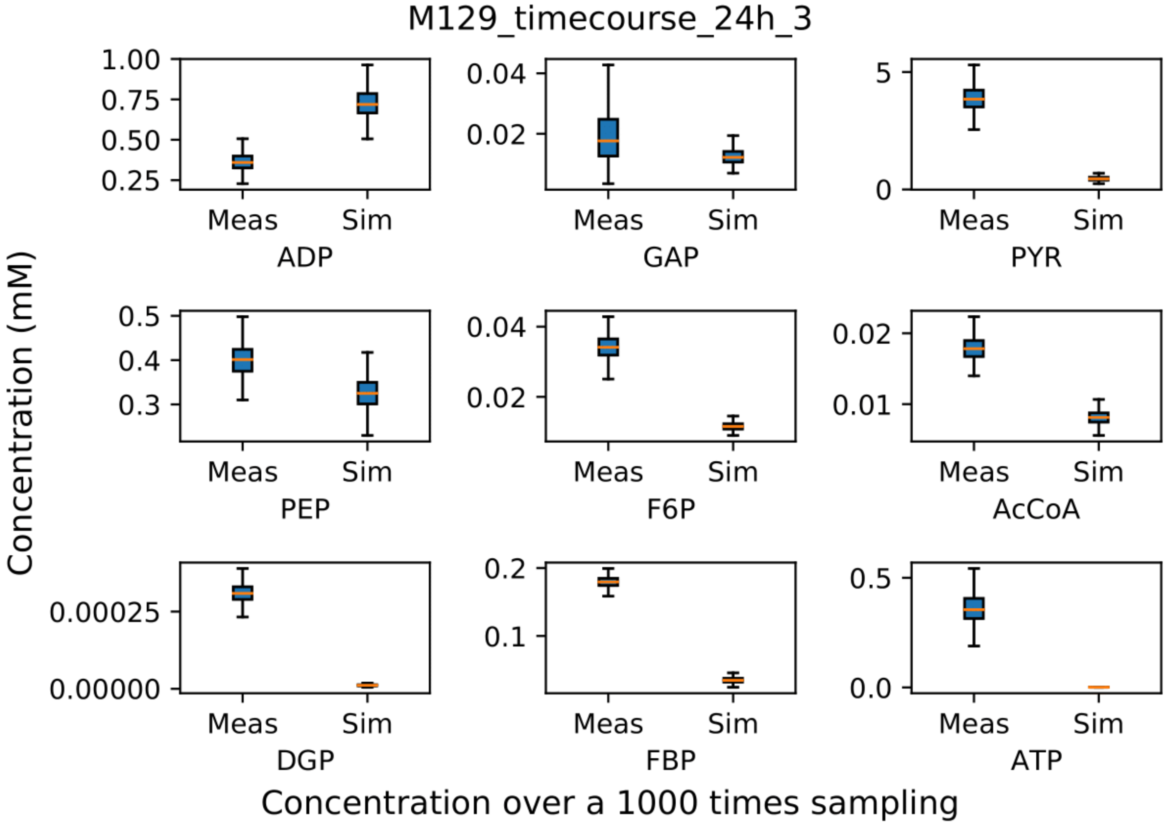
M129_timecourse_24h_3
Concentration (mM)
ADP
GAP
PYR
PEP
F6P
AcCoA
DGP
FBP
ATP
Meas
Sim
Concentration over a 1000 times sampling

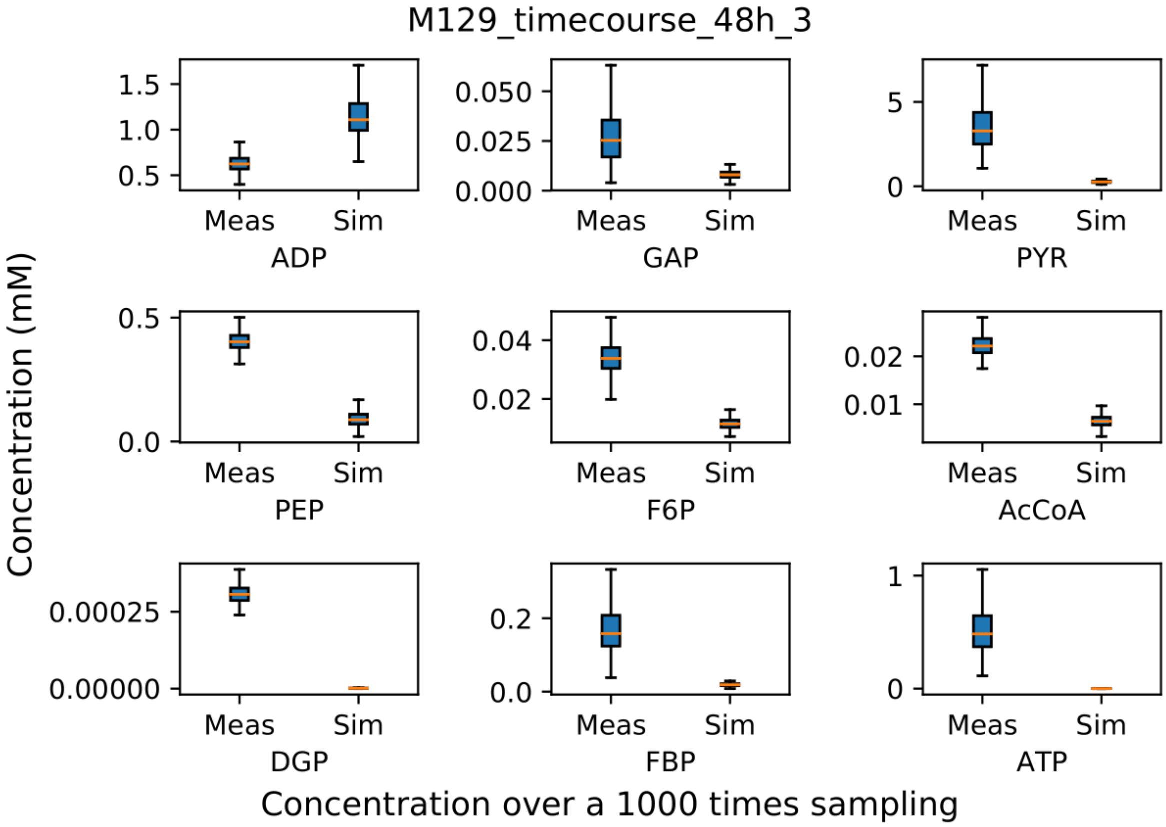
M129_timecourse_48h_3
Concentration (mM)
ADP
GAP
PYR
PEP
F6P
AcCoA
DGP
FBP
ATP
Meas
Sim
Concentration over a 1000 times sampling

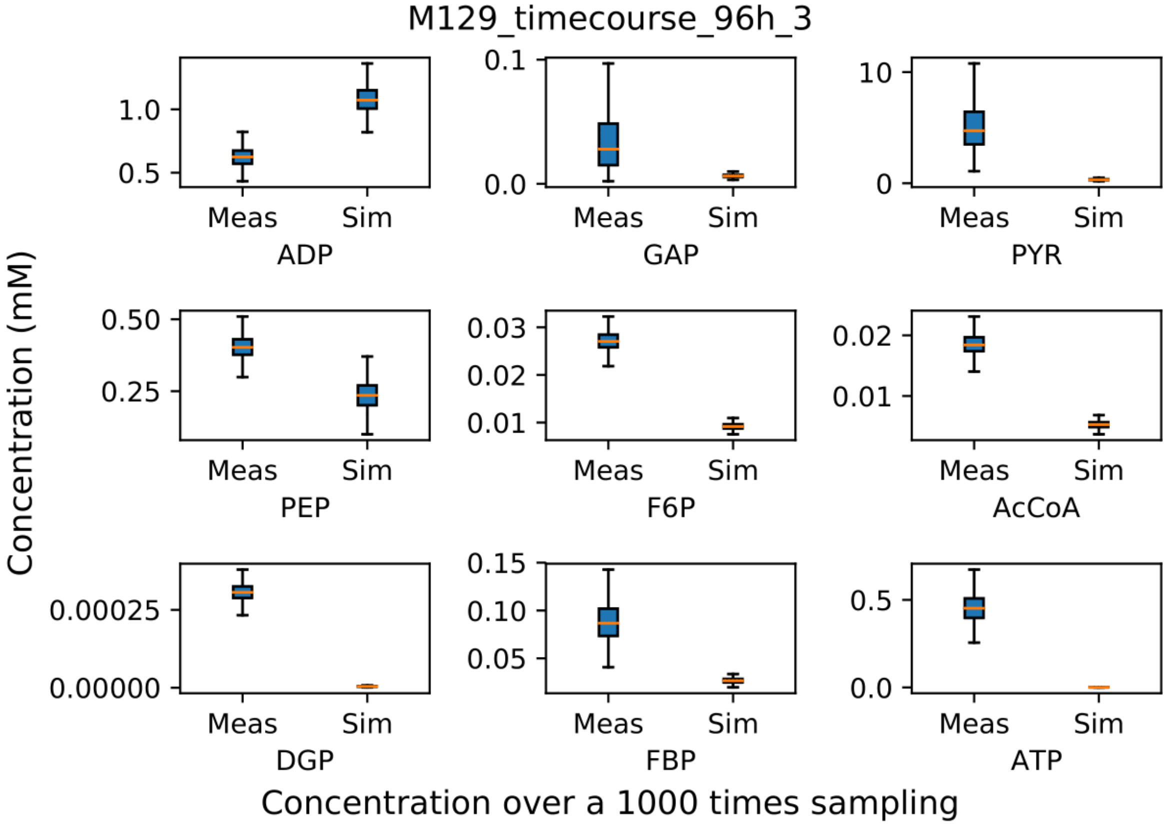
M129_timecourse_96h_3
Concentration (mM)
ADP
GAP
PYR
PEP
F6P
AcCoA
DGP
FBP
ATP
Meas
Sim
Concentration over a 1000 times sampling

### Supplementary file 1 D Metabolomics analysis.

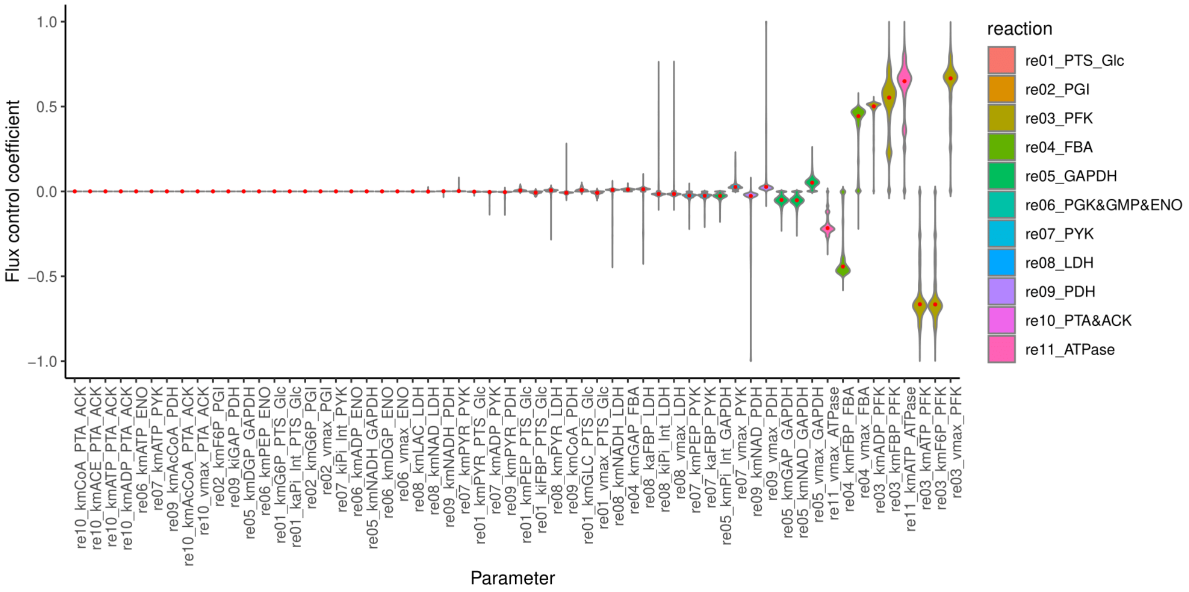


*Figure 3 Local sensitivity analysis using 1000x sampling from the measurement distribution.*

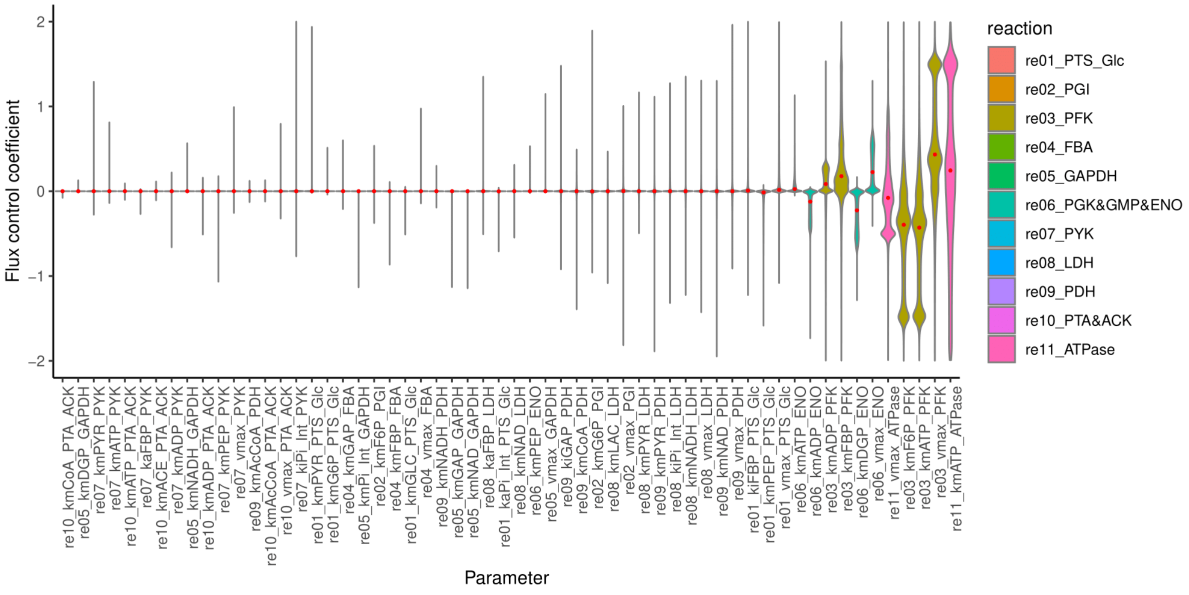


*Figure 4 Global sensitivity analysis using a 100,000 Latin Hypercube sampling withing the parameter search range.*

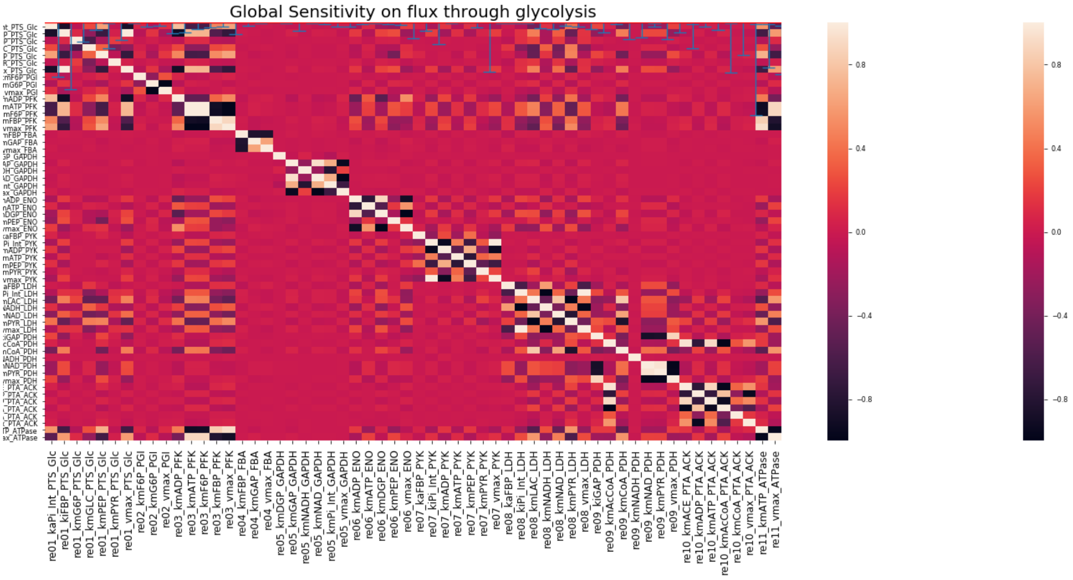


*Figure 5 Correlation analysis of control coefficients control over flux through glycolysis using a 100,000 Latin Hypercube sampling.*
